# Discovery of a first-in-class inhibitor of the PRMT5-substrate adaptor interaction

**DOI:** 10.1101/2021.02.03.429644

**Authors:** David C McKinney, Brian J McMillan, Matthew Ranaghan, Jamie A Moroco, Merissa Brousseau, Zachary Mullin-Bernstein, Meghan O’Keefe, Patrick McCarren, Michael F. Mesleh, Kathleen M. Mulvaney, Ritu Singh, Besnik Bajrami, Adam Skepner, David E. Timm, Dale Porter, Virendar K. Kaushik, William R. Sellers, Alessandra Ianari

## Abstract

PRMT5 and its substrate adaptor proteins (SAPs), pICln and Riok1, are synthetic lethal dependencies in MTAP-deleted cancer cells. SAPs share a conserved PRMT5 binding motif (PBM) which mediates binding to a surface of PRMT5 distal to the catalytic site. This interaction is required for methylation of several PRMT5 substrates, including histone and spliceosome complexes. We screened for small molecule inhibitors of the PRMT5-PBM interaction and validated a compound series which binds to the PRMT5-PBM interface and directly inhibits binding of SAPs. Mode of action and structure determination studies revealed that these compounds form a covalent bond between a halogenated pyridazinone group and cysteine 278 of PRMT5. Optimization of the starting hit produced a lead compound, BRD0639, which engages the target in cells, disrupts the PRMT5-RIOK1 complex, and reduces substrate methylation. BRD0639 is a first-in-class PBM-competitive small molecule that can support studies of PBM-dependent PRMT5 activities and the development of novel PRMT5 inhibitors that selectively target these functions.

## Introduction

Recent advances in cancer genomics and molecular biology have led to the identification of novel tumor selective therapeutic opportunities. For example, 15-50% of pancreatic cancers, glioblastoma and mesotheliomas, cancers for which there is a critical need for new drugs, bear deletions of the methylthioadenosine phosphorylase gene, MTAP. The MTAP gene is in close proximity to the tumor suppressor CDKN2A locus on chromosome 9, and is hence frequently codeleted. While loss of MTAP has as yet unknown functional consequences for tumorigenesis, it leads to accumulation of its substrate methylthioadenosine (MTA), which acts as an endogenous competitive inhibitor of S-adenosylmethionine (SAM), the methyl donor used by the protein arginine N-methyltransferase, PRMT5, which is involved in the regulation of gene expression, mRNA splicing, protein translation, DNA damage response and immune functions. PRMT5 has an obligate partner, WDR77, with which it forms a hetero-octamer, also known as the methylosome complex. PRMT5 belongs to the PRMT family, which consists of nine members, all of which use SAM as their methyl donor cofactor. However, MTA appears to have selective inhibitory activity on PRMT5 alone, likely due to key structural features within its SAM binding pocket ^*1, 2*^. This partial inhibition of PRMT5 sensitizes MTAP-deleted cells to further loss of PRMT5 function by siRNA. This observed synthetic lethal phenotype has led to the hypothesis that pharmacological PRMT5 inhibition could be a viable strategy with a suitable therapeutic window for clinical use against MTAP-deleted cancers ^*3, 1, 2*^.

Several potent and selective inhibitors of the PRMT5 catalytic pocket have been developed. However, these molecules act with either a SAM cooperative ^*4*^ or SAM/MTA-competitive mode of action ^*5, 6, 7*^ and thus do not appear to take advantage of MTAP deletion to provide the desired therapeutic window. An MTA-cooperative compound might potentially leverage this synthetic lethality, however such an inhibitor is not yet available ^*8*^. In addition, inhibitors of the upstream enzyme MAT2A, which catalyzes the synthesis of SAM, have been generated to exploit the unbalanced SAM/MTA ratio of MTAP-deleted cancers. Preclinical data support an MTAP-dependent and synergistic anti-tumor activity when used in combination with taxanes or gemcitabine ^*9*^. However, SAM is a ubiquitous methyl donor, and a reduction of its levels has an effect on multiple methyltransferases, potentially posing a toxicity risk to this approach. The efficacy and safety profile of MAT2A inhibitors are currently being evaluated in clinical trials (*<u>NCT03435250</u>*).

Large-scale shRNA screens have also implicated PRMT5 SAPs, pICln and RIOK1, as a potential alternative route to develop MTAP-null selective therapeutics ^*10*^. This mechanism would be distinct from catalytic site inhibition and potentially provide synergy with, or a different therapeutic profile than, the previously mentioned approaches. We recently elucidated the molecular mechanism mediating the interaction between PRMT5 and its SAPs. Specifically, we identified a conserved 7 residue peptide sequence, GQF(D/E)DA(D/E), present in all three known PRMT5 SAPs (pICln, RIOK1 and COPR5), that mediates substrate adaptor binding to PRMT5, which we termed the PRMT5 Binding Motif (PBM). We solved the structure of a PBM peptide bound to its cognate site on PRMT5, which is distinct and distal from the catalytic domain. We also determined that genetic perturbation of this site results in loss of substrate methylation and a reduction in MTAP-deleted cell growth relative to WT ^*11*^. Based on these observations, we launched a screening campaign to identify small molecules capable of inhibiting this interaction.

Here we describe the identification and development of the first-in-class PBM competitive covalent compound, BRD0639, a chemical probe with on-target cellular activity that can support the exploration of a new class of PRMT5 inhibitors.

## Results

### Hit finding activities

To identify inhibitors of the PBM peptide interaction we utilized several hit-finding approaches to screen lead-like or fragment compound libraries (Fig. 1a). As a primary strategy, we utilized a competition fluorescence polarization (FP) assay, to measure the interaction between a fluorophore-labeled RIOK1 PBM peptide and the purified PRMT5:WDR77 heterooctameric complex. In total, we screened more than 900K small molecules: 850K from the Diversity and Lead-like compound libraries from Charles River Laboratories, 50K from the Chembridge DIVERSet, 14K compounds from the Broad Institute Diversity Oriented Synthesis (DOS) library ^*12*^, and 1K compounds selected by virtual screening. We also carried out an NMR-based fragment screen and an *in silico* virtual pharmacophore screen (Fig. 1a and SI). As these approaches did not yield starting points for optimization, we focused on the FP identified screening hits.

**Figure 1.**
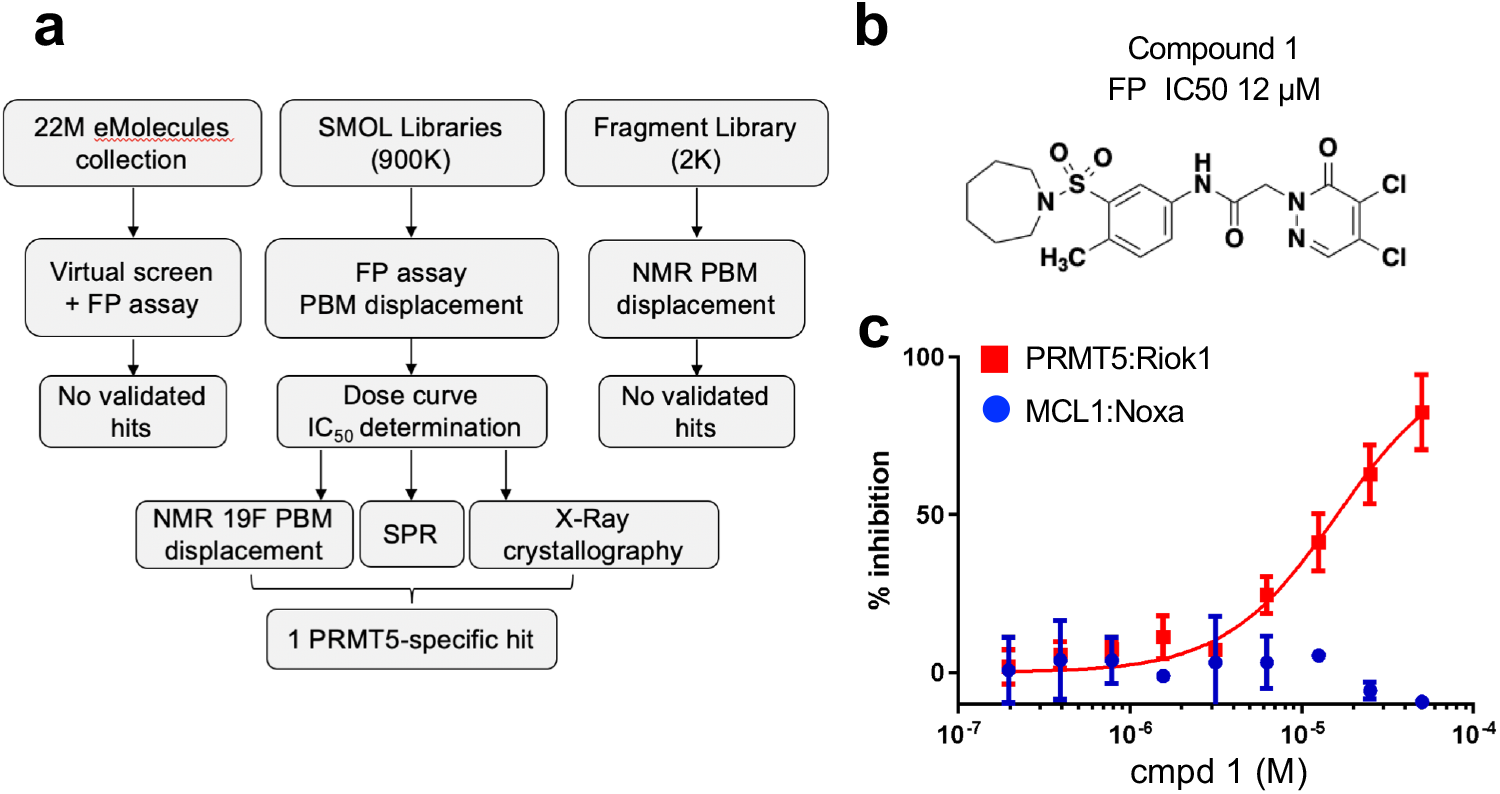
Hit Finding Activities. **(A) Left:** PBM peptide from Riok1 (gold) bound to PRMT5 (green). Residues involved in covalent compound binding are labeled. **Right:** Cartoon representation of the PRMT5(green):WDR77(grey) protomer structure. The PBM peptide is represented in gold and SAM in magenta **(B)** Hit discovery flow chart for inhibitors of the PBM:PRMT5 interaction. **(C)** Chemical structure of compound **1**. **(D)** Dose-dependent displacement of a fluorescently-labeled RioK1-derived PBM peptide probe by compound **1** as measured by FP.

The initial FP hits (>20% inhibition) were reassessed in duplicate using the PRMT5 FP system and counter-screened against an FP assay using an unrelated protein complex (MCL1 receptor and Noxa peptide) to remove potential non-specific compounds or other assay artifacts (Fig. 1a and 1c). The IC50 value of specific hits were then determined using the PRMT5 FP assay across an 8-point dose range. Only one cluster, based on a N-Aryl acetamide substituted dichloropyridazinone, as exemplified by compound **1** (Fig. 1b and Table 1) with an FP IC50 of 12 μM and low solubility (1.2 μM in PBS), was validated by SPR, NMR and X-ray crystallography.

**Table 1.**

| Compound Number | R<sub>1</sub> | PRMT5 IC50 (μM) | PBS Solubility (μM) | Plasma Stability (%Remaining) |  | GSH T ½ (min) |
| --- | --- | --- | --- | --- | --- | --- |
|  |  |  |  | Human | Mouse |  |
| 1 |  | 12.4 | 1.2 | 92 | 0 | 46 |
| 2 |  | 5.4 | 98 | 84 | 0 | ND |
| 3 |  | 12.7 | 0.8 | ND | ND | ND |
| 4 |  | 10.1 | 89 | 72 | 0 | ND |
| 5 |  | 14.1 | 56 | 89 | 0 | 25 |
| 6 |  | 2.2 | 47 | 82 | 14 | 31 |
| 7 |  | 4.6 | 88 | 100 | 21 | ND |
| 8 |  | 2.5 | 93 | 92 | 2 | ND |

Early hit exploration focused on improving solubility and potency of compound **1**. Analysis of commercially available analogs highlighted the requirement for the pyridazinone portion and as such, we began exploring the SAR of sulfonamide variations (Table 1). The majority of analogs were synthesized in five steps from available reagents (Suppl. Scheme 1). The azepane could readily be modified and solubility greatly enhanced by the introduction of a basic nitrogen (compound **2**, solubility 98 uM). Smaller rings (compound **4**, solubility 89 uM), as well as acyclic sulfonamides (compound **5**, solubility 56 uM), were both more amenable to rapid analog synthesis and retained significant potency (FP IC50 10 μM and 14 μM, respectively), opening the path forward for further SAR exploration. While these simple modifications significantly improved compound solubility, we found that pyridine isomers (compounds **6-8**) also led to an improvement in potency. Indeed, compound **6** showed a good balance of potency and solubility (Table 1). Modification of the aryl ring (Table 2), either by substitution of the methyl group (compounds **9-11**) or addition of a methyl group on open positions (compounds **12-14**) were tolerated, albeit with modest reduction in either potency, solubility, or both. Pyridine isomer **15** was tolerated, if poorly soluble, whereas other isomers **16, 17** were not, and saturated analog **18** demonstrated no ability to displace the PBM peptide. Modification of the acetamide at the center of these compounds (Table 3) afforded significant improvements in potency by the installation of single methyl group alpha to the carbonyl **20**, whereas the other enantiomer **21** and the N-methyl analog **19** were decidedly less active. Interestingly, these amide modifications all resulted in increased stability to mouse plasma, presumably by blocking amide hydrolysis by carboxylesterase, which is present in high concentrations in mice ^*13, 14*^.

**Table 2.**
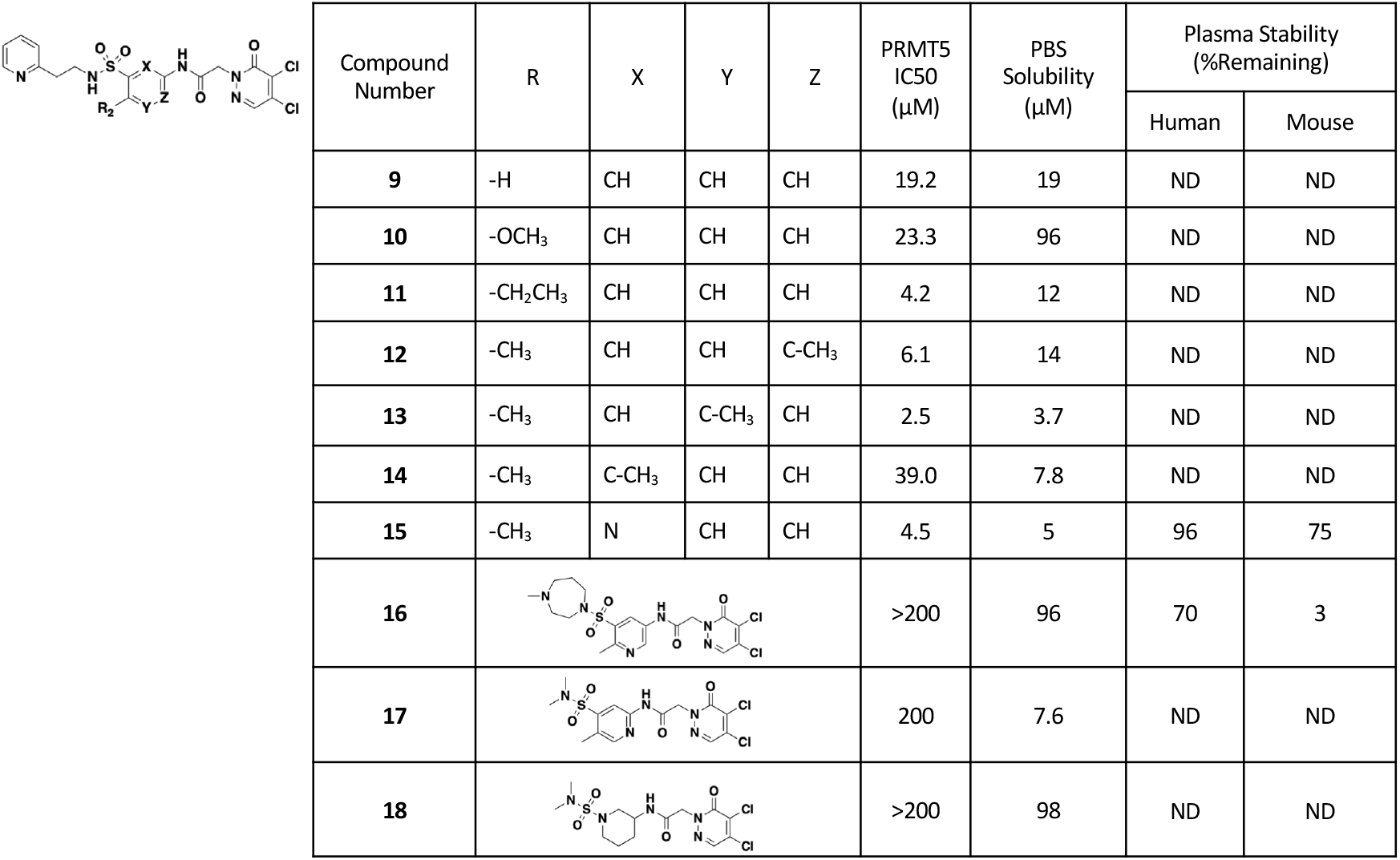

| Compound Number | R | X | Y | Z | PRMT5 IC50 (μM) | PBS Solubility (μM) | Plasma Stability (%Remaining) |  |
| --- | --- | --- | --- | --- | --- | --- | --- | --- |
|  |  |  |  |  |  |  | Human | Mouse |
| **9** | -H | CH | CH | CH | 19.2 | 19 | ND | ND |
| **10** | -OCH<sub>3</sub> | CH | CH | CH | 23.3 | 96 | ND | ND |
| **11** | -CH<sub>2</sub>CH<sub>3</sub> | CH | CH | CH | 4.2 | 12 | ND | ND |
| **12** | -CH<sub>3</sub> | CH | CH | C-CH<sub>3</sub> | 6.1 | 14 | ND | ND |
| **13** | -CH<sub>3</sub> | CH | C-CH<sub>3</sub> | CH | 2.5 | 3.7 | ND | ND |
| **14** | -CH<sub>3</sub> | C-CH<sub>3</sub> | CH | CH | 39.0 | 7.8 | ND | ND |
| **15** | -CH<sub>3</sub> | N | CH | CH | 4.5 | 5 | 96 | 75 |
| **16** |  |  |  |  | >200 | 96 | 70 | 3 |
| **17** |  |  |  |  | 200 | 7.6 | ND | ND |
| **18** |  |  |  |  | >200 | 98 | ND | ND |

**Table 3.**

| Compound Number | R<sub>3</sub> | R<sub>4</sub> | X | Y | PRMT5 IC<sub>50</sub> (μM) | PBS Sol (μM) | Plasma Stability (%Remaining) |  | GSH T<sub>1/2</sub> (min) |
| --- | --- | --- | --- | --- | --- | --- | --- | --- | --- |
|  |  |  |  |  |  |  | Human | Mouse |  |
| **6** | -H | -H | -Cl | -Cl | 2.2 | 47 | 82 | 14 | 31 |
| **19** | -CH<sub>3</sub> | -H | -Cl | -Cl | >200 | 86 | ND | ND | ND |
| **20** | -H | (S)-CH<sub>3</sub> | -Cl | -Cl | 0.3 | 64 | 92 | 92 | ND |
| **21** | -H | (R)-CH<sub>3</sub> | -Cl | -Cl | 55.4 | 70 | 95 | 97 | ND |
| **22** | -H | (S)-CH<sub>3</sub> | -Cl | -CH<sub>3</sub> | >200 | 83 | ND | ND | >3469 |
| **23** | -H | -H | -F | -Cl | 0.37 | 43 | 18 | 0 | <2 |
| **24** | -H | -H | -F | -CH<sub>3</sub> | 72.1 | 110 | ND | ND | 990 |
| **25** | -H | -H | -F | -H | 35.6 | 110 | ND | ND | 35 |
| **26** | -H | -H | -Cl | -H | 72.9 | 72 | 100 | 40 | 693 |
| **27 (BRD0639)** | -H | (S)-CH<sub>3</sub> | -Cl | -H | 13.8 | 82 | 87 | 79 | 916 |
| **28** | -H | (R)-CH<sub>3</sub> | -Cl | -H | 31.7 | 87 | 86 | 79 | 3105 |
| **29** | -CH<sub>3</sub> | (S)-CH<sub>3</sub> | -Cl | -H | >200 | 114 | ND | ND | ND |
| **30 (BRD2198)** | -H | -H | -H | -Cl | >200 | 90 | ND | ND | 3465 |

### Mechanism of Binding

To understand the mode of binding and support compound optimization, we generated a high resolution (1.9Å after elliptical truncation of anisotropic data) co-crystal structure of compound **1** with the PRMT5:WDR77 complex (Fig. 2a, PDB ID 6V0P, Suppl. Table 1 and Suppl. Fig. 1b). Only one site of new electron density was observed and overlapped with the known PBM binding site, confirming inhibition by direct competition. The refined electron density (Supp. Fig. 1a with 2Fo-Fc) suggested that the 4-position of the pyridazinone ring forms a covalent bond with PRMT5 Cys278 as a result of nucleophilic attack by the Cys278 thiol and subsequent loss of chlorine, driven by the re-aromatization of the pyridazinone. The crystal structure also points to key non-covalent interactions which are likely to drive initial binding and site specificity. The core aniline of the compound forms a pi-pi-stack with Phe243 and Tyr286, in a manner analogous to that formed with Phe230 of the Riok1 peptide^*11*^ (PDB ID: 6V0N). Several hydrogen bonds are also made between the compound and protein, including (1) between the compound sulfonamide and Asn239, (2) between the compound amide and Ser279, and (3) between Gln282 and the free nitrogen on the pyridazinone. Despite the high-resolution nature of the crystal structure, no unambiguous density was observed for the azepane functional group which might be due to its location at a crystal contact and this was left unmodelled (Supp. Fig. 1b).

**Figure 2.**
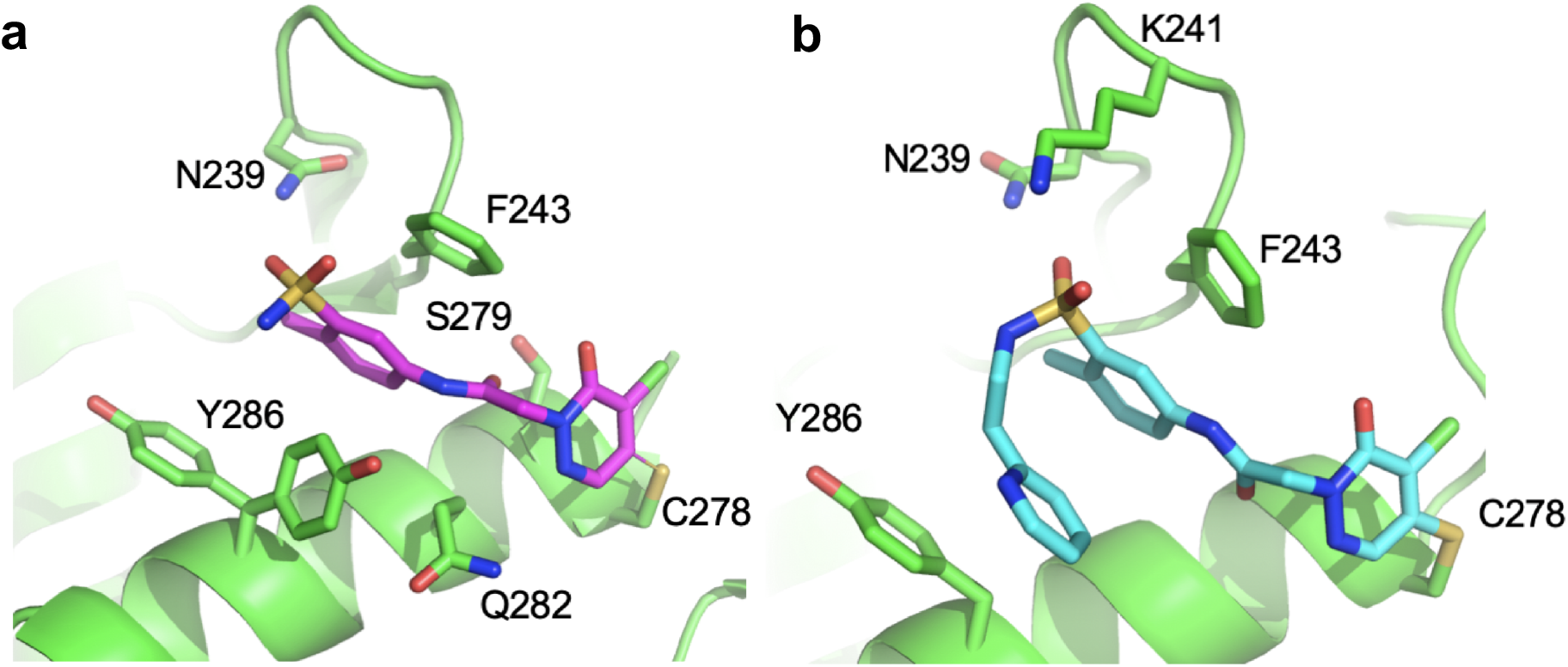
Mechanism of binding. **(A)** Crystal structure of compound **1** (magenta) bound to PRMT5 (green). Residues described in the text are labeled. Two alternate conformations of Y286 are shown. **(B)** Cryo-EM structure of compound **6** (cyan) bound to PRMT5 (green).

As significant potency improvements were made via substitution of the original azepane group, we sought to understand these changes from a structural perspective. However, none of the improved potency variants examined were amenable to high resolution crystallization likely due to the local crystal contact interface. We therefore solved a Cryo-EM structure of the PRMT5:WDR77 hetero-octameric complex bound to the pyridyl ethyl variant, compound **6** (Fig. 2b, Table 2 and Suppl. Fig 1c). This structure was solved to an overall resolution of 2.4Å, which represents a substantial improvement compared to previously published Cryo-EM structures of PRMT5 resolved at 3.4Å (PDB ID: 6UGH) and 3.7Å (EMD-7137 ^*15*^). The structure has excellent side-chain resolution and clear density for compound **6** (Fig. 2b and Suppl. 1c). The pyridyl ethyl position is well defined in the density and the other functional groups are in nearly identical poses compared to the X-ray structure of compound **1**. The pyridyl ethyl side chain is oriented back towards the core of the molecule via rotation of the sulfonamide and flexibility of the ethyl linker. This molecular conformation produces a new 4-ring stack of pi-pi interactions consisting of (from “front to back”) Tyr286-Pyridine-Aryl core-Phe243. Notably, this produces a folded-back structure analogous to the distinct conformation of the PBM peptide ^*11*^. Compared to the crystal structure of compound **1**, the density for Lys241 is better resolved and a hydrogen bond between this sidechain and the compound sulfonamide can now be modeled.

### Compound 6 is a covalent binder

We began exploring the putative covalent mode of compound binding by performing a time course FP analysis of compound **6**. Here IC50s were measured at 10 minutes intervals up to 4 hours. The observed IC50s decreased more than 10-fold over time, reaching the assay floor of ~100 nM (Fig. 3a). The covalent nature and site specificity of the compound was further verified using a PRMT5 complex with a Cys278 to Ala mutation. In FP competition experiments, this compound series, as exemplified by Compound **6** (blue squares), has little or no activity against the C278A mutant (red squares, 0.6 vs >100 μM) in contrast to a competitor PBM peptide which is comparable against both WT and C278A proteins (blue and red circles IC50 values 2.0 vs 1.4 μM, respectively) (Fig. 3b).

**Figure 3.**
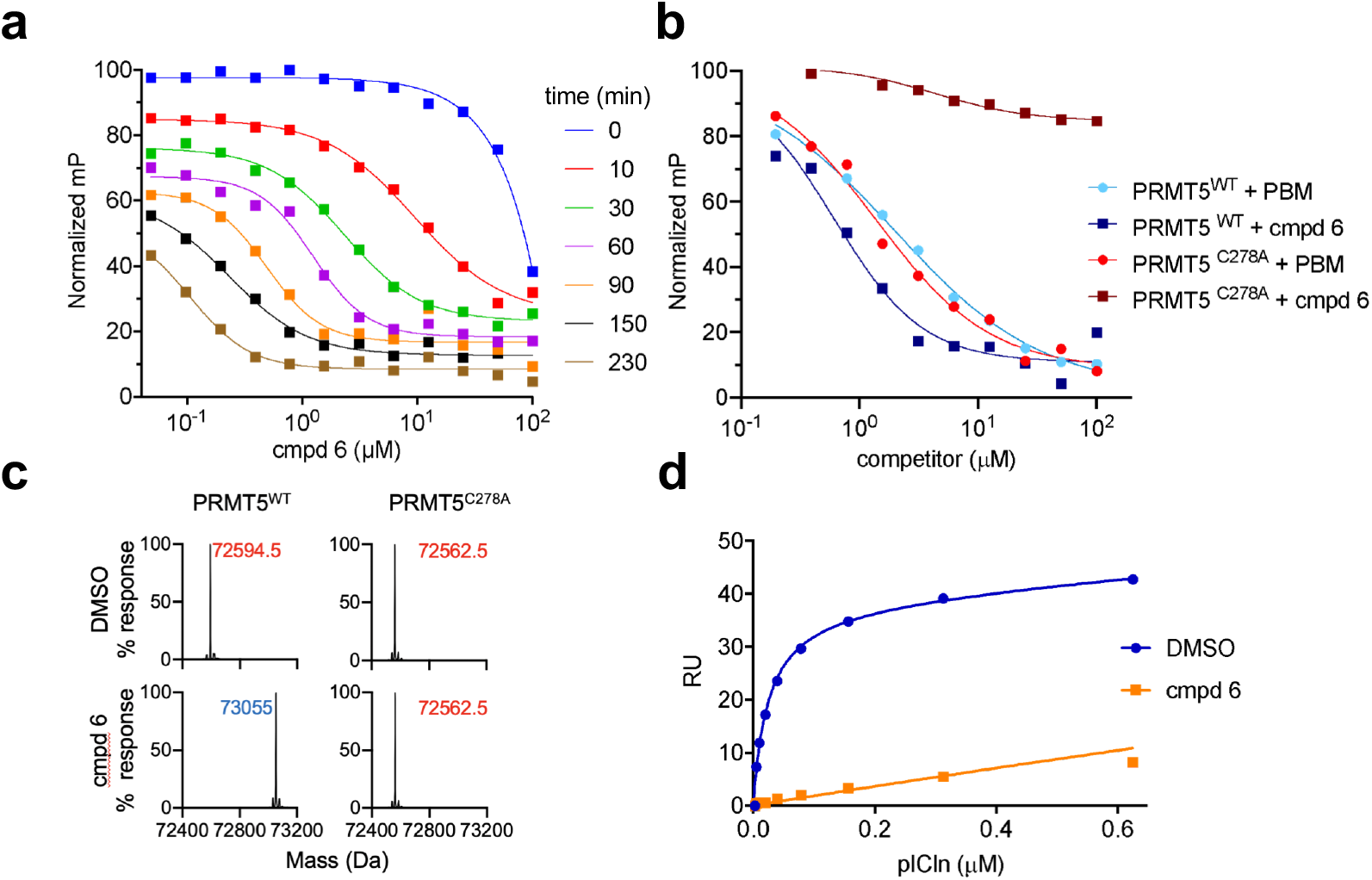
Compound 6 is a covalent binder. **(A)** compound **6** shows time-dependent displacement of a fluorescently-labeled PBM peptide probe as measured by FP **(B)** FP competition assay with compound **6** (squares) or pICln 13-mer control peptide (circles) displacing the fluorescently-labeled PBM probe from WT PRMT5 (blue) and C278A mutant (green). **(C)** Deconvoluted mass spectra for WT and C278A PRMT5 +/- compound **6 (D)** SPR competition assay, PRMT5:WDR77 complex was pre-incubated with compound **6** or DMSO and then immobilized to the chip surface. Full-length pICln protein was titrated as analyte.

In addition, intact mass measurement of the WT PRMT5:WDR77 complex before and after compound treatment showed an increase for PRMT5 of 460 Da with compound **6** (Fig. 3c) correlating with the expected mass of the compound, less one molecule of HCl that is lost upon covalent attachment of a single adduct to PRMT5. No change in the mass of WDR77 was observed (Supp. Fig. 2). Consistent with the above findings, treatment of PRMT5^C278A^ with compound **6** did not result in a mass shift, indicating that the covalent attachment was taking place at Cys278. For reference, the PRMT5 and WDR77 proteins contain 12 and 13 total cysteine residues, respectively, indicating selective binding.

Finally, we investigated whether pre-incubation with compound **6** was sufficient to block binding, as measured by SPR, between PRMT5 and the full-length SAP pICln. To do so, PRMT5 was preincubated with either DMSO or compound **6**, and then immobilized to the chip surface without further compound treatment. Titration of pICln to DMSO-treated PRMT5 produced binding with a K_D_ of 32 nM. In contrast, a greatly reduced binding response was observed for compound-treated PRMT5 consistent with linear, non-specific binding (Fig. 3d). Together, these results demonstrate that the compound binds by an irreversible covalent mechanism and reacts with a single cysteine on the PRMT5:WDR77 complex.

### Optimization of reactivity

Concerned that the highly reactive nature of these molecules could negatively impact their selectivity and off-target toxicity, we explored modifications of the warhead with the goal of decoupling potency from reactivity. The electrophilic reactivity of compounds towards the cysteine residue of glutathione (GSH) was evaluated and used as a surrogate for intrinsic reactivity ^*16*^. Not surprisingly, modifications outside of the warhead had negligible effects on reactivity to GSH, but affected FP potency (see acetamide, sulfonamide and phenyl ring substitutions, (Tables 1, 2). However, with changes at the pyridazinone warhead we observed a strong correlation between GSH reactivity and FP assay potency (Table 3). Of the various substitutions tested, the most interesting was the monochloro substitution, exemplified by compound **26**, which provided an acceptable balance between activity and reactivity (FP IC5040min 72.9μM; GSH T ½ 693 min). From **26,** the addition of a chiral alpha methyl substituent led to BRD0639, which, as observed before, significantly improved potency while retaining low reactivity (FP IC50_40min_ 13.8μM; GSH T ½ 916 min).

To quantify the contribution of various substitutions to both initial reversible binding (K_I_) and rate of maximum covalent reactivity (k_inact_), we developed a mass spectrometry-based k_inact_/K_I_ assay. The k_inact_/K_I_ ratio is considered the best time-independent measure of covalent compounds potency ^*17*^. In this assay, we incubate purified PRMT5:WDR77 protein with varying concentrations of the inhibitors and quench over a time course by rapidly reducing the pH with the addition of formic acid. Subsequent LC-MS analysis enabled the quantitation of the loss of unmodified PRMT5 represented as percent occupancy as a function of time across a range of concentrations (Fig. 4a). This allowed the generation of K_obs_ vs concentration curves (Fig. 4c, Supp. Fig. 3) from which we were able to generate k_inact_/K_I_ values for a number of inhibitors (Fig. 4c) ^*17*^.

**Figure 4.**
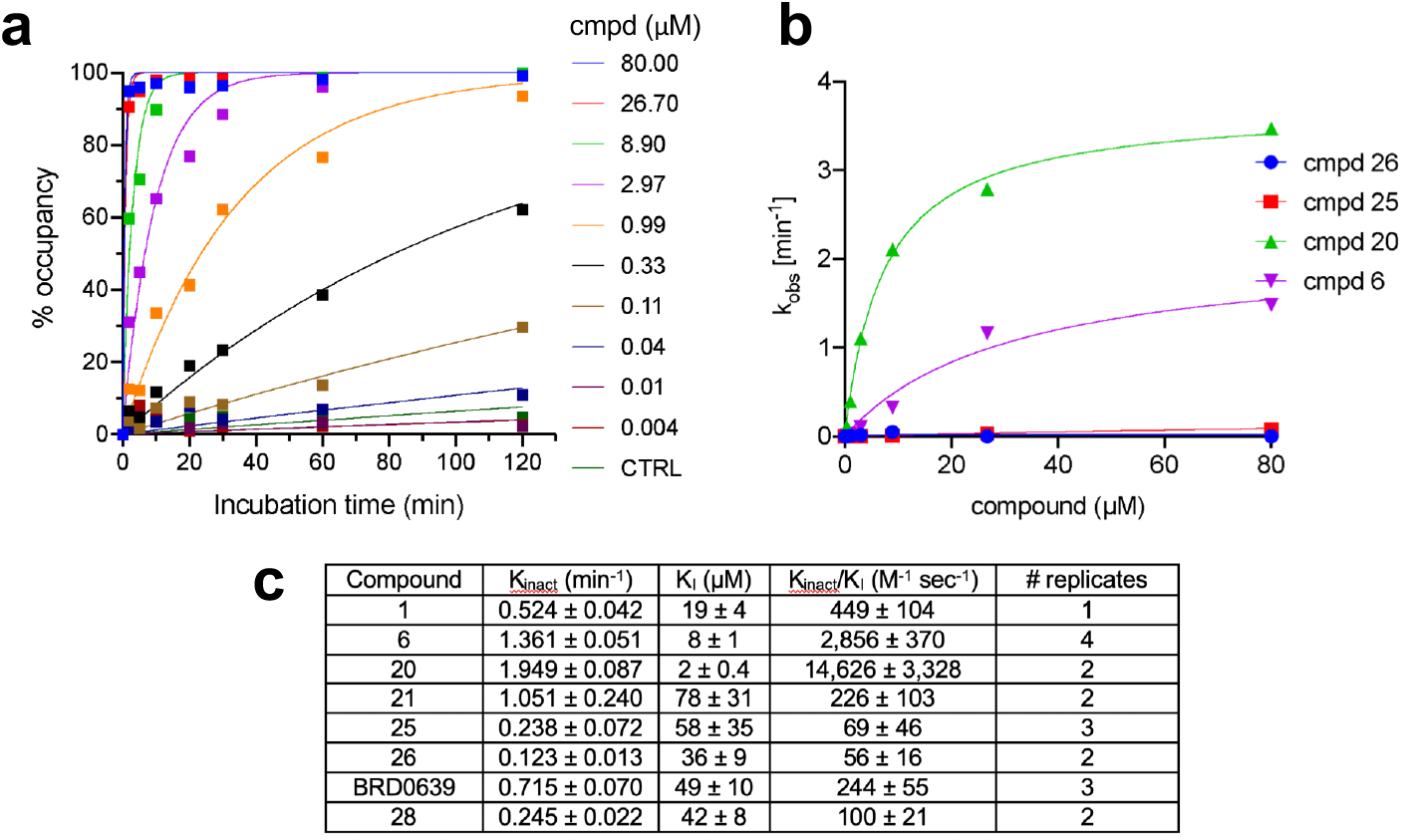
Kinetic formation of the covalent adducts between cmpd 6 and PRMT5. **(A)** Dose dependent kinetics of the compound **6**-PRMT5 reaction as determined by intact mass spectrometry **(B)** K_I_/K_inact_ analysis of covalent modification by compound **26** (blue), compound **25** (red), compound **20** (green), and compound **6** (purple) as determined by intact mass spectrometry **(C)** Table comparing kinetic parameters for select compounds

This assay revealed a high degree of correlation between the FP IC50_40min_ and the rate of complex formation for dichloro compounds (Fig. 4c). Indeed, a highly potent compound **20,** shows a greater than 30-fold increase in the overall rate of adduct formation, compared to compound **1** (k_inact_/K_I_ = 14,626 vs 449 (M^-1^ sec^-1^), respectively). Here, an improvement in both initial binding (K_I_) and maximum potential inactivation rate (k_inact_) contributed significantly to compound **20**’s increase in potency. The monochloro compounds show a different trend, where the potency difference observed between *S*-methyl compound BRD0639 and its non-methyl variant **26** by FP IC50_40min_ (13.8 vs 72.9 μM) correlates with overall rate of adduct formation (k_inact_/K_I_ = 244 vs 56 (M^-1^ sec^-1^), respectively). In this case, this is primarily the result of a higher k_inact_ (0.715 vs 0.123 M^-1^ sec^-1^), which compensates for a modest reduction in apparent K_I_ (49 vs 36 μM). Interestingly, dichloro **6** and monofluoro **25** have similar reactivity to GSH (T ½ 31 vs 35 min), but more than a 40-fold different rate of covalent modification of PRMT5 (k_inact_/K_I_ = 2,856 vs 69 (M^-1^ sec^-1^)), which correlates well with the observed 17-fold difference in FP IC50_40min_ (2.2 μM vs 35.6 μM, respectively).

Taken together, the above observations support that the nature of the leaving group, chlorine or fluorine, has significant effects on overall rate of adduct formation, not fully explained by their differences in intrinsic reactivity. The same is true for other modifications in close proximity to the leaving group. This kinetic analysis guided us to the monochloro pyridazinone as a low intrinsic reactivity warhead that retains good potency.

### Cellular activity of BRD0639: on-target and on-mechanism mode of action

To determine whether BRD0639 was able to engage PRMT5 in a cellular context, we treated Expi293 cells overexpressing HA-tagged WDR77 and WT PRMT5 for 6 hours and then analyzed anti-HA immunoprecipitates by LC-MS. This approach revealed a dose dependent formation of PRMT5-adducts (Fig. 5a). An EC50 of 3 μM was observed, however only ~40% of total PRMT5 protein was labeled by **27**, possibly related to the high protein production rate in this overexpression system.

**Figure 5.**
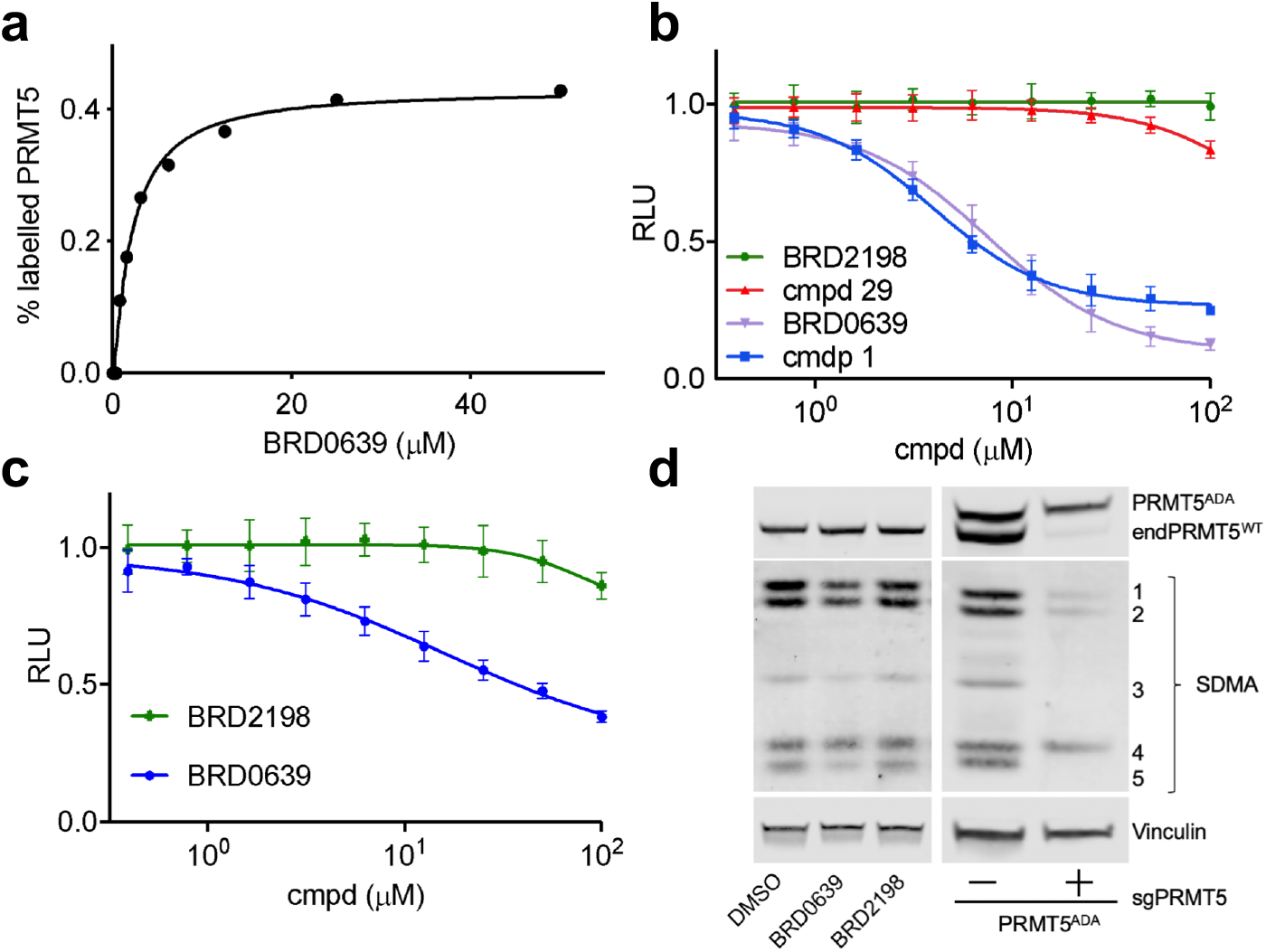
Cellular activities of BRD0639. **(A)** Time course of PRMT5 adduct formation using BRD0639. Expi293 cells were transiently transfected for 48 hrs with an HA-tagged PRMT5:WDR77 complex before treatment with BRD0639. Cultures were harvested at 6 hours, PRMT5 complex isolated by HA-affinity, and analyzed for modification by LC/MS. **(B)** PRMT5-RIOK1 NanoBiT assay in permeabilized cells. 293T cells stably expressing SmBiT-PRMT5 and LgBiT-RIOK1 proteins. **(C)** NanoBit assay performed in intact cells after treatment with BRD0639 or BRD2198. **(D)** Left panel: WB analysis of total symmetric dimethylation levels in MTAP-/- HCT116 cells, in response to BRD0639 or BRD2198. Right panel: WB analysis of total symmetric dimethylation levels in MTAP-/- HCT116 cells overexpressing the PRMT5^ADA^ mutant which is unable to bind PBM peptide (Mulvaney, 2020), with and without KO of endogenous PRMT5.

To study the functional consequences on PRMT5-SAP interaction following the engagement of the PBM groove, we used a PRMT5-RIOK1 NanoBiT assay ^*11*^. Compounds **1** and BRD0639 were tested alongside related inactive compounds **29** and BRD2198 in permeabilized cells (Fig. 5b). These experiments showed that both compounds **1** and BRD0639, but none of the inactive compounds, disrupt the PRMT5-RIOK1 complex with an IC50 of 4 μM and 7.5 μM, respectively. Next, we assessed whether BRD0639 was able to engage the target in intact, non-permeabilized cells. To this end, we treated PRMT5-RIOK1 NanoBit expressing cells with either BRD0639 or BRD2198 for 40 minutes and then assessed the stability of the PRMT5-RIOK1 complex (Fig. 5c). Consistent with what was observed in permeabilized cells, BRD0639 but not BRD2198 was able to disrupt the complex (IC50 16 μM vs >100 μM). Finally, cellular treatment with BRD0639 resulted in inhibition of PRMT5 methyltransferase function, as demonstrated by the reduction in symmetric dimethylation levels by WB (Fig. 5d). We found that methylation of some, but not all substrates, were inhibited by treatment with BRD0639. Notably, this phenotype closely recapitulates the one observed following genetic disruption of the PBM-PBM groove interaction^*11*^ (Fig. 5d). Here, a PBM interaction site mutant of PRMT5 (PRMT5^ADA^) can rescue the methylation of some PRMT5 “PBM-dependent” substrates (Fig. 5d, arrows) but not others, after CRISPR KO of endogenous PRMT5. This finding strongly supports the conclusion that BRD0639 is on target and on mechanism.

## Discussion

We report here the discovery and characterization of a first-in-class PRMT5-PBM competitive inhibitor. Our recent characterization of the binding interface between PRMT5 and its SAPs enabled the development of a robust screening system to identify PBM-competitive small molecule inhibitors. Despite our considerable efforts in hit identification using three independent screening approaches (a small molecule FP-based HTS, an NMR-based fragment screen, and a virtual pharmacophore screen), we identified only one chemical series. As proteinprotein interaction sites can be challenging, our results leave open the possibility that larger and more diverse libraries, potentially including DNA-encoded libraries, might yield additional scaffolds.

Our only validated hit was a highly reactive covalent binder. Other liabilities of our initial hit were poor aqueous solubility and high instability in mouse plasma. Our hit-to-lead efforts resolved the solubility and the plasma instability issues and significantly reduced the intrinsic GSH reactivity to levels comparable with or below that for clinically-approved acrylamide warheads^23^. In addition to a robust FP system for HTS and routine potency measurements, we developed a novel LC-MS assay to quantitatively measure K_I_ and K_inact_ to more fully understand the effects of compound modifications. This system revealed that key points of compound modification, such as the azepane to ethyl pyridine substitution and *S*-methyl addition, made significant improvements in reversible binding without otherwise altering the maximum rate of reactivity. Nonetheless, it is clear that the covalent reaction is a necessary component of compound activity, which proved difficult to remove while still retaining potency. However, we were able to tune reactivity to acceptable levels via alteration from the initial dichloro-pyridazinone to a monochloro. Importantly, our lead compound BRD0639 can engage the cellular target and effectively outcompete binding between full-length PRMT5 and RIOK1 proteins with an IC50 of 7.5uM and 16 uM in permeabilized and living cells, respectively. Moreover, we demonstrate a pharmacodynamic response to the most proximal marker of PRMT5 activity, the formation of symmetrically dimethylated arginine. This effect appears to be on-target as closely related compounds with the same warhead, but inactive in FP assays, are also inactive in the cellular context. Consistent with an on-target effect, BRD0639 reduces SDMA in the same subset of proteins also affected by genetic perturbation of the PBM binding site^*11*^.

The value of a specific PBM-competitive inhibitor lies in the differential therapeutic effect that can be achieved, as compared to that of a catalytic inhibitor. Indeed, PRMT5 plays key roles in regulating multiple essential cellular activities, including transcription, ribosomal biogenesis and mRNA splicing. The latter is mediated by PRMT5-induced symmetric dimethylation of Sm proteins, and requires its interaction with the substrate adaptor, pICln. Genetic disruption of the PBM-PBM groove, by impeding PRMT5-pICln binding, leads to a selective impairment of this function ^*11*^. Available inhibitors of the PRMT5 catalytic pocket, by inhibiting all PRMT5-mediated methylation events, perturb all of its activities, and could result in excessive toxicity. Conversely, pharmacological inhibition of PBM-PBM groove interaction would only affect mRNA splicing, and could lead to a much greater therapeutic index in specific indications.

Moving forward, it would be interesting to also explore additional strategies for targeting the PRMT5 methylosome. It is notable that the relevance of PRMT5 to MTAP-deleted cancers was observed by shRNA knockdown, suggesting that partial reduction in total PRMT5 protein concentration would also be sufficient to see selective viability effects. As such, PRMT5 degraders could represent an alternative therapeutic approach to this target. It should be noted that the development of heterobifunctional degraders directed against PRMT5 have been reported recently ^*18*^. However, their potency is limited, possibly due to the fact that conventional PRMT5 catalytic inhibitors were used as the target bait. Here, we believe that the depth of the catalytic pocket might pose a structural challenge in connecting a catalytic PRMT5 inhibitor, via a linker, to an E3-ligase binder. From this perspective, the discovery of a new surface exposed and druggable site on PRMT5, the development of a fully validated suite of assays and the generation of a tool compound to support screenings for new, non-covalent binders, could offer a unique starting point for the development of potent PRMT5 degraders.

## METHODS

### Fluorescence polarization

The FP competition assay had final concentrations 200 nM PRMT5:WDR77 protomer, (experimentally determined K_D_ for the interaction ^*11*^), 10 nM peptide probe, 50 mM HEPES pH 7.4, 100 mM NaCl, 0.5 mM TCEP, and 0.01% v/v Tween 20. For HTS, 1536-well black non-binding surface plates (Corning) were used with a 5 μL assay volume. The assay reagents were added to pre-plated compounds with a final compound concentration of 20 μM. The plates were incubated at room temperature and the final endpoint read at 40 minutes. For SAR data, experiments were performed in triplicate in 384-well black non-binding surface plates (Corning) at a final assay volume of 20 μL. Data were collected using either a Spectramax Paradigm with Rhodamine FP filter set or a Perkin-Elmer Envision with Bodipy TMR FP filter set. The peptide probe was a KU560 fluorophore (KU dyes, catalog KU560-R-6)-labeled peptide derived from the RIOK1 PBM sequence: [acetyl]SRVVPGQFDDADSSD[C^KU560][amide]). As a positive control, peptide [acetyl]LMSRVVPGEFDDADSSD[amide] was used at a 20 μM concentration.

### Nuclear Magnetic Resonance

Experiments were performed on a 600 MHz Bruker Avance III Spectrometer equipped with a 5 mm QCI cryoprobe and a SampleJet for automated sample handling. All experiments were conducted at 280 K. For. For STD NMR^19, 20^ final sample conditions involved 200 μM ligand, 2% DMSO, 1 μM protomer, 25 mM HEPES-d, pH=7.4, 150 mM NaCl, 1 mM TCEP-d, followed by competition with 20 μM 13-mer peptide (K_D_ 300 nM). All NMR data was analyzed using Topspin (Bruker). The SPY peptide was synthesized by Thermo-Fisher, sequence: [acetyl]GQF*EDAD[amide], where F* is 3-Fluoro phenylalanine; ^19^F signal at −113.75 ppm.

### Surface Plasmon Resonance

Biacore experiments were performed as previously described ^*11*^ with the exception that PRMT5 protein was pre-incubated at a concentration of 25 nM with 2 μM compound **6** or DMSO overnight at 4C prior to immobilization. Labeling was confirmed to be complete by LC-MS. Data were fit in Prism using a one site, total and non-specific binding model.

### Crystallography

Crystals were produced as previously described ^*11*^ and then soaked with compound **1** at an approximate concentration of 500 μM for 72 hr before harvesting. Diffraction images were indexed and integrated using XDS and further processed for anisotropy via elliptical truncation using the STARANISO server (Global Phasing). Cambridge, United Kingdom: Global Phasing Ltd.). Refinement was performed in Buster (Global Phasing) and Phenix with manual building/review in Coot. Compound restraints were generated using Glide (Global Phasing).

### CryoEM

PRMT5:WDR77 complex at a 5 μM concentration was co-incubated with 25 μM compound **6**, and 20 μM JNJ-64619178 overnight at 4C. Covalent modification was confirmed by LC-MS. PRMT5 complex was then purified by size exclusion on a Superose 6 Increase 10/300 GL (GE Healthcare) column in mobile phase buffer 10 mM HEPES pH 7.4, 150 mM NaCl, 10% glycerol, 1 mM TCEP, and 100 nM JNJ-64619178. Protein was concentrated to 10 mg/ml using a Proteus X-spinner with 10 kDa filter (Anatrace) and snap frozen in liquid nitrogen. UltrAuFoil 300 mesh grids were pretreated by glow discharge for 30 sec. Immediately prior to application, protein samples were thawed and diluted 7-fold in a buffer containing 10 mM HEPES pH 7.4, 150 mM NaCl, 1 mM TCEP and 100 nM JNJ-64619178. Using a FEI Vitrobot Mark IV, 3.5 μl of the protein sample was applied to the grid before blotting and plunge-freezing into liquid ethane. Data were acquired on a Titan Krios microscope with Gatan K2 Quantum detector at a nominal magnification of 130,000x and randomized defocus values of −1.4, −1.7 or −2.1 Å. For each image, 40 movie frames were recorded with a total exposure time of 8 sec and estimated electron dose of 62.4 e^-^/Å^2^. A total of 2169 movies were collected from one grid using automatic acquisition. All data processing was performed in the cisTEM software suite^22^. Frames 4-40 from each movie were motion corrected, summed and then processed by CTF estimation. All images with a CTF fit resolution ≤ 3.1 Å (1836 images) were retained for single particle analysis. A total of 956,646 particles were picked for 2D classification. Due to particle crowding, only 474,404 particles (22 of 50 classes) were selected for 3D refinement. EMD-7137 was used as a starting volume model with an initial high-resolution limit of 20 Å and a defined D2 symmetry. 3D refinement converged at an estimated resolution of 2.39 Å using half-map analysis with a 0.143 FSC cutoff. Phenix Autosharpen was used prior to rigid body docking of the 6V0P crystal structure as a starting model. Manual model building was performed in Coot followed by real space refinement in Phenix.

### Stability to GSH

Ten μl of compound at a final concentration of 0.1, 1 and 10 μM, or control working solution, were diluted in 190 μl of 5 mM glutathione (GSH) in PBS (pH 7.4) and incubated at 37°C for 0, 15, 30, 60, 120 and 1440 min, in duplicate. As a negative control, incubations without GSH were carried out at two time-points (0 and 1440 min). Ibrutinib and afatinib were used as positive controls and were tested at a final concentration of 10 μM. At each time-point, the reaction was terminated by adding 600 μl cold acetonitrile containing labetalol as the internal standard. The samples were stored at - 80°C until the last incubation time-point was completed. Sample plates were defrosted and placed on a shaker for 5 min, followed by centrifugation at 4000 rpm for 20 min. 100 ml of each sample, after centrifugation, was further diluted with water before mass spec analysis. Samples with compound concentration at 0.1, 1 and 10 mM were diluted with 100 μl, 300 μl and 600 μl water, respectively. Samples were analyzed by LC-MS/MS (Sciex API 4000). The percent of the parent compound compound remaining at each time-point was determined based on peak area ratios at the 0 min time-point and half-life was calculated using the first order kinetics equation. In addition, the samples were analyzed for formation of the predicted GSH adduct at each time-point.

### Intact mass measurement by LC-MS

Purified PRMT5:WDR77 was thawed on ice and centrifuged to remove potential aggregates from the freeze/thaw process. The protein was then solvent exchanged into the reaction buffer (10 mM HEPES 7.4, 150 mM NaCl, 1 mM TCEP, using a 40 kDa Zeba desalting column (ThermoFisher), and diluted to 10 μM. For single time point experiments, a mixture of 10 μL of protein, 1 μL of compound **6** (0.5 mM in DMSO) or DMSO and 90 μL reaction buffer were incubated at RT for 4 hours. Intact mass measurement of the PRMT5:WDR77 complex with and without compounds was performed using the BioAccord LC-ToF (composed of an ACQUITY I-Class UPLC and RDa detector with ESI source, Waters Corporation). 1 μl of each sample was injected onto a C4 column (ACQUITY UPLC Protein BEH, 300Å, 1.7 μm, 2.1 × 50 mm, Waters Corporation) held at 80 °C. Mobile phases A and B consisted of 0.1% formic acid (MilliporeSigma LiChroPur) in LC-MS grade water or LC-MS grade acetonitrile (JTBaker), respectively, with initial column conditions set to 95% water/5% acetonitrile. Protein was desalted for one minute before elution with a gradient of 5% to 85% mobile phase B in 2.5 min followed by ionization in positive ionization mode with the cone voltage set to 55 V and desolvation temperature of 550 °C. The instrument scan rate was 5 Hz over 50 to 2000 m/z. PRMT5 and WDR77 coeluted at 2.38 minutes and mass spectra were deconvoluted using UNIFI and the MaxEnt1 algorithm.

### Covalent modification of PRMT5 and kinetic analysis (k_inact_/K_I_) by LC-MS

Purified PRMT5:WDR77 was thawed and treated as described in the previous section. For k_inact_/k_i_ analyses, PRMT5:WDR77 was diluted to 50 nM and dispensed into 384-well plates (Greiner Bio-One #781280) containing a 12-point dilution series of various compounds to collect 8 time points (0, 2, 5, 10, 20, 30, 45, and 60 min) using a 384ST-head Agilent Bravo. Time points were quenched with formic acid (LiChropur, MilliporeSigma, final concentration 0.5%). To quantify the unmodified PRMT5 and WDR77 a multiple-reaction monitoring (MRM) method was developed using a Waters ACQUITY UPLC I-Class PLUS chromatography system connected to a Xevo TQ-XS mass spectrometer by focusing on a single charge state for each protein (PRMT5, m/z 855.0 → m/z 854.90, z = 85; WDR77, m/z 870.80 → m/z 870.75, z = 46). ESI source parameters were set as follows: positive ionization mode, capillary voltage 2.5 kV, cone voltage, 50 V; collision energy, 5 V; cone gas flow, 200 L/h; collision gas flow, 0.16 mL/min; source temperature, 150 °C; desolvation temperature, 350 °C; and desolvation gas flow, 900 L/h. Analytes were separated on an Agilent PLRP-S column (5 μm; 2.1 × 50 mm, 1000Å Agilent Technologies) at 60 °C. Sample storage temperature was set to 20 °C. Mobile phases and initial conditions were the same as described above. Proteins were eluted using a 1.5 min gradient of 5% to 98% acetonitrile at 0.4 ml/min. MassLynx and TargetLynx software (Waters, version 4.2) were used for sample acquisition and data quantification, respectively. Analytes were quantified by integration of peak areas for each charge state described above. For analysis, the peak areas were normalized to the 0 min time point. Kinetic analysis of the k_inact_ and K_I_ values were fit in Graphpad Prism 7 ^17^.

### Target engagement in cells

Expi293 cells were cultured in a 1:1 mixture of Expi293:Freestyle media (Thermofisher). At a cell density of ~2.5×10^6^ cells/ml, 250 ml of cells were transfected using 200 μl FectoPRO (Polyplus Transfection) reagent mixed with 100 μg PRMT5 and 100 μg HA-tagged WDR77 plasmids ^*11*^. One day after transfection, 3 mM valproic acid and 0.4% w/v glucose were added to the culture. Two days after transfection, cells were split into multiple flasks with 30 ml cells each and treated with compound or DMSO at a final 0.2% DMSO concentration. Cell sample was collected after 6 hours, pelleted and washed 3 times with ice cold PBS. Additional aliquots were analyzed for viability and cell number using a Vi-Cell XR Cell Counter (Beckman Coulter) and all samples were determined to have >98% viability based on trypan blue exclusion. Each sample was lysed in buffer 50 mM HEPES pH 7.5, 300 mM NaCl, 1 mM TCEP, 1% v/v Tween-20 and 2 mM reduced glutathione. The PRMT5:WDR77 complex was immunoprecipitated by anti-HA agarose resin (Thermofisher) and then eluted by 50 μM 3X-HA peptide (AnaSpec). Each eluate was measured by intact mass LC-MS to determine the percentage of complex with compound adduct.

#### NanoBiT

HEK293T cells stably expressing PRMT5 tagged with an N-terminal SmBiT peptide and RIOK1 tagged with N-terminal LgBiT peptide were described before ^*11*^. Cells were cultured at 37°C in 5% CO_2_ in Dulbecco’s modified Eagle’s medium (DMEM, Life Technologies) supplemented with 10% FBS (Life Technologies) and 1% penicillin-streptomycin (Life Technologies). For the assay, cells were centrifuged at 150 rad/s, media removed, and resuspended in Opti-MEM (Life Technologies). Cells were plated at (10×10^4^ cells/ well) in 96well black wall, clear bottom, tissue culture treated plates (Corning) and treated with the indicated compounds using D300e Digital Printer (Tecan). All wells normalized to highest DMSO concentration (1% v/v). Each experimental condition was tested in triplicate.

##### NanoBiT in permeabilized cells

50uL of lysis buffer was added to each well (10% glycerol, 50mM Tris-HCL, 150mM KCl, 2mM EDTA, 0.1% NP40) and cells were incubated for 5min at RT. Nano-Glo^®^ Luciferase Assay Substrate (Promega #N1110) was diluted 1:50 in buffer, 100uL added to each well and pipetted to mix. *NanoBiT in intact cells*: Cells were treated for 40min, at which point Nano-Glo^®^ Live Cell Substrate (Promega, # N2011) was diluted 1:20 in buffer, 25uL mixture was added to each well and the plate tapped gently to mix.

Luciferase signal was measured using the EnVision plate reader (Perkin Elmer). Results shown are representative of three independent experiments.

#### Western Blot

HCT116 MTAP -/- were cultured at 37°C in 5% CO_2_ in Dulbecco’s modified Eagle’s medium (DMEM, Life Technologies), supplemented with 10% FBS (Life Technologies) and 1% penicillin-streptomycin (Life Technologies). Cells were plated at 1×10^6^/well in 6-well tissue culture treated plates (Corning), and treated the next day with the indicated compounds to a final concentration of 25uM. DMSO and non-treated control wells included. 12hr after treatment, the media was refreshed and cells retreated. 12hr later, cells were washed 1X with PBS and lysed on ice for 15min in 50uL lysis buffer (1mL RIPA, 10% glycerol, 1% protease inhibitor, 1% phosphatase inhibitor). Protein concentration was calculated using Pierce BCA Protein Assay Kit, 40ug protein per sample was run on a 4-12% Bis-Tris gels (NuPAGE, Life Technologies) using MES buffer. Gels were dry-transferred to nitrocellulose membrane (iBlot system, Life Technologies). Membranes were blocked using Intercept Blocking Buffer (LI-COR) for 1hr and probed overnight with primary antibodies (Rabbit anti-SDMA, CST13222; mouse anti-vinculin, Sigma, RABBIT anti-PRMT5 (Abcam Ab31751). Blots were washed 3X with 1% TBST buffer and probed with secondary antibodies 680RD goat anti-Mouse and 800CW goat anti-Rabbit (LI-COR) for 1hr. Membranes were washed 3X with 1% TBST and imaged using LI-COR Odyssey imaging system. Results shown are representative of three independent experiments.

## Chemical Synthesis

## See Supplemental information

## Contributions

A.I., D.C.M. and B.J.M. directed project planning and execution. W.R.S. supervised the overall project. WRS and B.J.M developed the therapeutic concept of PBM inhibition. A.I. and D.C.M. wrote the manuscript with input from B.J.M., M.R., J.A.M., P.M., M.M., and W.R.S.

V.K.K., D.P., B.J.M., D.C.M., M.M. and P.M. conceived and coordinated the hit finding activities. P.M. designed compounds and conducted the pharmacophore screen. M.M. conceived and executed the NMR fragment screen. B.B. and M.B. developed the MS KInact/KI assay, established analysis workflow, interpreted results. J.A.M. and M.R. assisted with MS analysis and experimental interpretation of the KInact/KI assay. B.J.M. designed and supervised the execution and data interpretation of the FP assays and performed the crystallography. B.J.M. and D.E.T. performed and interpreted the CryoEM. R.S. assisted with ADME data interpretation and compound design. K.M.M. developed the Nanobit assay in permeabilized cells. D.C.M. designed compounds and assisted with overall data interpretation. A.I. designed and supervised the execution of cell-based assays and data interpretation. B.J.M., D.C.M., M.R., MB, M.O., A.S. and Z.M-B. performed the experiments. All authors provided critical feedback which helped shape the research and reviewed the final manuscript.

## Supplemental Information

### NMR-based fragment screening

This screening was conducted using a ‘rule of three’ compliant library of 1,920 fragments supplied from commercial vendors using a pool size of 8 fragments per sample. To achieve a suitable probe for low affinity fragment competition, a shorter peptide sequence, acGQ(F*)EDADam (14.3 μM IC50 by FP) was used as the spy molecule, which included a synthetic 3-fluorophenylalanine to allow measurement of displacement via ^19^F-NMR line-broadening. For fragment screening, a compound concentration of 500 μM was used^21^. No molecules were identified that were able to displace this peptide using a 20% change in 19F signal intensity as a cutoff. The same fragment library was also used for ligand observed STD-NMR screening in the presence (PBM site blocked) or absence (PBM site available) of a ten-fold excess of the RIOK1 13-mer PBM peptide. NMR samples were prepared by pooling the fragments just-in-time into a 96-well plate using an Echo dispenser and the pre-mixed protein/peptide solution was added to the plate. Samples were then transferred to 3 mm NMR tubes using a GilsonPrep system. For the ^19^F-based screen, spectra were acquired using a 160 msec Carr-Purcell-Meiboom-Gill filter (CPMG, D/2 = 20 msec, N = 4) to enhance the effects of ligand binding and displacement of the peptide. For each spectrum, 128 scans were acquired over a sweep width of 20 ppm with a repetition time of 3 seconds. Experiments were conducted using a 1 μM protomer concentration, 25 μM spy probe, and with the catalytic site occupied using a ten-fold molar excess of the SAM binding site small molecule JNJ-64619178. The ligand-observed screen was conducted using STD-NMR^19, 20^. On-resonance irradiation of the protein was done at −0.25 ppm and off-resonance irradiation at 30 ppm. To saturate the protein, a 2 s train of 50 msec gaussian pulses separated by 1 msec delays was used. A 27 msec spin-lock pulse was used to suppress protein signals, and water suppression was accomplished using the excitation sculpting with gradients pulse scheme.

#### Virtual screening

The pharmacophore virtual screen was performed using MOE^*2*^ using an EHT pharmacophore query based on the pICln-PRMT5:WDR77 structure (PDB ID: 6V0O) and a pre-enumerated database containing 8.6M structures from 8 major vendors called CoCoCo^3^. The query was constructed using sidechain pharmacophore points from D8, F6, and Q5 (anion, aromatic, and hydrophobic at Cβ respectively) and backbone carbonyl points for E7 and Q5 (hydrogen bond acceptor). A docking virtual screen based on the pICln protein binding site was also performed using the Schrodinger virtual screening workflow (Glide HTVS and Glide SP only)^4,5^. A pre-enumerated and conformer expanded library totaling 22M compounds from the eMolecules commercially available compounds provided by Schrodinger using default settings in ligprep and confgen was used as input. The docking grid was constructed using the pICln-PBM bound protein prepared using the Schrodinger Protein Preparation Workflow. The pICln binding site was defined by the residues 5-8 (QFED) of the PBM peptide and using a default grid box size. Hits in both screens were filtered to lead-like properties using the Oprea leadlike criteria in MOE 2018 ^*6*^ and using Lilly MedChem Rules with default settings ^7^. The top 1000 resulting compounds in each method were clustered by 2D fingerprint and the top compounds per cluster were inspected. A total of 672 compounds were purchased for screening from the two methods and included with other screening collections in the FP screening.

**Supplemental Table 1.** Data collection and refinement statistics for X-ray crystal structure.

|  | Compound 1 |
| --- | --- |
| Data collection |  |
| Space group | I 2 2 2 |
| Cell dimensions |  |
| a, b, c (Å) | 99.11, 138.57,<br>178.42 |
| $\alpha$, $\beta$, $\gamma$ (°) | 90, 90, 90 |
| Diffraction limits (Å), $CC_{1/2} > 30\%$ | |
| a* | 2.75 |
| b* | 2.45 |
| c* | 1.83 |
| Resolution (Å) | 45-1.88 (2.10-1.88) |
| $CC_{1/2}$ (%), ellipsoidal | 99.8 (48.0) |
| R-meas (%), ellipsoidal | 11.5 (160.1) |
| $\langle I / \sigma I \rangle$, ellipsoidal | 32.7 (1.1) |
| Completeness (%), ellipsoidal | 93.9 (86.0) |
| Completeness (%), spherical | 50.8 (8.7) |
| Multiplicity | 6.5 (5.5) |
| Refinement |  |
| Resolution (Å) | 36.73-1.88 |
| No. reflections | 50976 |
| $R_{work} / R_{free}$ | 0.187/0.230 |
| No. atoms |  |
| Protein | 7258 |
| Ligands | 64 |
| Solvent | 637 |
| B-factors |  |
| Protein | 45.1 |
| Ligand | 80.98 |
| Solvent | 41.68 |
| Ramachandran |  |
| Favored (%) | 97.1 |
| Allowed (%) | 2.7 |
| Outliers (%) | 0.2 |
| R.m.s. deviations |  |
| Bond lengths (Å) | 0.014 |
| Bond angles (°) | 1.76 |
| Clashscore | 1.38 |
| Rotamer outliers (%) | 2.97 |

**Supplemental Table 2.**
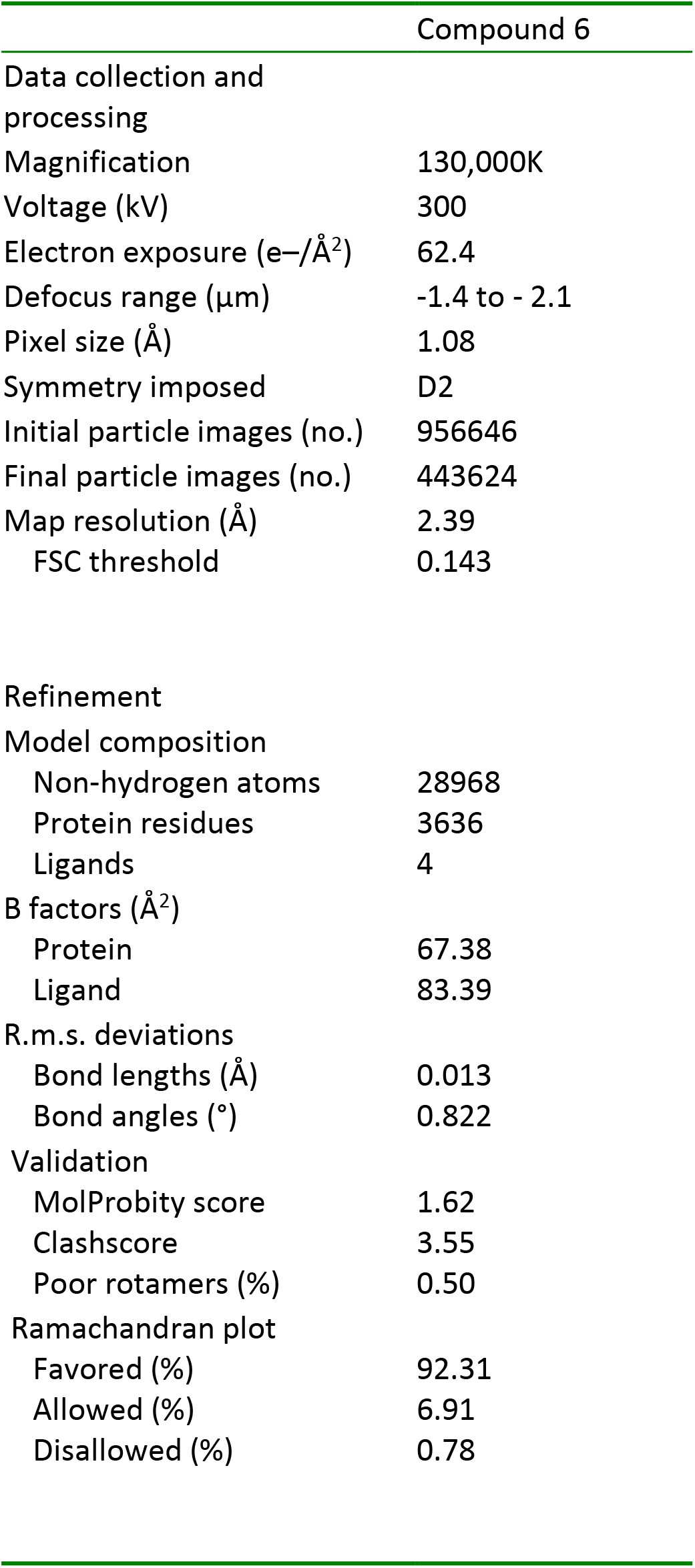
Data collection and refinement statistics for CryoEM structure.

**Supplemental Fig. 1.**
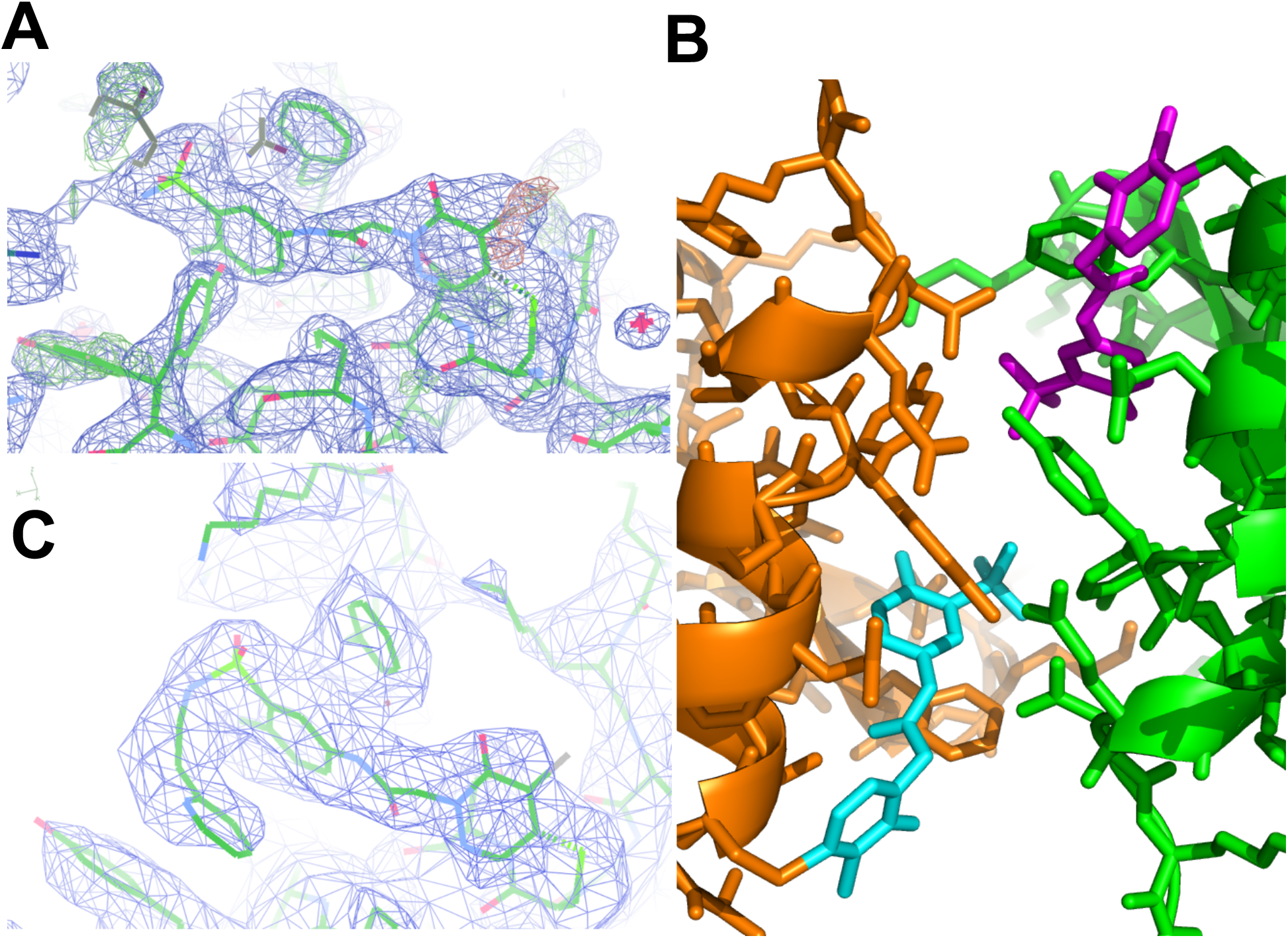
**(A)** Crystal structure of compound **1** bound to PRMT5 (carbon, green). 1σ 2Fo-Fc density in blue, 3σ Fo-Fc in red/green. A crystal contact neighbor is in grey. **(B)** Crystal contact between two neighboring monomers of PRMT5 (green and orange). Compound 1 is shown in magenta and cyan. **(C)** Cryo-EM structure of compound 6 bound to PRMT5 (carbon, green). Electron density is shown at 5σ.

**Supplemental Fig. 2.**
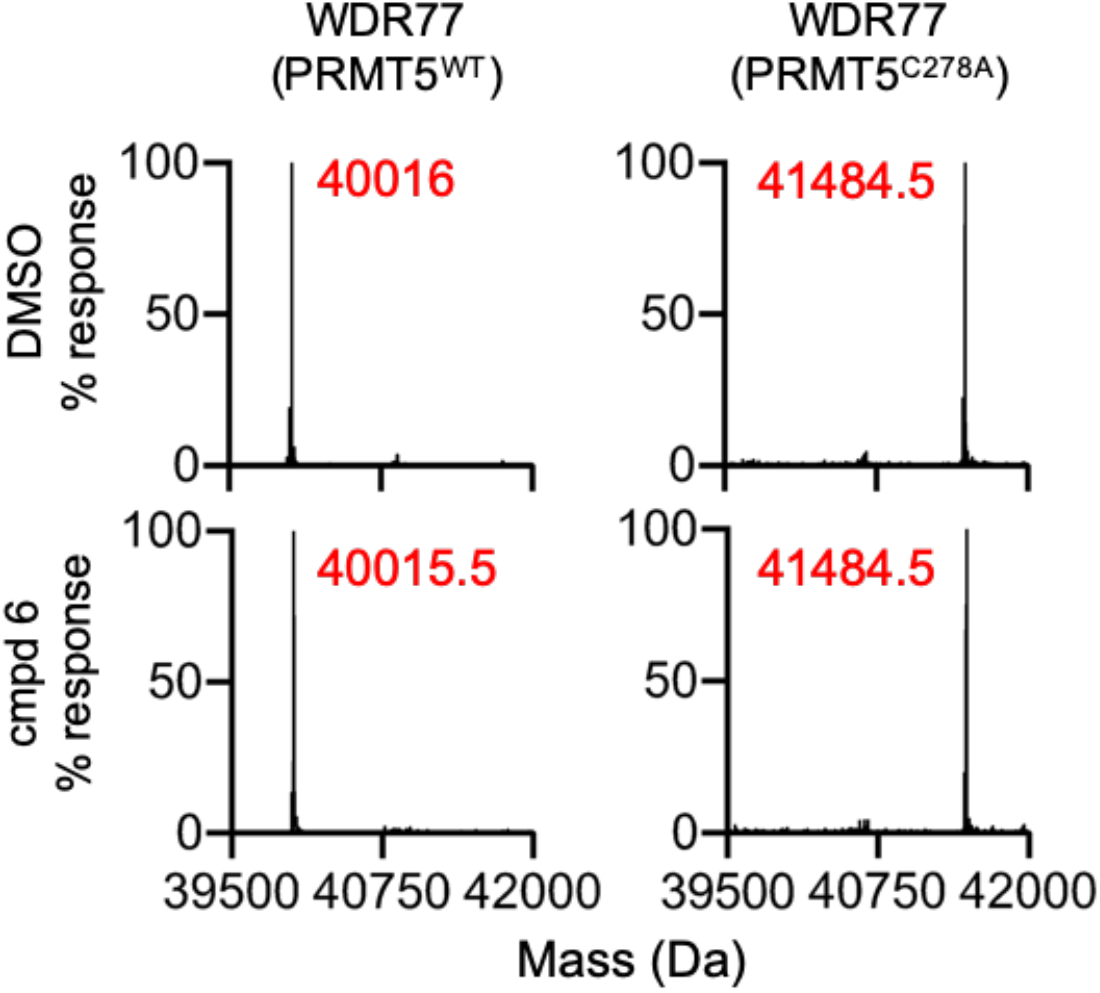
WDR77 is unmodified by compound 6. WDR77 is unmodified by compound 6. Deconvoluted mass spectra of both WDR77 and PRMT5 in the PRMT5^WT^ and PRMT5^C278A^ complexes, +/- compound 6. The same data for the PRMT5 protein is represented in Fig. 3c. Table shows the theoretical molecular weights and measured masses for both WDR77 and PRMT5 proteins in the PRMT5^WT^ and PRMT5^C278A^ complexes. All theoretical masses account for removal of the N-terminal methionine residue and subsequent N-terminal acetylation. No compound modification is expected on any protein except for PRMT5^WT^. The difference in masses of WDR77 in the PRMT5^WT^ and PRMT5^C278A^ complexes is due to the difference in expression tags.

**Supplemental Figure 3.**
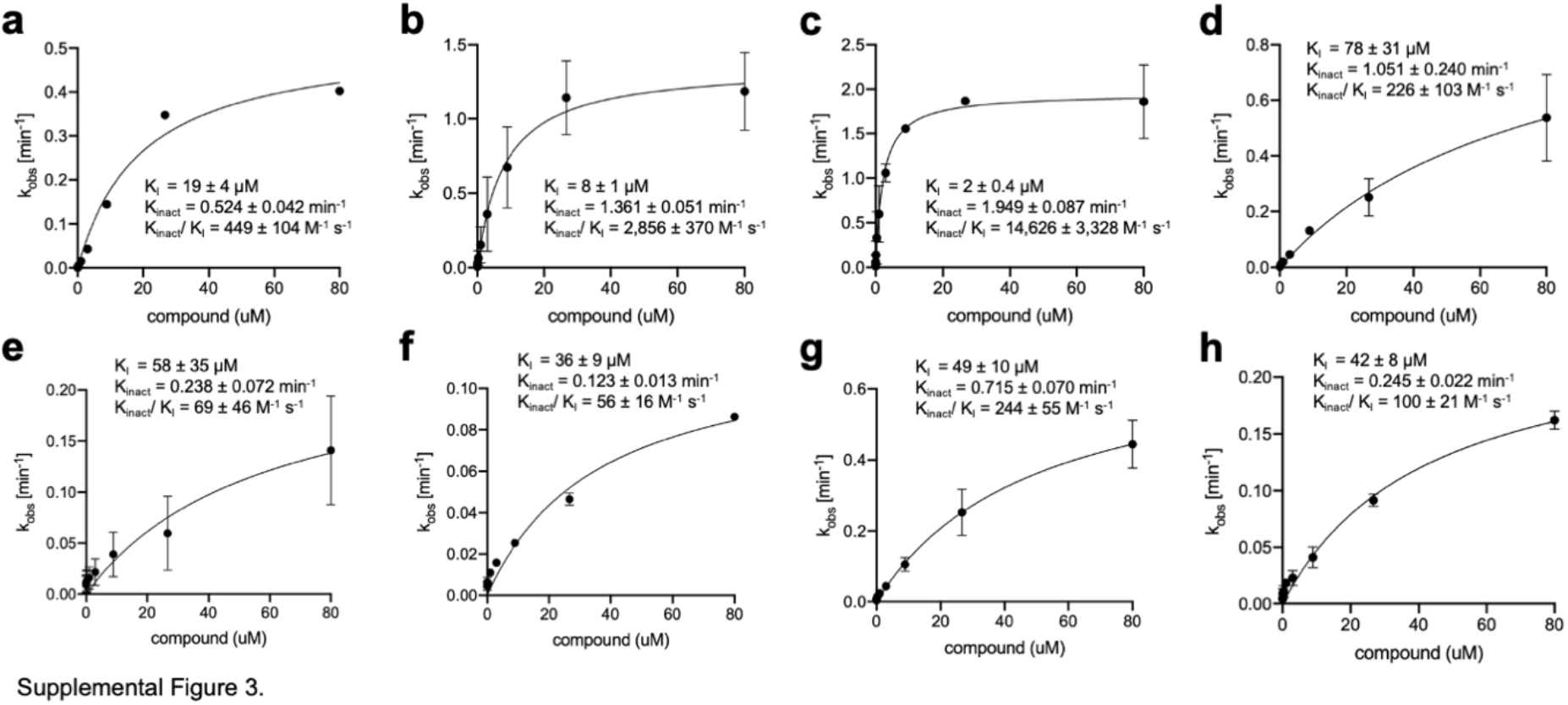
Kobs vs compound concentration for select compounds. a) Compound **1**, b) compound **6**, c) compound **20**, d) compound **21**, e) compound **25**, f) compound **26**, g) compound **BRD0639**, h) compound **28**.

#### General experimental conditions for small molecule synthesis

All anhydrous solvents, reagent grade solvents for chromatography and starting materials were purchased from either Sigma Aldrich Chemical Co. or Fisher Scientific. Water was distilled and purified through a Milli-Q water system (Millipore Corp., Bedford, MA). General methods for purification of compounds involved the use of silica cartridges purchased from Grace or Combiflash Purification systems. The reactions were monitored by TLC on precoated Merck 60 F254 silica gel plates and visualized using UV light (254 nm). All compounds were analyzed for purity by HPLC and characterized by 1H NMR using Bruker 400 MHz NMR spectrometers. Chemical shifts are reported in ppm (δ) relative to the residual solvent peak in the corresponding spectra (chloroform δ 7.26, methanol δ 3.31, DMSO δ 3.33) and coupling constants (J) are reported in hertz (Hz) (where s = singlet, bs = broad singlet, d = doublet, dd = double doublet, bd = broad doublet, ddd = double doublet of doublet, t = triplet, tt = triple triplet, q = quartet, m = multiplet) and analyzed using ACD NMR or MestReNova data processing. Mass spectra values are reported as m/z. All reactions were conducted under nitrogen unless otherwise noted. Solvents were removed in vacuo on a rotary evaporator. All final compounds for biological testing were with ? 95% purity [Shimadzu HPLC instrument with a Hamilton reversed phase column (HxSil, C18, 3μm, 2.1 mm × 50 mm (H2)). Eluent A: 5% CH3CN in H2O, eluent B: 90% CH3CN in H2O. A flow rate of 0.2 mL/min was used with UV detection at 254 and 214 nm].

#### Analytical UPLC methods

LRMS (LC-MS) Instrument: Agilent 1200\G1956A; Column: Kinetex EVO C18 30*2.1mm,5um; eluent A: 0.0375% TFA in water (v/v); eluent B: 0.01875% TFA in Acetonitrile (v/v); gradient 0-0.8 min5-95% B, 0.8-1.2 min 95% B, 1.2-1.5 5%B; flow 1.5 mL/min; temperature 50°C; DAD scan: 200-500 nm.

HRMS (LC-MS) was performed on purified compounds, diluted to 0.1 mM in DMSO, injecting 3 μl, reported data is an average of triplicate runs. Instrument: Agilent 1290 UHPLC; Column: Waters Acquity UPLC BEH C18, 1.7 μm, 2.1×50 mm; eluent A: Water with 0.1% formic acid, B: Acetonitrile with 0.1% formic acid; gradient 0-0.1 min 5% B, 0.1-5.0 min 5-95% B, 5.0-5.5 min 95% B, 5.5-5.6 min 5% B, 5.6-6.0 min 5% B; flow 0.5 mL/min; temperature 45°C; DAD scan: 200-500 nm, MS System Agilent 6545 Quadrupole Time of Flight Source Parameters: Gas Temp (°C) 350, Drying Gas (l/min) 10, Nebulizer (psi) 20, Sheath gas Temp (°C) 400, Sheath Gas Flow (l/min) 12, Capillary Voltage (V) 3500, Nozzle Voltage (V) 2000.

## Abbreviations

DCM: dichloromethane
DEA: diethyl amine
DIEA or DIPEA: N,N-diethyl isopropylamine
DIAD: diisopropyl azodicarboxylate
DMF: N,N-dimethyl formamide
HATU - 1: [Bis(dimethylamino)methylene]-1H-1,2,3-triazolo[4,5-b]pyridinium 3-oxide hexafluorophosphate
IPA: isopropyl alcohol (2-propanol)
LCMS: liquid chromatography, mass spectrometry
Pd_2_(dba)_3_: Tris(dibenzylideneacetone)dipalladium
Pd(Dppf): [1,1’-Bis(diphenylphosphino)ferrocene]dichloropalladium(II) dichloride, complex with dichloromethane
prepTLC: Preparative thin-layer chromatography
PRMT5: Protein Arginine Methyltransferase 5
Psi or PSI: pounds per square inch, gauge
SAR: structure activity relationship
T3P: 2,4,6-Tripropyl-1,3,5,2λ5,4λ5,6λ5-trioxatriphosphinane 2,4,6-trioxide
TEA: triethylamine
TFA: trifluoroacetic acid
THF: tetrahydrofuran
TLC: thin layer chromatography
Xantphos: 4,5-Bis(diphenylphosphino)-9,9-dimethylxanthene

---

**Supplemental Scheme 1.**
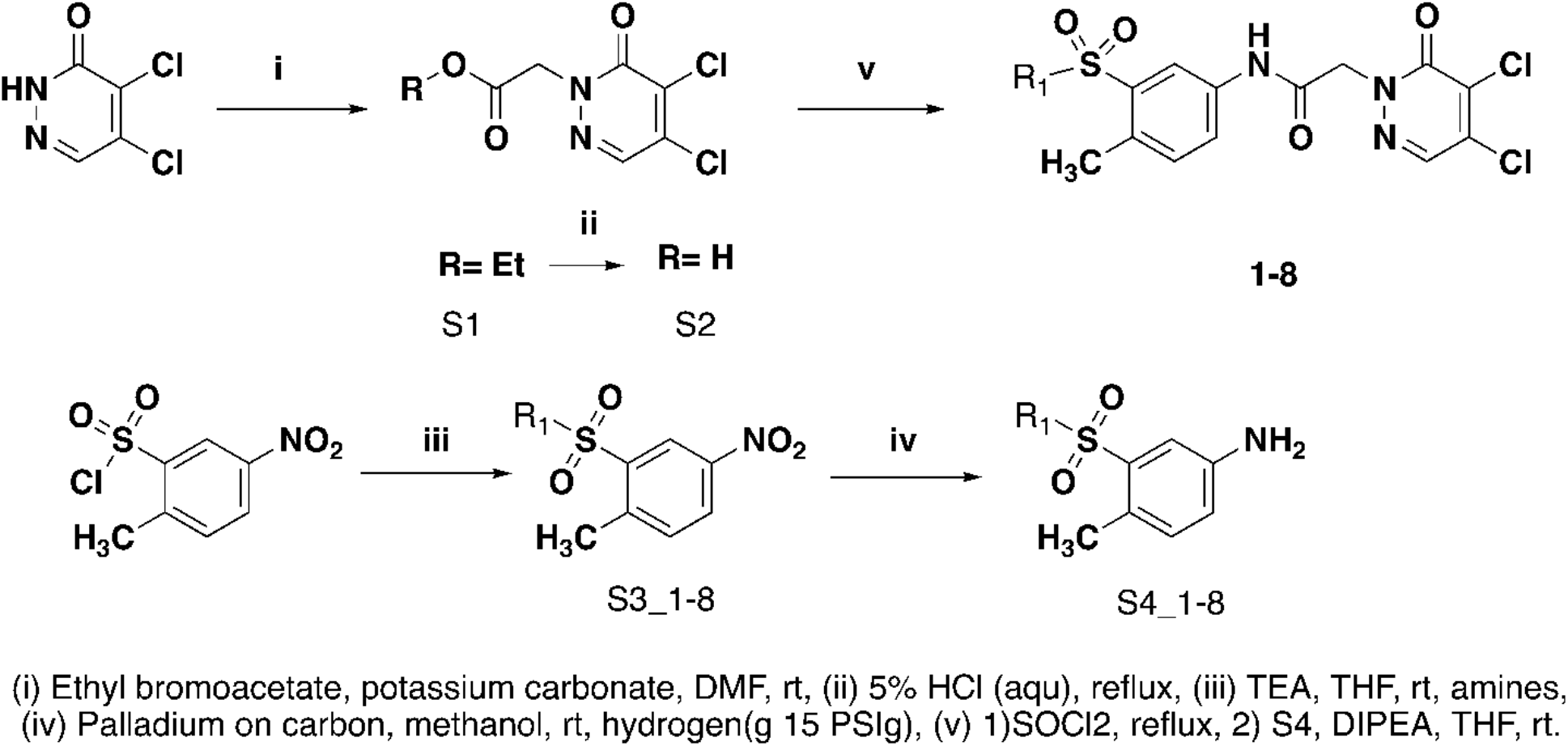
General synthetic scheme describing the synthesis of compounds.

ethyl 2-(4,5-dichloro-6-oxopyridazin-1(6H)-yl)acetate (S1)

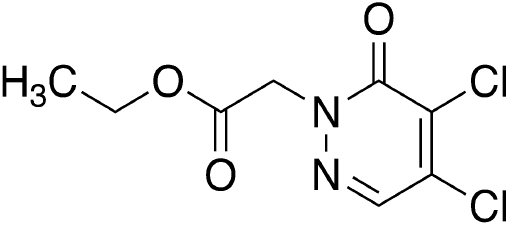

To a mixture of 4,5-dichloropyridazin-3(2H)-one (CAS 932-22-9, 30 g, 181.84 mmol) and K_2_CO_3_ (50.27 g, 363.69 mmol) in DMF (150 mL) was added ethyl 2-bromoacetate (33.40 g, 200.03 mmol, 22.12 mL) at 25 °C, and the mixture was stirred at 50 °C for 2 hours. The reaction mixture with diluted with water (100 ml) and ethyl acetate (300 ml), and the layers separated. The organic layer was washed with saturated aqueous sodium chloride (200 mL), dried over anhydrous sodium sulfate, insoluble materials removed by filtration, and volatiles removed under reduced pressure, the resulting residue was purified by column chromatography on silica gel eluting with a gradient of ethyl acetate in petroleum ether to give the title compound as a white solid (38.4 g, 84% yield). LRMS (m/z): Calcd [M+H]^+^ for C_8_H_10_Cl_2_N_2_O_3_ 260.0; found 250.9; ^1^H-NMR (400 MHz, CHLOROFORM-d) δ = 7.82 (s, 1H), 4.90 (s, 2H), 4.28 (q, 2H), 1.30 (t, 3H).

2-(4,5-dichloro-6-oxopyridazin-1(6H)-yl)acetic acid (S2)

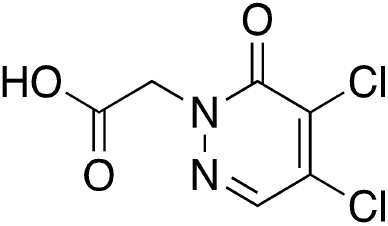

Ethyl 2-(4,5-dιchloro-6-oxopyrιdazιn-1(6H)-yl)acetate (S1, 15 g, 59.75 mmol) was suspended in aqueous hydrochloric acid (5% w/v, 300 mL) and the mixture was stirred at 105 °C for 90 minutes, cooled to room temperature, and the resulting solid was isolated by filtration to give the title compound (10 g, 75% yield) as a white solid. LRMS (m/z): Calcd [M+H]^+^ for C_6_H_5_Cl_2_N_2_O_3_ 223.0; found 223.0; ^1^H-NMR (400 MHz, CHLOROFORM-d) δ = 7.85 (s, 1H), 4.95 (s, 2H).

1-((2-methyl-5-nitrophenyl)sulfonyl)azepane (S3_1)

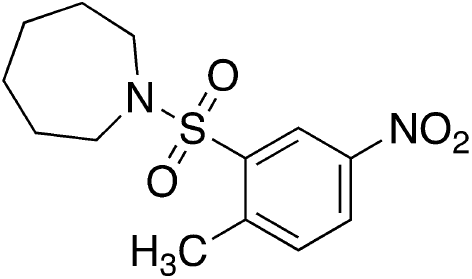

To a solution of 2-methyl-5-nitrobenzene-1-sulfonyl chloride (5 g, 21.22 mmol) and azepane (4.32 g, 31.83 mmol, 4.91 mL) in THF (100 mL) at 0 °C was slowly added TEA (6.44 g, 63.66 mmol, 8.86 mL). The mixture was stirred for 12 h at 25°C, volatiles removed under reduced pressure, and the residue partitioned between ethyl acetate and water, the layers were separated and the aqueous phase was extracted twice more with ethyl acetate, the combined organic layers were washed with saturated aqueous sodium chloride, dried over anhydrous sodium sulfate, insoluble materials removed by filtration, volatiles removed under reduced pressure to provide the title compound as a yellow oil (3.7 g), which was used without further manipulation. LRMS (m/z): Calcd [M+H]^+^ for C_13_H_19_N_2_O_4_S 289.1; found 299.0; ^1^H NMR (400 MHz, CHLOROFORM-d) δ = 8.60 (d, J = 2.4 Hz, 1H), 8.27 (dd, J = 2.4, 8.4 Hz, 1H), 7.51 (d, J = 8.3 Hz, 1H), 3.47 - 3.40 (m, 4H), 2.75 (s, 3H), 1.88 - 1.78 (m, 4H), 1.74 - 1.65 (m, 4H).

3-(azepan-1-ylsulfonyl)-4-methylaniline (S4_1)

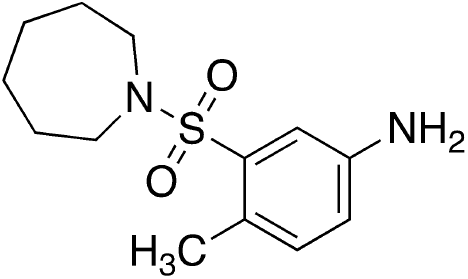

A mixture of 1-((2-methyl-5-nitrophenyl)sulfonyl)azepane (S3_1, 3.7 g, 12.40 mmol) and palladium on carbon (100 mg, 10% w/w) in methanol (80 mL) was degassed and purged with hydrogen three times, and the mixture was stirred at 25 °C for 16 hours under hydrogen atmosphere (15 psi). The mixture was filtered through Celite, and volatiles removed under reduced pressure to afford the title compound as a yellow oil (3.4 g). LRMS (m/z): Calcd [M+H]^+^ for C_13_H_21_N_2_O_2_S 269.1; found 269.0.

N-(3-(azepan-1-ylsulfonyl)-4-methylphenyl)-2-(4,5-dichloro-6-oxopyridazin-1(6H)-yl)acetamide (1)

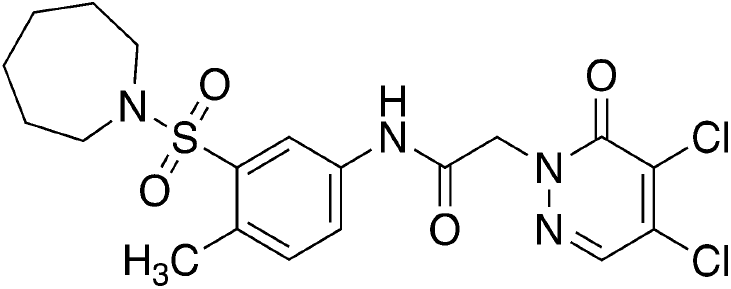

A mixture of 2-(4,5-dichloro-6-oxopyridazin-1(6H)-yl)acetic acid (S2, 997.17 mg, 4.47 mmol) and DMF (27.24 mg, 372.61 umol, 28.67 uL) and thionyl chloride (664.95 mg, 5.59 mmol) in THF (20 mL) at 0 °C was heated at 60 °C for 30 minutes. The solution was cooled to 0 °C, treated with 3-(azepan-1-ylsulfonyl)-4-methylaniline (Intermediate 20, 1 g, 3.73 mmol) and TEA (754.10 mg, 7.45 mmol, 1.04 mL) and warmed to 25 °C and stirred for 30 minutes. The reaction mixture was diluted with saturated aqueous ammonium chloride, extracted twice with ethyl acetate, combined organic layers were dried over sodium sulfate, insoluble materials were removed by filtration, volatiles were removed under reduced pressure and the residue was purified by prep-HPLC (column: Kromasil 250*50 mm*10 um; mobile phase: [water (0.225% FA)-ACN]; B%: 37ACN%-67ACN%, 26 min, 85% min) to give the title compound as a white solid (1.4 g, 79% yield). LCMS: Rt=0.805 min, [M+H]^+^= 418.2; ^1^H NMR (400 MHz, DMSO) δ 10.60 (s, 1H), 8.29 (s, 1H), 8.06 (d, J = 2.3 Hz, 1H), 7.66 (dd, J = 8.3, 2.3 Hz, 1H), 7.38 (d, J = 8.3 Hz, 1H), 4.98 (s, 2H), 3.27 (t, J = 5.8 Hz, 4H), 2.47 (s, 3H), 1.65 (tt, J = 6.4, 3.8 Hz, 4H), 1.56 (dq, J = 5.8, 4.2, 2.7 Hz, 4H); ^13^C{^1^H} NMR (101 MHz, DMSO) δ 164.70, 155.89, 138.32, 136.64, 136.54, 136.11, 133.33, 132.85, 131.26, 122.50, 118.77, 55.79, 47.63, 28.91, 26.35, 19.26. HRMS (ESI/Q-TOF) RT 3.35 min, m/z: [M + H]+ Calcd for C_19_H_22_Cl_2_N_4_O_4_S 473.0817; Found 473.0371.

Compounds 2, 4-14 were made analogously to compound 1

2-(4,5-dichloro-6-oxopyridazin-1(6H)-yl)-N-(4-methyl-3-((4-methyl-1,4-diazepan-1-yl)sulfonyl)phenyl)acetamide (2)

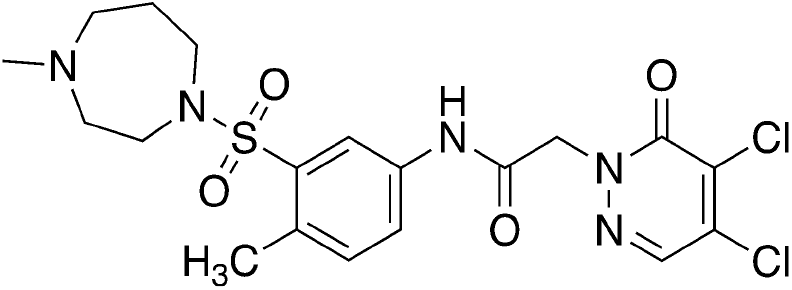

LCMS: Rt=0.742 min, [M+H]^+^= 488.1; ^1^H NMR (400 MHz, DMSO-d6) δ = 10.66 - 10.53 (m, 1H), 8.33 - 8.25 (m, 1H), 8.12 - 8.01 (m, 1H), 7.77 - 7.62 (m, 1H), 7.49 - 7.29 (m, 1H), 5.12 - 4.91 (m, 2H), 3.34 - 3.30 (m, 2H), 2.60 - 2.52 (m, 6H), 2.49 - 2.44 (m, 3H), 2.29 - 2.20 (m, 3H), 1.83 - 1.75 (m, 2H); HRMS (ESI/Q-TOF) RT: 2.01 min, m/z: [M + H]+ Calcd for C_19_H_24_Cl_2_N_5_O_4_S 488.0926; Found 488.0919

2-(4,5-dichloro-6-oxopyridazin-1(6h)-yl)-n-(4-methyl-3-(piperidin-1-ylsulfonyl)phenyl)acetamide (3)

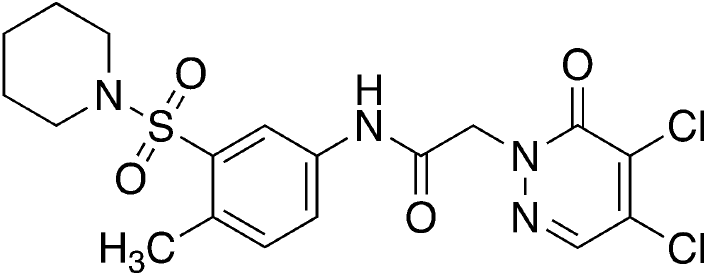

purchased from Enamine and used without further purification. HRMS (ESI/Q-TOF) RT: 3.22 min, m/z: [M + H]+ Calcd for C_18_H_21_Cl_2_N_4_O_4_S 459.0660; Found 459.0205.

2-(4,5-dichloro-6-oxopyridazin-1(6H)-yl)-N-(4-methyl-3-((4-methylpiperazin-1-yl)sulfonyl)phenyl)acetamide (4)

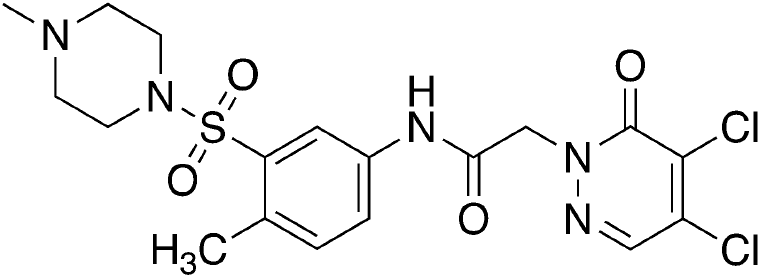

LCMS: Rt=0.753 min, [M+H]^+^= 474.0; ^1^H NMR (400 MHz, DMSO-d6) δ = 10.66 (s, 1H), 8.30 (s, 1H), 8.11 (d, J = 2.1 Hz, 1H), 7.69 (br d, J = 2.1 Hz, 1H), 7.42 (d, J = 8.3 Hz, 1H), 4.99 (s, 2H), 3.06 (br s, 4H), 2.53 - 2.52 (m, 3H), 2.35 (br s, 4H), 2.16 (s, 3H). HRMS (ESI/Q-TOF) RT: 1.97 min, m/z: [M + H]+ Calcd for C_18_H_22_Cl_2_N_5_O_4_S 474.0769; Found 474.0763.

2-(4,5-dichloro-6-oxopyridazin-1(6H)-yl)-N-(3-(N,N-dimethylsulfamoyl)-4-methylphenyl)acetamide (5)

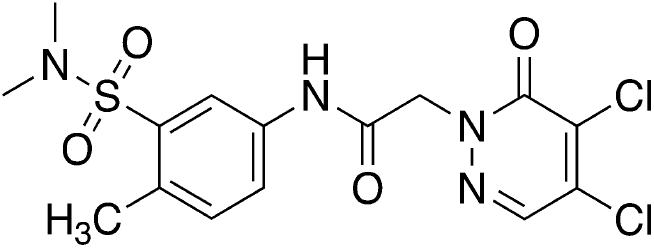

LCMS (ESI): Rt=0.877 min, [M+H]^+^= 418.9; ^1^H NMR (400 MHz, DMSO-d_6_) δ = 10.67 - 10.62 (m, 1H), 8.30 (s, 1H), 8.09 (d, J = 2.1 Hz, 1H), 7.70 (dd, J = 2.3, 8.3 Hz, 1H), 7.42 (s, 1H), 4.99 (s, 2H), 2.73 (s, 6H), 2.54 - 2.53 (m, 3H); HRMS (ESI/Q-TOF) RT: 2.72 min, m/z: [M + H]+ Calcd for C15H17Cl2N4O4S 419.0347; Found 419.0345.

2-(4,5-Dichloro-6-oxopyridazin-1(6H)-yl)-N-(4-methyl-3-(N-(2-(pyridin-2-yl)ethyl)sulfamoyl)phenyl)acetamide (6)

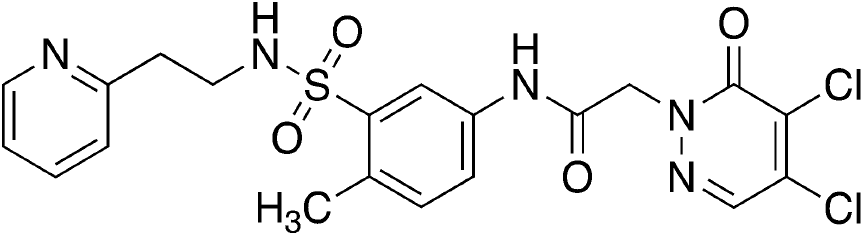

LCMS: Rt=0.732 min, [M+H]^+^= 496.1; ^1^H NMR (400 MHz, DMSO) δ 10.58 (s, 1H), 8.40 (ddd, J = 4.8, 1.9, 0.9 Hz, 1H), 8.29 (s, 1H), 8.09 (d, J = 2.3 Hz, 1H), 7.77 (t, J = 5.8 Hz, 1H), 7.69 (dd, J = 8.2, 2.3 Hz, 1H), 7.64 (td, J = 7.7, 1.9 Hz, 1H), 7.31 (d, J = 8.4 Hz, 1H), 7.17 (ddd, J = 7.6, 4.8, 1.2 Hz, 1H), 7.13 (dt, J = 7.8, 1.1 Hz, 1H), 4.97 (s, 2H), 3.19 – 3.08 (m, 2H), 2.81 (t, J = 7.3 Hz, 2H), 2.43 (s, 3H); ^13^C{^1^H} NMR (101 MHz, DMSO) δ 164.61, 158.23, 155.90, 148.97, 138.72, 136.50, 136.45, 136.40, 136.08, 132.99, 132.85, 131.22, 123.20, 122.55, 121.58, 119.29, 55.76, 41.96, 37.28, 19.0;. HRMS (ESI/Q-TOF) RT: 1.95 min, m/z: [M + H]+ Calcd for C20H20Cl2N5O4S 496.0613; Found 496.0606.

2-(4,5-Dichloro-6-oxopyridazin-1(6H)-yl)-N-(4-methyl-3-(N-(2-(pyridin-3-yl)ethyl)sulfamoyl)phenyl)acetamide (7)

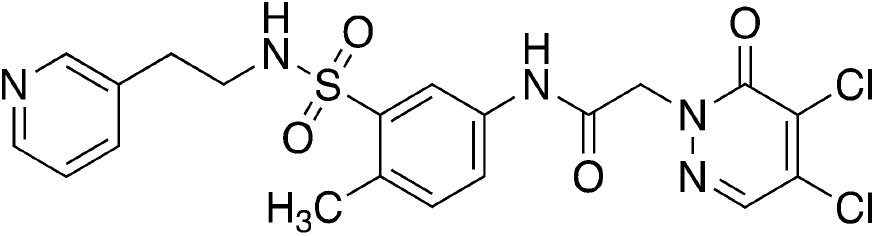

LCMS: Rt=0.732 min, [M+H]^+^= 496.1;^1^H NMR (400 MHz, DMSO-d6) δ = 10.62 (s, 1H), 8.38 (br d, J = 3.6 Hz, 1H), 8.32 (s, 1H), 8.29 (s, 1H), 8.12 (d, J = 1.9 Hz, 1H), 7.80 (br t, J = 5.3 Hz, 1H), 7.66 (dd, J = 1.9, 8.2 Hz, 1H), 7.51 (br d, J = 7.8 Hz, 1H), 7.30 (d, J = 8.3 Hz, 1H), 7.24 (dd, J = 4.8, 7.6 Hz, 1H), 4.98 (s, 2H), 3.03 (q, J = 6.5 Hz, 2H), 2.68 (br t, J = 6.9 Hz, 2H), 2.42 (s, 3H); HRMS (ESI/Q-TOF) RT: 1.93 min, m/z: [M + H]+ Calcd for C20H20Cl2N5O4S 496.0613; Found 496.0606.

2-(4,5-Dichloro-6-oxopyridazin-1(6H)-yl)-N-(4-methyl-3-(N-(2-(pyridin-4-yl)ethyl)sulfamoyl)phenyl)acetamide (8)

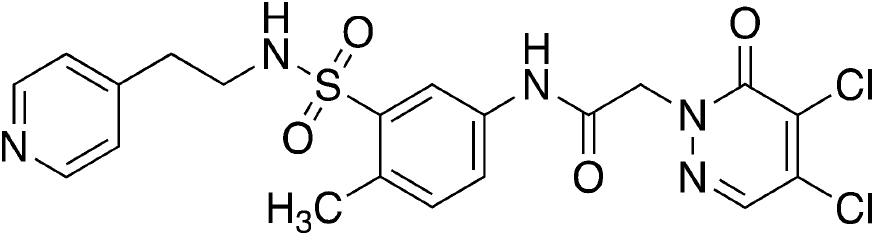

LCMS: Rt=0.752 min, [M+H]^+^= 496.0; ^1^H NMR (400 MHz, DMSO-d6) δ = 10.58 (s, 1H), 8.40 (br d, J = 5.0 Hz, 2H), 8.29 (s, 1H), 8.13 (d, J = 2.1 Hz, 1H), 7.79 (t, J = 5.7 Hz, 1H), 7.65 (dd, J = 2.2, 8.2 Hz, 1H), 7.31 (d, J = 8.3 Hz, 1H), 7.11 (d, J = 5.8 Hz, 2H), 4.98 (s, 2H), 3.08 - 3.03 (m, 2H), 2.70 - 2.66 (m, 2H), 2.41 (s, 3H); HRMS (ESI/Q-TOF) RT: 1.91 min, m/z: [M + H]+ Calcd for C_20_H_20_Cl_2_N_5_O_4_S 496.0613; Found 496.0606.

2-(4,5-dichloro-6-oxo-pyridazin-1-yl)-N-[3-[2-(2-pyridyl)ethylsulfamoyl]phenyl]acetamide (9)

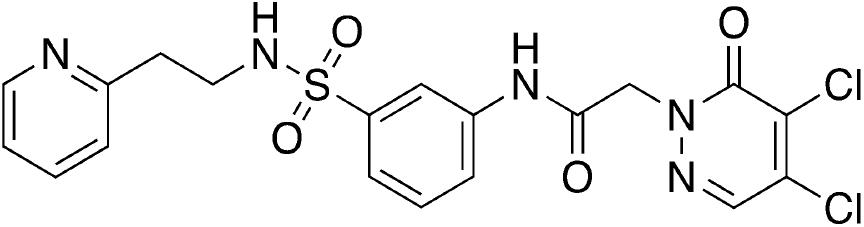

^1^H NMR (400 MHz, Chloroform-d) δ 8.96 (s, 1H), 8.52 (d, J = 5.1 Hz, 1H), 8.14 (s, 1H), 7.95 (dd, J = 8.3, 1.8 Hz, 1H), 7.92 (s, 1H), 7.79 (d, J = 2.3 Hz, 1H), 7.71 (td, J = 7.7, 1.7 Hz, 1H), 7.54 (d, J = 7.7 Hz, 1H), 7.38 (t, J = 8.0 Hz, 1H), 7.29 (s, 1H), 7.21 (d, J = 7.8 Hz, 1H), 5.06 (s, 2H), 3.41 (t, J = 6.2 Hz, 2H), 3.04 (t, J = 6.3 Hz, 2H);HRMS (ESI/Q-TOF) RT: 1.84 min, m/z: [M + H]+ Calcd for C_19_H_18_Cl_2_N_5_O_4_S 482.0456; Found 482.0447.

2-(4,5-dichloro-6-oxopyridazin-1(6h)-yl)-n-(4-methoxy-3-(n-(2-(pyridin-2-yl)ethyl) sulfamoyl)phenyl)acetamide (10)

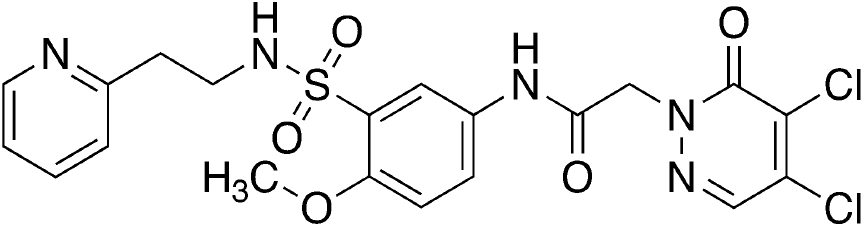

^1^H NMR (400 MHz, DMSO-d6) δ = 10.45 (s, 1H), 8.45 (d, J = 3.9 Hz, 1H), 8.29 (s, 1H), 8.00 (d, J = 2.6 Hz, 1H), 7.77 (dd, J = 2.6, 9.0 Hz, 1H), 7.67 (dt, J = 1.8, 7.6 Hz, 1H), 7.31 (t, J = 5.8 Hz, 1H), 7.23 - 7.19 (m, 1H), 7.18 (d, J = 2.1 Hz, 1H), 7.17 - 7.14 (m, 1H), 4.95 (s, 2H), 3.82 (s, 3H), 3.17 - 3.11 (m, 2H), 2.83 (t, J = 7.2 Hz, 2H); HRMS (ESI/Q-TOF) RT: 1.79 min, m/z: [M + H]+ Calcd for C_20_H_20_Cl_2_N_5_O_5_S 512.0562; Found 512.0556.

2-(4,5-Dichloro-6-oxopyridazin-1(6H)-yl)-N-(4-ethyl-3-(N-(2-(pyridin-2-yl)ethyl)sulfamoyl)phenyl)acetamide (11)

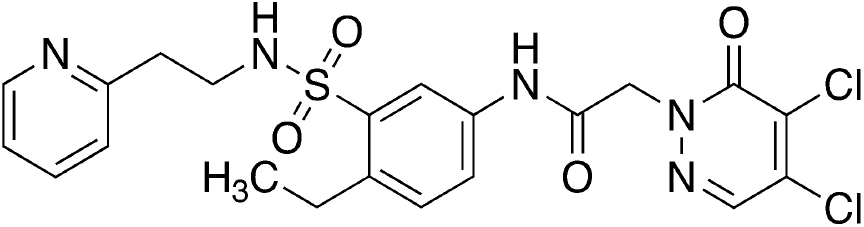

^1^H NMR (400 MHz, DMSO-d6) δ = 10.60 (s, 1H), 8.42 (br d, J = 4.3 Hz, 1H), 8.29 (s, 1H), 8.09 (d, J = 1.8 Hz, 1H), 7.82 (br t, J = 5.6 Hz, 1H), 7.74 (dd, J = 1.8, 8.3 Hz, 1H), 7.65 (dt, J = 1.5, 7.6 Hz, 1H), 7.38 (d, J = 8.3 Hz, 1H), 7.22 - 7.11 (m, 2H), 4.98 (s, 2H), 3.20 - 3.14 (m, 2H), 2.90 - 2.81 (m, 4H), 1.16 (t, J = 7.5 Hz, 3H); HRMS (ESI/Q-TOF) RT: 2.16 min, m/z: [M + H]+ Calcd for C_21_H_22_Cl_2_N_5_O_4_S 510.0769; Found 510.0760.

2-(4,5-dichloro-6-oxopyridazin-1(6h)-yl)-n-(2,4-dimethyl-5-(n-(2-(pyridin-2-yl)ethyl)sulfamoyl)phenyl)acetamide (12)

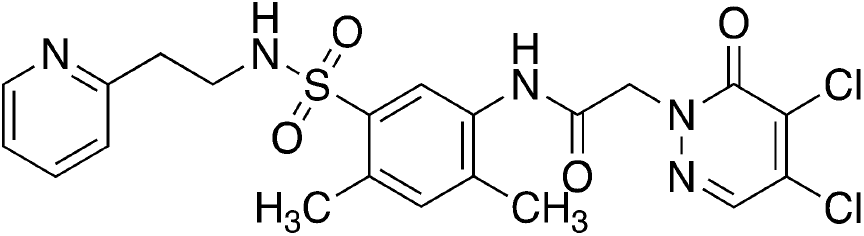

^1^H NMR (400 MHz, DMSO-d6) δ = 9.82 (s, 1H), 8.40 (br d, J = 4.0 Hz, 1H), 8.29 (s, 1H), 7.90 (s, 1H), 7.69 (br t, J = 5.6 Hz, 1H), 7.63 (dt, J = 1.5, 7.6 Hz, 1H), 7.22 (s, 1H), 7.20 - 7.15 (m, 1H), 7.13 (d, J = 7.8 Hz, 1H), 5.03 (s, 2H), 3.12 (q, J = 6.9 Hz, 2H), 2.81 (t, J = 7.3 Hz, 2H), 2.43 (s, 3H), 2.25 (s, 3H); HRMS (ESI/Q-TOF) RT: 1.96 min, m/z: [M + H]+ Calcd for C_21_H_22_Cl_2_N_5_O_4_S 510.0769; Found 510.0762.

2-(4,5-dichloro-6-oxopyridazin-1(6h)-yl)-n-(3,4-dimethyl-5-(n-(2-(pyridin-2-yl)ethyl)sulfamoyl)phenyl)acetamide (13)

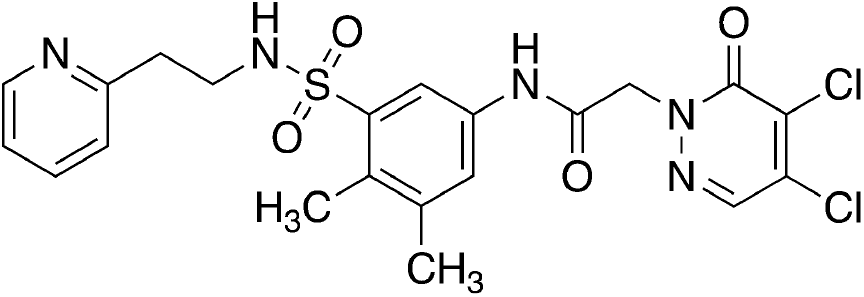

^1^H NMR (400 MHz, DMSO-d_6_) δ = 10.49 (s, 1H), 8.43 - 8.36 (m, 1H), 8.29 (s, 1H), 7.97 (d, J = 1.7 Hz, 1H), 7.76 (t, J = 5.7 Hz, 1H), 7.66 - 7.61 (m, 2H), 7.17 (dd, J = 5.1, 6.9 Hz, 1H), 7.11 (d, J = 7.8 Hz, 1H), 4.97 (s, 2H), 3.17 (q, J = 6.9 Hz, 2H), 2.82 (t, J = 7.2 Hz, 2H), 2.33 - 2.24 (m, 6H);HRMS (ESI/Q-TOF) RT: 2.08 min, m/z: [M + H]+ Calcd for C_21_H_22_Cl_2_N_5_O_4_S S 510.0769; Found 510.0762.

2-(4,5-Dichloro-6-oxopyridazin-1(6H)-yl)-N-(2,4-dimethyl-3-(N-(2-(pyridin-2-yl)ethyl)sulfamoyl)phenyl)acetamide (14)

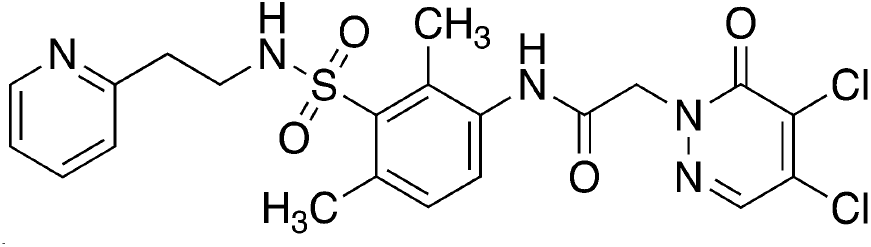

^1^H NMR (400 MHz, DMSO-d_6_) δ = 9.85 (s, 1H), 8.39 (br d, J = 4.0 Hz, 1H), 8.29 (s, 1H), 7.70 (br t, J = 5.6 Hz, 1H), 7.61 (dt, J = 1.8, 7.6 Hz, 1H), 7.36 (br d, J = 8.1 Hz, 1H), 7.22 - 7.16 (m, 1H), 7.16 - 7.12 (m, 1H), 7.09 (br d, J = 7.7 Hz, 1H), 5.01 (s, 2H), 3.20 - 3.12 (m, 2H), 2.81 (br t, J = 7.2 Hz, 2H), 2.56 (s, 3H), 2.42 (s, 3H); HRMS (ESI/Q-TOF) RT: 1.89 min, m/z: [M + H]+ Calcd for C_21_H_22_Cl_2_N_5_O_4_S 510.0769; Found 510.0764.

**Supplemental Scheme 2.**
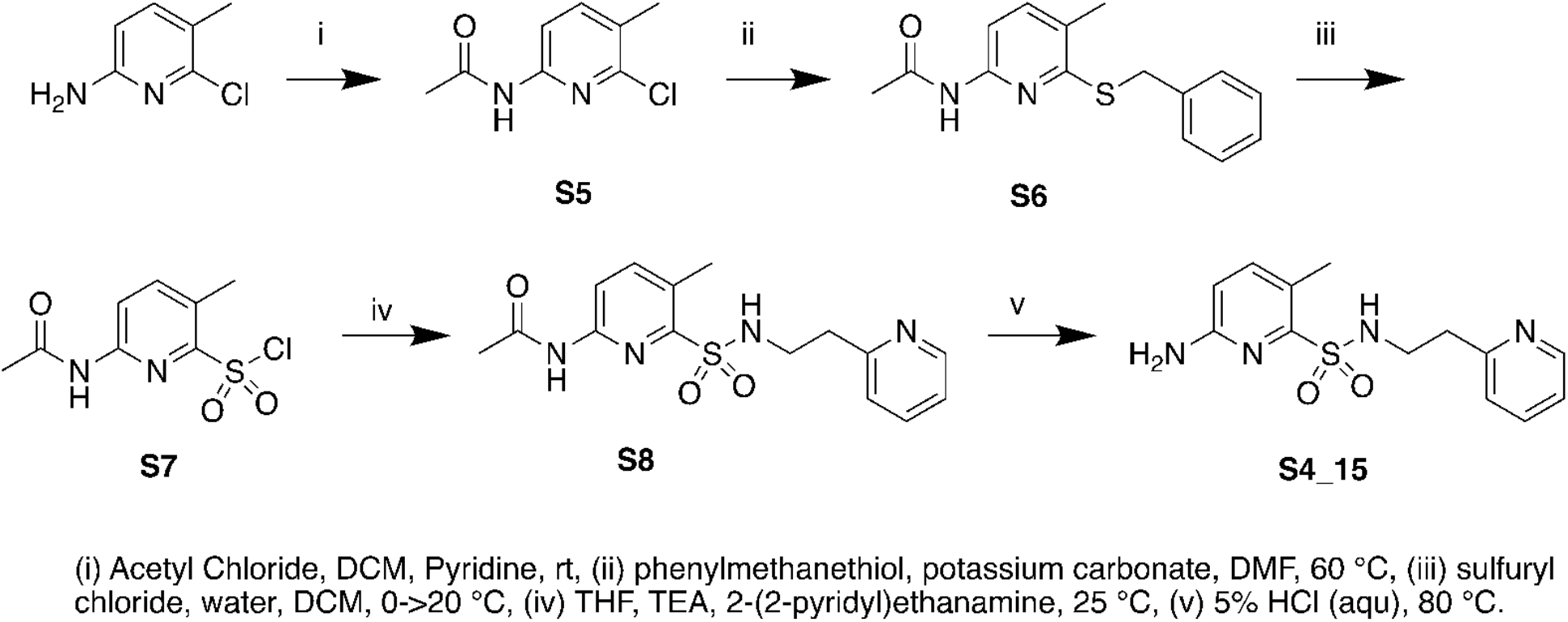
Synthesis of Compound 15

N-(6-chloro-5-methylpyridin-2-yl)acetamide (S5)

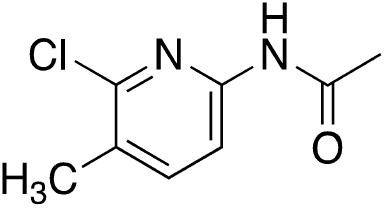

To a mixture of 6-chloro-5-methylpyπdm-2-amine (CAS 442129-37-5, 6 g, 42.08 mmol) and pyridine (4.99 g, 63.12 mmol, 5.09 mL) in DCM (50 mL) was added acetyl chloride (4.29 g, 54.70 mmol, 3.90 mL) and the mixture was stirred at 20 °C for 12 hr. The mixture was diluted with water and extracted twice with ethyl acetate, combined organic layers were washed with saturated aqueous sodium chloride, dried over sodium sulfate, filtered to remove insoluble materials, and volatiles removed to provide the title compound as a yellow solid (8.1 g). LRMS (m/z): Calcd [M+H]^+^ for C_8_H_10_ClN_2_O 185.4; found 185.4; ^1^H NMR (400 MHz, CHLOROFORM-d) δ = 8.12 - 8.02 (m, 1H), 8.02 - 7.83 (m, 1H), 7.61 - 7.50 (m, 1H), 2.36 - 2.32 (m, 3H), 2.20 - 2.16 (m, 3H).

N-(6-(benzylthio)-5-methylpyridin-2-yl)acetamide (S6)

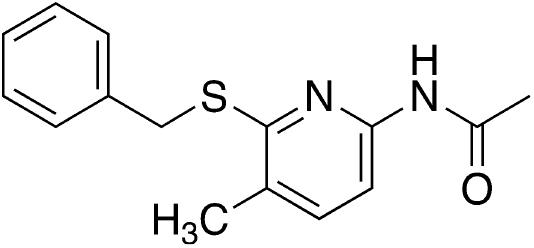

To a solution of N-(6-chloro-5-methylpyridin-2-yl)acetamide (S5, 10.1 g, 54.71 mmol) in DMF (100 mL) was added potassium carbonate (30.24 g, 218.82 mmol) and phenylmethanethiol (20.38 g, 164.12 mmol, 19.23 mL) at 20°C. The mixture was stirred at 60°C for 12 h, diluted with water and extracted twice with ethyl acetate, the combined organic layers were washed with water, saturated aqueous sodium chloride, dried over anhydrous sodium sulfate, insoluble materials removed by filtration, volatiles removed under reduced pressure and the residue was purified by column chromatography on silica gel eluting with a gradient of ethyl acetate in petroleum ether to afford the title compound as a yellow solid (9.33 g, 63% yield).^1^H NMR (400 MHz, CHLOROFORM-d) δ = 7.88 - 7.77 (m, 1H), 7.77 - 7.63 (m, 1H), 7.42 - 7.37 (m, 2H), 7.36 - 7.29 (m, 3H), 7.28 - 7.23 (m, 1H), 4.49 - 4.35 (m, 2H), 2.29 - 2.21 (m, 3H), 2.21 - 2.18 (m, 3H); ^1^H NMR (400 MHz, DMSO-d6) δ = 10.33 (s, 1H), 7.73 (br d, J = 8.1 Hz, 1H), 7.49 - 7.43 (m, 3H), 7.33 - 7.28 (m, 2H), 7.26 - 7.21 (m, 1H), 4.50 (s, 2H), 2.13 (s, 3H), 2.11 (s, 3H).

6-acetamido-3-methylpyridine-2-sulfonyl chloride (S7)

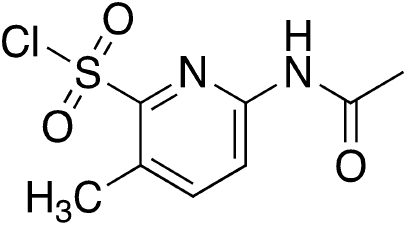

To a solution of N-(6-(benzylthio)-5-methylpyridin-2-yl)acetamide (S6, 9.33 g, 34.26 mmol) in DCM (100 mL) was added water (20 mL) and sulfuryl chloride (32.36 g, 239.79 mmol, 23.97 mL) at 0 °C. The reaction mixture slowly warmed to 20°C and stirred for 2 hr. Water was added and the mixture was extracted three times with dichloromethane, combined organic layers were washed with saturated aqueous sodium chloride, dried over sodium sulfate, insoluble materials removed by filtration, and volatiles removed under reduced pressure to afford the title compound as a yellow oil (11.6 g).

N-[5-methyl-6-[2-(2-pyridyl)ethylsulfamoyl]-2-pyridyl]acetamide (S8)

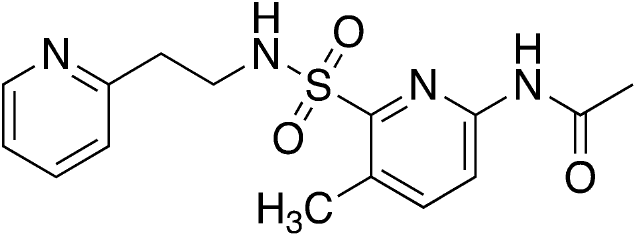

To a solution of 6-acetamido-3-methyl-pyridme-2-sulfonyl chloride (S7, 6 g, 24.13 mmol) in THF (60 mL) was added TEA (7.32 g, 72.38 mmol, 10.07 mL) and 2-(2-pyridyl)ethanamine (4.42 g, 36.19 mmol, 4.33 mL). The mixture was stirred at 25°C for 12 hr, diluted with water, and extracted twice with ethyl acetate. Combined organic layers were washed with saturated aqueous sodium chloride, dried over sodium sulfate, insoluble materials were removed by filtration, volatiles removed under reduced pressure and the residue purified by column chromatography on silica gel, eluting with a gradient of ethyl acetate in petroleum ether to provide the title compound as a yellow solid (4.7 g, 58% yield). LRMS (m/z): Calcd [M+H]^+^for C_15_H_19_N_4_O_3_S 335.1; found 335.3.

6-amino-3-methyl-N-[2-(2-pyridyl)ethyl]pyridine-2-sulfonamide (S4_15)

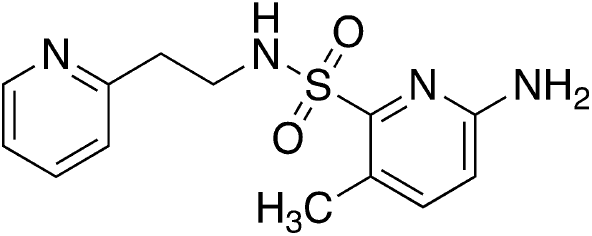

A mixture of N-[5-methyl-6-[2-(2-pyridyl)ethylsulfamoyl]-2-pyridyl]acetamide (S8, 4.7 g, 14.06 mmol, 1 eq) in hydrogen chloride (6M in water, 50 mL) was stirred at 80°C for 12 hr. Volatiles were removed under reduced pressure to provide the title compound as a white solid (4.6 g, HCl salt). ^1^H NMR (400 MHz, DMSO-d6) δ = 8.87 - 8.77 (m, 1H), 8.54 (dt, J = 1.5, 7.9 Hz, 1H), 8.11 (br s, 1H), 8.03 - 7.90 (m, 2H), 7.50 (d, J = 8.4 Hz, 1H), 6.73 (d, J = 8.6 Hz, 1H), 3.61 (q, J = 5.9 Hz, 2H), 3.30 (t, J = 6.5 Hz, 2H), 2.29 (s, 3H).

2-(4,5-dichloro-6-oxopyridazin-1(6H)-yl)-N-(5-methyl-6-(N-(2-(pyridin-2-yl)ethyl)sulfamoyl)pyridin-2-yl)acetamide (15)

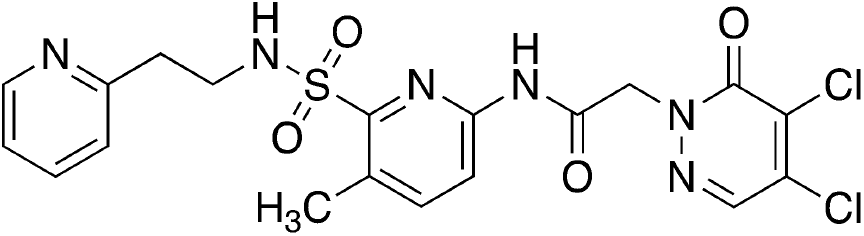

Prepared in a manner analogous Compound 6 using S4_15. 1H NMR (400 MHz, DMSO-d_6_) δ = 11.05 (s, 1H), 8.51 (d, J = 4.2 Hz, 1H), 8.09 (br d, J = 2.2 Hz, 1H), 7.97 (br t, J = 5.7 Hz, 1H), 7.89 (d, J = 8.3 Hz, 1H), 7.74 (dt, J = 1.8, 7.6 Hz, 1H), 7.31 - 7.23 (m, 2H), 5.16 (br s, 2H), 3.62 - 3.50 (m, 2H), 2.99 (t, J = 7.3 Hz, 2H), 2.53 (s, 3H); HRMS (ESI/Q-TOF) RT: 2.08 min, m/z: [M + H]+ Calcd for C_19_H_19_Cl_2_N_6_O_4_S 497.0565; Found 497.0558.

**Supplemental Scheme 3.**
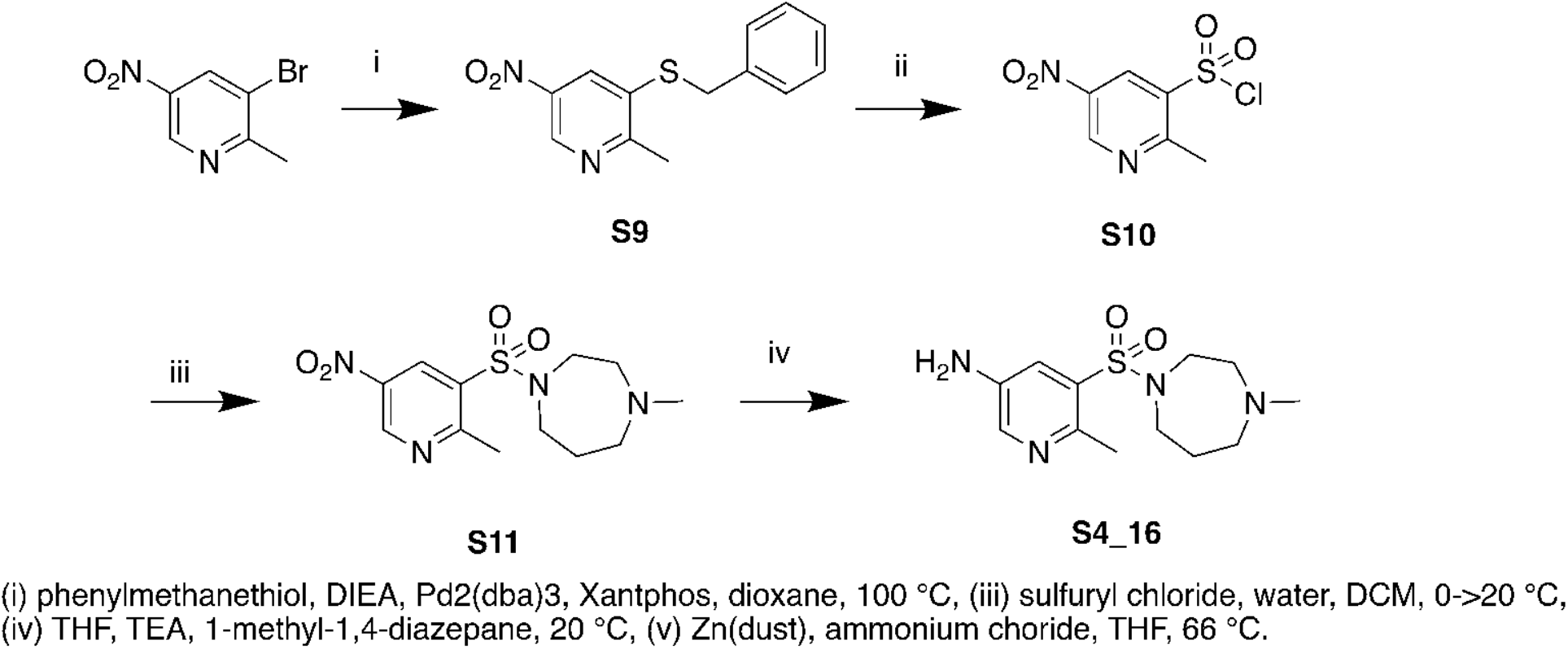
Synthesis of Compound 16.

3-(benzylthio)-2-methyl-5-nitropyridine (S9)

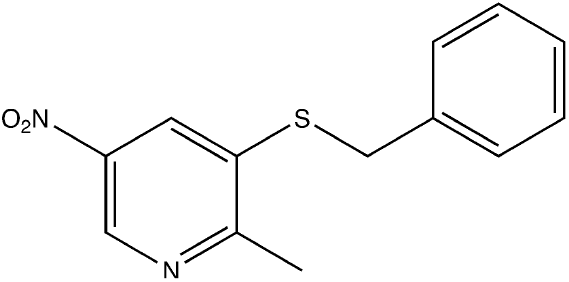

A mixture of 3-bromo-2-methyl-5-nitropyridine (2 g, 9.22 mmol) was added phenylmethanethiol (1.49 g, 11.98 mmol, 1.40 mL), DIEA (2.38 g, 18.43 mmol, 3.21 mL), Pd2(dba)3 (421.95 mg, 460.79 umol) and Xantphos (533.24 mg, 921.57 umol) in dioxane (30 mL) was heated at reflux (100°C) for 12 hours. The mixture was cooled to room temperature, diluted with water, extracted three times with ethyl acetate, the combined organic layers were washed with water, saturated aqueous sodium chloride, dried over sodium sulfate, insoluble materials removed by filtration, volatiles removed under reduced pressure and the residue purified by column chromatography on silica gel, eluting with a gradient of ethyl acetate in petroleum ether, to afford the title compound as a yellow solid (2.6g). 1H NMR (400 MHz, CHLOROFORM-d) δ = 9.04 - 8.91 (m, 1H), 8.25 - 8.04 (m, 1H), 7.33 - 7.22 (m, 5H), 4.23 - 4.06 (m, 2H), 2.62 - 2.53 (m, 3H).

2-methyl-5-nitropyridine-3-sulfonyl chloride (S10)

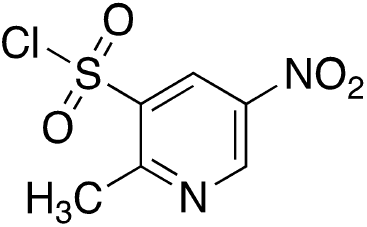

To 3-(benzylthio)-2-methyl-5-nitropyridine (S9, 1.0 g, 3.84 mmol) in DCM (20 mL) was added water (4 mL) and sulfuryl chloride (3.63 g, 26.89 mmol, 2.69 mL) at 0 °C. The reaction mixture slowly warmed to 20°C and stirred for 30 min, volatiles were removed under reduced pressure, the residue was suspended in dichloromethane, dried over sodium sulfate, insoluble materials removed by filtration, and volatiles removed under reduced pressure to afford the title compound as a yellow oil (1.0 g), which was used without further manipulation.

1-methyl-4-((2-methyl-5-nitropyridin-3-yl)sulfonyl)-1,4-diazepane (S11)

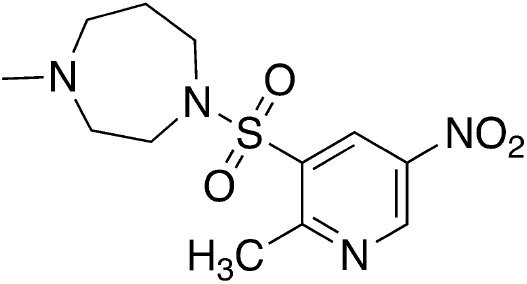

A mixture of 2-methyl-5-nitropyridine-3-sulfonyl chloride (S10, 1.0 g, 4.23 mmol) 1-methyl-1,4-diazepane (723.83 mg, 6.34 mmol, 788.49 uL) and TEA (1.28 g, 12.68 mmol, 1.76 mL) in THF (20 mL) was stirred for 12 h at 20°C. Volatiles were removed under reduced pressure and the residue was purified by prep-HPLC (column: Kromasil 250*50mm*10um; mobile phase: [water(0.225%FA)-ACN];B%: 5%-30%,15min) to afford the title compound as a brown solid (430 mg).

6-methyl-5-((4-methyl-1,4-diazepan-1-yl)sulfonyl)pyridin-3-amine (S4_16)

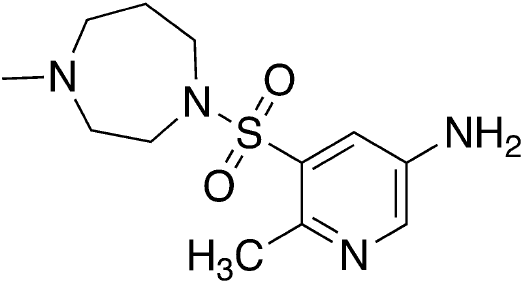

A mixture of 1-methyl-4-((2-methyl-5-nitropyridin-3-yl)sulfonyl)-1,4-diazepane (S11, 100 mg, 318.11 umol) ammonium chloride (170.16 mg, 3.18 mmol) and Zn dust (83.20 mg, 1.27 mmol) in THF (2 mL) was stirred at 66 °C for 12 hr. After cooling to room temperature, insoluble materials were removed by filtration through Celite, aqueous hydrochloric acid was added to the filtrate, and volatiles removed under reduced pressure to afford the title compound as a yellow solid (75 mg, di-HCl salt).

2-(4,5-dichloro-6-oxopyridazin-1(6H)-yl)-N-(6-methyl-5-((4-methyl-1,4-diazepan-1-yl)sulfonyl)pyridin-3-yl)acetamide (16)

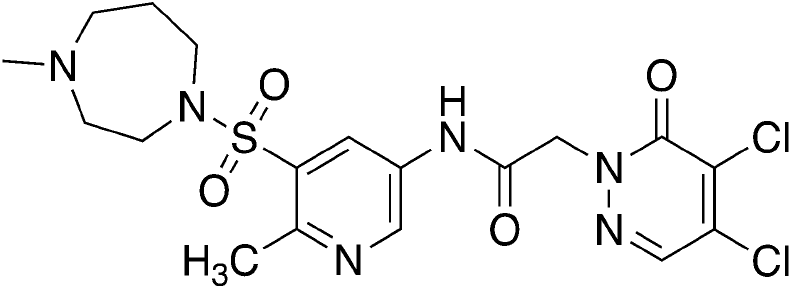

A mixture of 6-methyl-5-((4-methyl-1,4-diazepan-1-yl)sulfonyl)pyridin-3-amine (S4_16, 75 mg, 209.91 umol, 2 HCl salt), 2-(4,5-dichloro-6-oxo-pyridazin-1-yl)acetic acid (S2, 56.17 mg, 251.89 umol), 2-chloro-1-methyl-pyridin-1-ium iodide (80.44 mg, 314.86 umol), and DIEA (135.64 mg, 1.05 mmol, 182.81 uL, 5 eq) in THF (5 mL) was stirred at 20°C for 12 h. Volatiles were removed under reduced pressure and the residue was purified by prep-HPLC (column: Shim-pack C18 150*25*10um;mobile phase: [water(0.225%FA)-ACN];B%: 3%-33%,10min) to afford the title compound as a yellow solid (10 mg, 9% yield). ^1^H NMR (400 MHz, METHANOL-d4) δ = 8.75 - 8.70 (m, 1H), 8.65 - 8.60 (m, 1H), 8.14 - 8.10 (m, 1H), 5.12 - 5.06 (m, 2H), 3.74 - 3.67 (m, 2H), 3.58 - 3.51 (m, 2H), 3.24 - 3.18 (m, 4H), 2.80 - 2.77 (m, 3H), 2.76 - 2.72 (m, 3H), 2.20 - 2.07 (m, 2H); HRMS (ESI/Q-TOF) RT: 1.74 min, m/z: [M + H]+ Calcd for C_18_H_23_Cl_2_N_6_O_4_S 489.0878; Found 489.0872.

**Supplemental Scheme 4.**
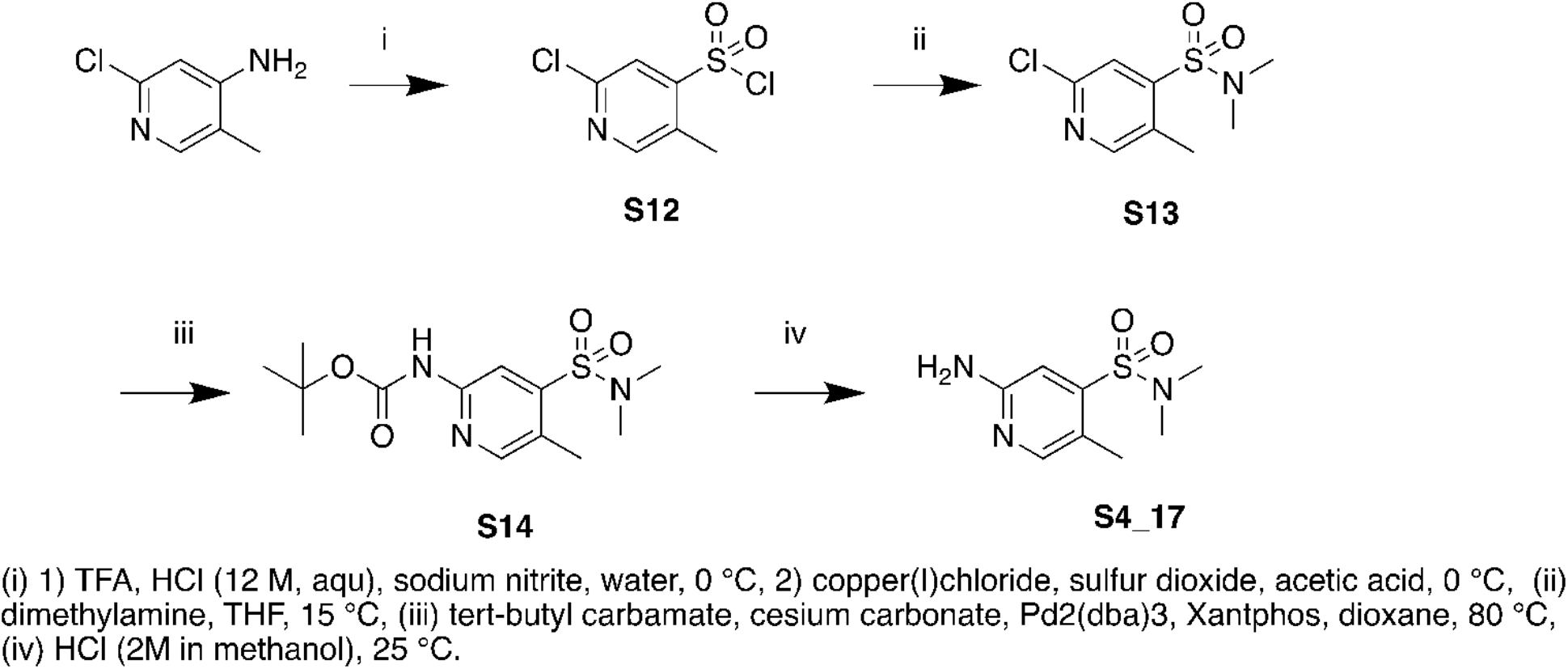
Synthesis of S4_17 for the synthesis of Compound 17

2-chloro-5-methyl-pyridine-4-sulfonyl chloride (S12)

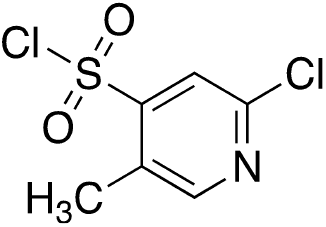

2-chloro-5-methyl-pyridin-4-amine (500 mg, 3.51 mmol), TFA (5 mL), and hydrochloric acid (12M in water, 2.5 mL) was treated with sodium nitrite (723.40 mg, 10.48 mmol) in water (3.5 mL) and stirred for 1 h at 0°C, insoluble materials were removed by filtration. Sulfur dioxide gas was bubbled through a mixture of copper(I)chloride (34.95 mg, 353.00 umol), copper(II)chloride (234.14 mg, 1.74 mmol) in acetic acid (30 mL) for 30 min at 0°C. The mixtures were combined and stirred at 0°C for 1.5 h, diluted with dichloromethane, washed twice with ice-water, twice with saturated aqueous sodium hydrogen carbonate, and saturated aqueous sodium chloride, dried over sodium sulfate, insoluble materials removed by filtration, and volatiles removed under reduced pressure to afford the title compound as a yellow oil (530 mg, 67% yield).

2-chloro-N,N,5-trimethyl-pyridine-4-sulfonamide (S13)

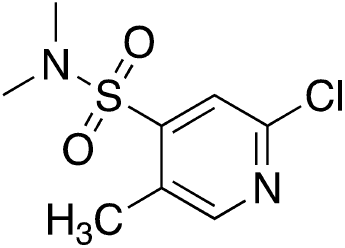

A mixture of 2-chloro-5-methyl-pyridine-4-sulfonyl chloride (S12, 530 mg, 2.34 mmol) and N-methylmethanamine (2M in THF, 5 mL) was stirred for 2 h at 15°C. The mixture was diluted with ethyl acetate, washed with saturated aqueous sodium chloride, dried over sodium sulfate, insoluble materials removed by filtration, volatiles removed under reduced pressure and the residue was purified by column chromatography on silica gel eluting with a gradient of ethyl acetate in petreolum ether to afford the title compound as a white solid (300 mg, 55% yield). 1H NMR (400 MHz, CHLOROFORM-d) δ = 8.41 (s, 1H), 7.69 (s, 1H), 2.90 (s, 6H), 2.58 (s, 3H).

tert-butyl N-[4-(dimethylsulfamoyl)-5-methyl-2-pyridyl]carbamate (S13)

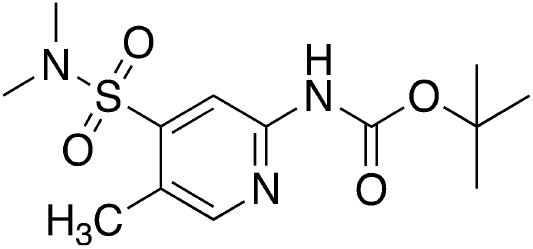

A mixture of 2-chloro-N,N,5-trimethyl-pyridine-4-sulfonamide (S12, 250 mg, 1.07 mmol), tertbutyl carbamate (623.91 mg, 5.33 mmol), cesium carbonate (624.70 mg, 1.92 mmol), Xantphos (73.96 mg, 127.82 umol) and Pd2(dba)3 (39.02 mg, 42.61 umol) in dioxane (5 mL) was stirred at 80°C for 4 hr. The mixture was diluted with water, extracted three times with ethyl acetate, combined organic layers were washed with saturated aqueous sodium chloride, dried over sodium sulfate, insoluble materials were removed by filtration, volatiles removed under reduced pressure and the residue was triturated with petroleum ether:ethyl acetate (1:1) and the title compound isolated by filtration as a yellow solid (260 mg, 77% yield). 1H NMR (400 MHz, DMSO-d6) δ = 10.14 (s, 1H), 8.36 (s, 1H), 8.15 - 8.12 (m, 1H), 2.79 (s, 6H), 2.44 (s, 3H), 1.49 (s, 9H).

2-amino-N,N,5-trimethyl-pyridine-4-sulfonamide (S4_17)

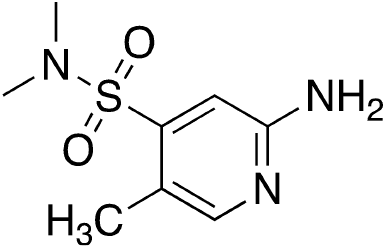

Tert-butyl N-[4-(dimethylsulfamoyl)-5-methyl-2-pyridyl]carbamate (S13, 240 mg, 760.97 umol) in hydrochloric acid (2M in methanol, 5 mL) was stirred at 25°C for 12 hr. Volatiles were removed under reduced pressure to afford the title compound as a yellow solid (300 mg, HCl salt). LRMS (m/z): Calcd [M+H]+ for C_8_H_14_N_3_O_2_S 216.1; found 216.1.

2-(4,5-dichloro-6-oxo-pyridazin-1-yl)-N-[4-(dimethylsulfamoyl)-5-methyl-2-pyridyl]acetamide (17)

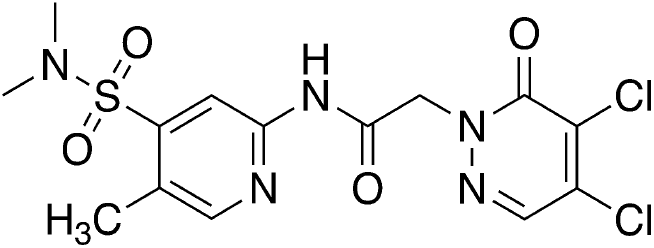

Prepared analogously to Compound 16, using S4_17 and purified by prep-HPLC (column: Phenomenex Synergi C18 150*25*10um; mobile phase: [water (0.225%FA)-ACN];B%: 26%-56%,10min) to afford the title compound as a white solid (45 mg, 38% yield). 1H NMR (400 MHz, DMSO-d6) δ = 11.29 (s, 1H), 8.49 (s, 1H), 8.32 (s, 1H), 8.29 (s, 1H), 5.07 (s, 2H), 2.78 (s, 6H), 2.48 (s, 3H); HRMS (ESI/Q-TOF) RT 2.61 min, m/z: [M + H]+ Calcd for C_14_H_16_Cl_2_N_5_O_4_S 420.0300; Found 420.0295.

**Supplemental Scheme 5.**
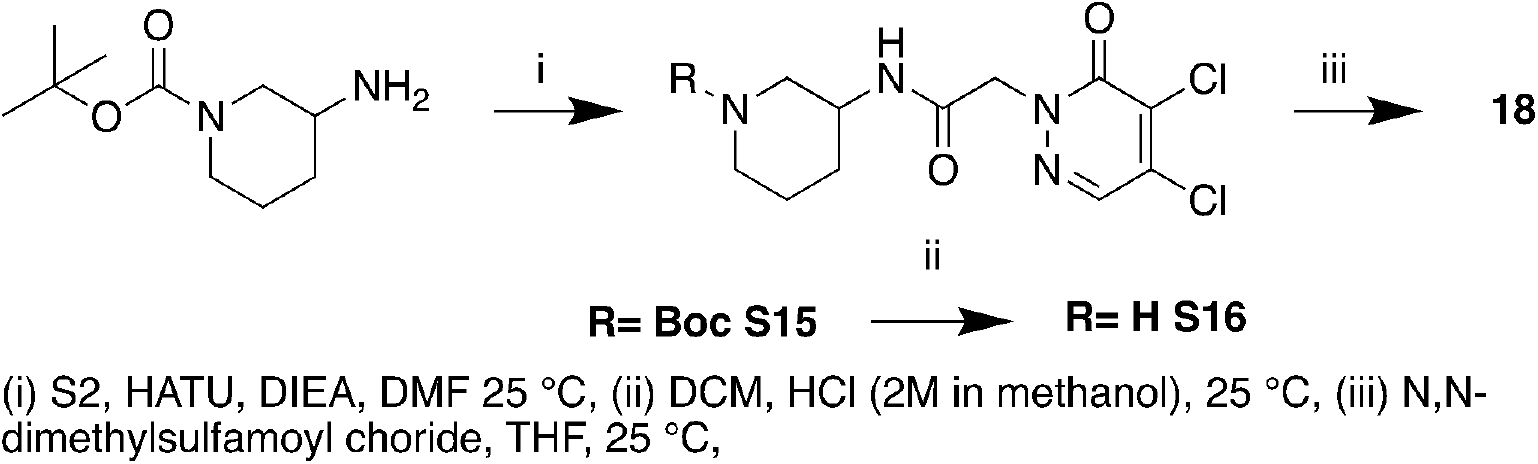
Synthesis of compound 18.

Tert-butyl 3-(2-(4,5-dichloro-6-oxopyridazin-1(6H)-yl)acetamido)piperidine-1-carboxylate (S15)

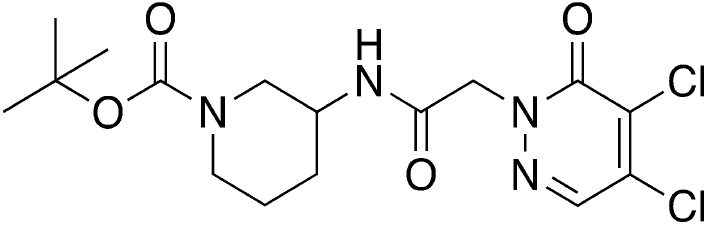

A mixture of tert-butyl 3-aminopiperidine-1-carboxylate (179.61 mg, 896.80 umol), 2-(4,5-dichloro-6-oxopyridazin-1(6H)-yl)acetic acid (S2, 200 mg, 896.80 umol), DIEA (347.72 mg, 2.69 mmol, 468.62 uL) and HATU (511.49 mg, 1.35 mmol) in DMF (4 mL) was stirred at 25 °C for 12 hours. Insoluble materials were removed by filtration, volatiles removed under reduced pressure and the residue was purified by prep-HPLC (column: Phenomenex Luna C18 150 *40mm 10 um; mobile phase: [water (0.225%FA)-ACN]; B%: 28%-58%, 8.5min) to afford the title compound as a white solid (220 mg, 61% yield). LRMS (m/z): [M+H]^+^= 305.1;^1^H NMR (400 MHz, DMSO-d_6_) δ = 8.23 (s, 1H), 8.19 (br d, J = 7.3 Hz, 1H), 4.74 (d, J = 6.4 Hz, 2H), 3.58 (br d, J = 2.9 Hz, 2H), 3.32 (s, 1H), 2.90 (br d, J = 2.8 Hz, 2H), 1.80 (br s, 2H), 1.68 (br d, J = 8.3 Hz, 2H), 1.39 (s, 9H).

2-(4,5-Dichloro-6-oxopyridazin-1(6H)-yl)-N-(piperidin-3-yl)acetamide (S16)

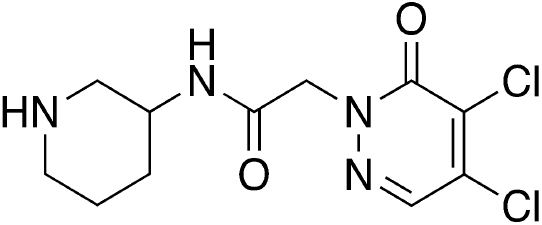

To tert-butyl 3-(2-(4,5-dichloro-6-oxopyridazin-1(6H)-yl)acetamido)piperidine-1-carboxylate (S15, 120 mg, 296.10 umol) in DCM (1 mL) was added hydrochloric acid (4 M in methanol, 4 mL). The mixture was stirred at 25 °C for 1 hour. Volatiles were removed under reduced pressure to afford the title compound as a white solid (100 mg, HCl salt, 98% yield). LRMS (m/z): Calcd [M+H]^+^ for C_11_H_15_Cl_2_N_4_O_2_ 305.1; found 304.8.

2-(4,5-Dichloro-6-oxopyridazin-1(6H)-yl)-N-(1-(N,N-dimethylsulfamoyl)piperidin-3-yl)acetamide (18)

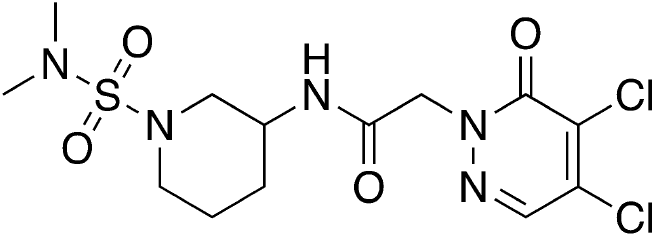

A mixture of 2-(4,5-dichloro-6-oxopyndazin-1(6H)-yl)-N-(piperidin-3-yl)acetamide (S16, 100 mg, 292.72 umol, HCl salt), N,N-dimethylsulfamoyl chloride (63.05 mg, 439.08 umol, 47.05 uL) and TEA (88.86 mg, 878.17 umol, 122.23 uL) in THF (2 mL) was stirred at 25 °C for 2 hours. Volatiles were removed under reduced pressure and the residue was purified by prep-TLC on silica gel, eluting with ethyl acetate to afford the title compound as a white solid (30 mg, 25% yield).^1^H NMR (400 MHz, DMSO-d6) δ = 8.33 (d, J = 7.3 Hz, 1H), 8.31 (s, 1H), 4.81 (d, J = 4.9 Hz, 2H), 3.83 - 3.66 (m, 1H), 3.56 (dd, J = 3.9, 11.8 Hz, 1H), 3.00 - 2.91 (m, 1H), 2.80 (s, 6H), 2.77 - 2.66 (m, 2H), 1.92 - 1.78 (m, 2H), 1.64 - 1.51 (m, 1H), 1.49 - 1.37 (m, 1H); HRMS (ESI/Q-TOF) RT 2.18 min, m/z: [M + H]+ Calcd for C_13_H_20_Cl_2_N_5_O_4_S 412.0613; Found 412.0609.

**Supplemental Scheme 6.**
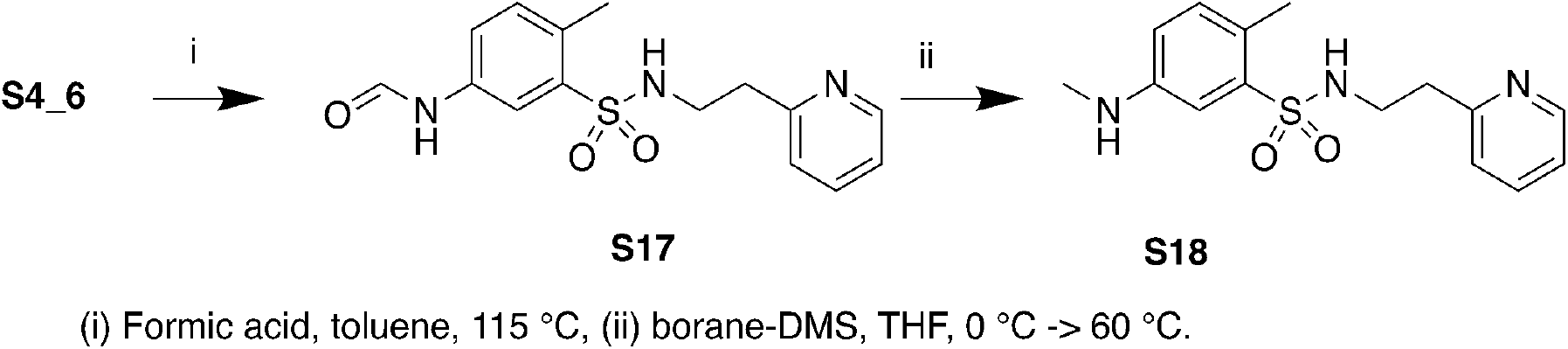
Synthesis of intermediate S18 for the synthesis of compound 19.

N-[4-methyl-3-[2-(2-pyridyl)ethylsulfamoyl]phenyl]formamide (S17)

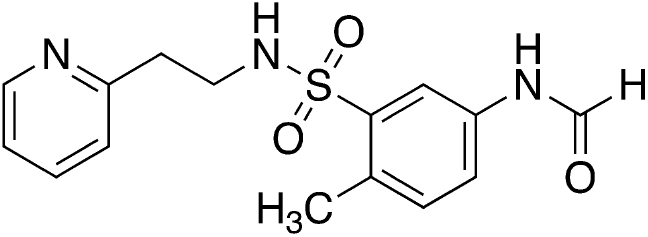

To a mixture of 5-amino-2-methyl-N-[2-(2-pyridyl)ethyl]benzenesulfonamide (S4_6, 1 g, 3.43 mmol, 1 eq) and formic acid (1.90 g, 41.19 mmol, 1.55 mL) in toluene (10 mL) was stirred at 115 C for 2 hr. The mixture was cooled to room temperature and diluted with water, extracted four times with ethyl acetate, combined organics washed repeatedly with saturated aqueous sodium chloride, dried over sodium sulfate, insoluble materials removed by filtration, and volatiles removed under reduced pressure to afford the title compound as a light-yellow oil (1.23 g). ^1^H NMR (400 MHz, DMSO-d_6_) δ = 10.39 (s, 1H), 8.43 - 8.40 (m, 1H), 8.30 (d, J = 1.8 Hz, 1H), 8.12 (d, J = 2.1 Hz, 1H), 7.79 (br t, J = 5.6 Hz, 1H), 7.71 (dd, J = 2.3, 8.1 Hz, 1H), 7.65 (dt, J = 1.8, 7.7 Hz, 1H), 7.30 (d, J = 8.3 Hz, 1H), 7.21 - 7.13 (m, 2H), 3.22 - 3.12 (m, 2H), 2.83 (t, J = 7.4 Hz, 2H), 2.44 (s, 3H).

2-methyl-5-(methylamino)-N-[2-(2-pyridyl)ethyl]benzenesulfonamide (S4_19)

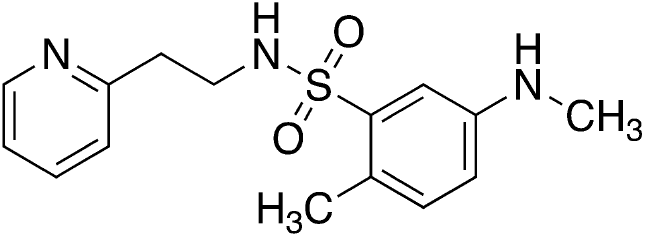

To a mixture of N-[4-methyl-3-[2-(2-pyridyl)ethylsulfamoyl]phenyl]formamide (S17, 1 g, 3.13 mmol) in THF (10 mL) was added borane dimethyl sulfide complex (626.22 uL) drop-wise at 0°C over a period of 15 mins. The mixture was heated to 60°C for 12 hours. The reaction mixture was diluted with water and extracted 4 times with ethyl acetate, combined organic phases were washed three times with saturated aqueous sodium chloride, dried over sodium sulfate, insoluble materials were removed by filtration, volatiles removed under reduced pressure and the residue purified by column chromatography on silica gel eluting with a gradient of ethyl acetate in petroleum ether to afford the title compound as an off-white solid (565 mg, 59% yield). LRMS (m/z): Calcd [M+H]^+^ for C_15_H_19_N_3_O_2_S 306.1; found 306.0; ^1^H NMR (400 MHz, DMSO-d_6_)δ = 8.45 - 8.41 (m, 1H), 8.45 - 8.41 (m, 1H), 7.66 (dt, J = 1.9, 7.7 Hz, 1H), 7.53 (t, J = 5.8 Hz, 1H), 7.21 - 7.13 (m, 2H), 7.07 - 7.03 (m, 2H), 6.62 (dd, J = 2.6, 8.2 Hz, 1H), 5.90 (q, J = 4.9 Hz, 1H), 3.15 - 3.08 (m, 2H), 2.86 - 2.80 (m, 2H), 2.66 (d, J = 5.0 Hz, 3H), 2.33 (s, 3H).

2-(4,5-dichloro-6-oxo-pyridazin-1-yl)-N-methyl-N-[4-methyl-3-[2-(2-pyridyl)ethylsulfamoyl]phenyl]acetamide (19)

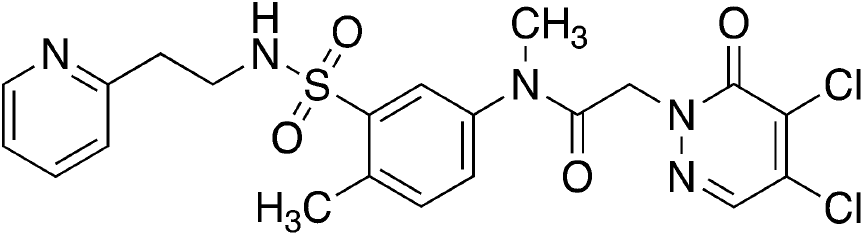

Prepared in an analogous manner to Compound 6 using S4_19.

^1^H NMR (400 MHz, METHANOL-d_4_) δ = 8.27 (br d, J = 4.2 Hz, 1H), 7.95 (s, 1H), 7.89 (br s, 1H), 7.62 (br t, J = 7.0 Hz, 1H), 7.52 (br d, J = 7.2 Hz, 1H), 7.42 (br d, J = 7.8 Hz, 1H), 7.21 - 7.10 (m, 2H), 4.64 (s, 2H), 3.31 - 3.27 (m, 2H), 3.25 (br d, J = 1.7 Hz, 3H), 2.86 (br t, J = 7.0 Hz, 2H), 2.51 (s, 3H); HRMS (ESI/Q-TOF) RT 1.98 min, m/z: [M + H]+ Calcd for C21H22Cl2N5O4S 510.0769; Found 510.0763.

**Supplemental Scheme 7.**
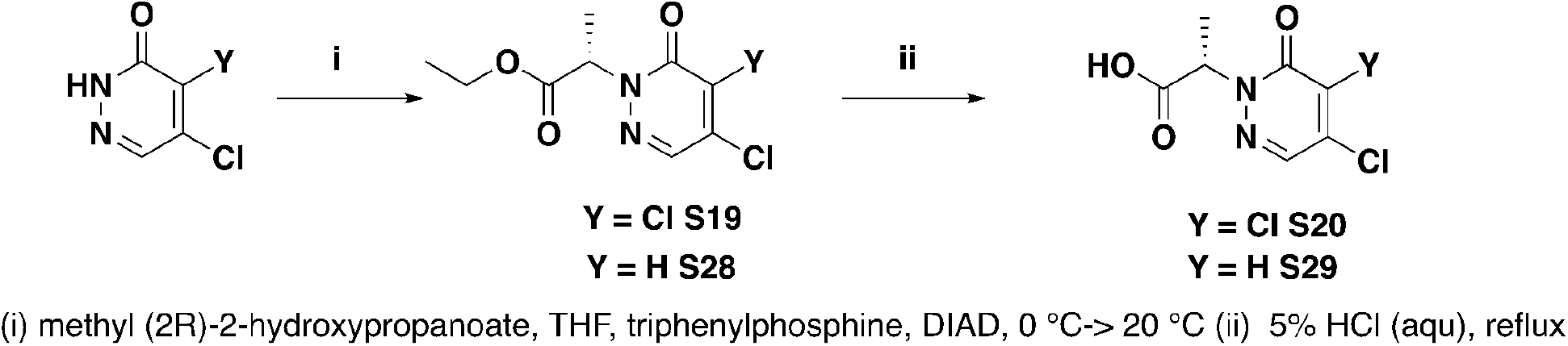
Synthesis of intermediate S20, S29 for the synthesis of Compound 20, 27

methyl (S)-2-(4,5-dichloro-6-oxopyridazin-1(6H)-yl)propanoate (S19)

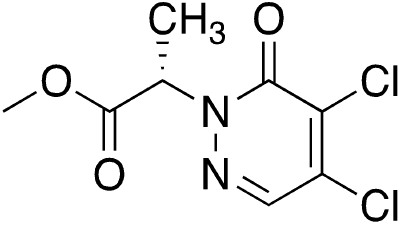

To a solution of 4,5-dichloropyridazin-3(2H)-one (5 g, 30.31 mmol) and methyl (2R)-2-hydroxypropanoate (6.31 g, 60.61 mmol, 5.79 mL) in THF (50 mL) was added triphenylphosphine (15.90 g, 60.61 mmol) and stirred at 20°C for 10 min, cooled to 0°C, and DIAD (12.26 g, 60.61 mmol, 11.79 mL) was added and the reaction mixture was stirred at 20°C for 2h. Volatiles were removed under reduced pressure and the residue was purified by column chromatography on silica gel eluting with a gradient of ethyl acetate in petroleum ether to provide the title compound as a white solid (7.6 g, 99% yield). ^1^H NMR (400 MHz, CHLOROFORM-d) δ = 7.84 (s, 1H), 5.58 (q, J = 7.3 Hz, 1H), 3.76 (s, 3H), 1.70 (d, J = 7.1 Hz, 3H).

(S)-2-(4,5-dichloro-6-oxopyridazin-1(6H)-yl)propanoic acid (S20)

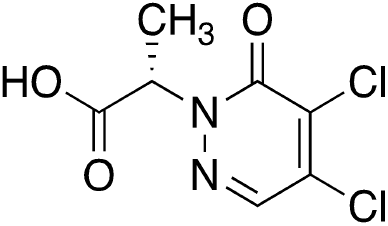

methyl (S)-2-(4,5-dichloro-6-oxopyridazin-1(6H)-yl)propanoate (S19, 7.6 g, 30.27 mmol) was suspended in aqueous hydrogen chloride (5% w/v, 150 mL) and the mixture was stirred at 105°C for 1.5 h. The reaction was cooled to room temperature and the resulting solid was isolated by filtration and washed with water to afford the title compound as a white solid (5.7 g, 79% yield).

(S)-2-(4,5-dichloro-6-oxopyridazin-1(6H)-yl)-N-(4-methyl-3-(N-(2-(pyridin-2-yl)ethyl)sulfamoyl)phenyl)propanamide (20)

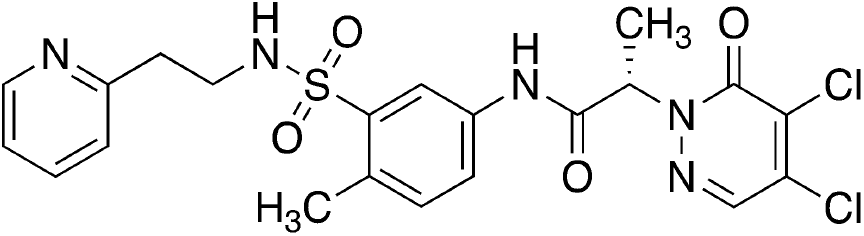

A mixture of (2S)-2-(4,5-dichloro-6-oxo-pyridazin-1-yl)propanoic acid (S20, 100 mg, 421.87 umol) and 5-amino-2-methyl-N-[2-(2-pyridyl)ethyl]benzenesulfonamide (Intermediate 22, 122.92 mg, 421.87 umol), T3P (295.31 mg, 464.06 umol, 275.99 uL, 50% w/w in ethyl acetate) and DIEA (65.43 mg, 506.24 umol, 88.18 uL) in DMF (1 mL) was stirred at 25°C for 3 hr. Volatiles were removed under reduced pressure and the residue was purified by prep-HPLC (column: Phenomenex Synergi C18 150*25*10um;mobile phase: [water(0.225%FA)-ACN];B%: 13%-43%, 10min) to afford the title compound as a white solid (35 mg, 16% yield). ^1^H NMR (400 MHz, DMSO) δ 10.41 (s, 1H), 8.40 (ddd, J = 4.9, 1.9, 1.0 Hz, 1H), 8.32 (s, 1H), 8.07 (d, J = 2.3 Hz, 1H), 7.76 (t, J = 5.8 Hz, 1H), 7.71 (dd, J = 8.3, 2.3 Hz, 1H), 7.64 (td, J = 7.6, 1.9 Hz, 1H), 7.29 (d, J = 8.4 Hz, 1H), 7.17 (ddd, J = 7.6, 4.8, 1.2 Hz, 1H), 7.13 (dt, J = 7.8, 1.1 Hz, 1H), 5.44 (q, J = 7.0 Hz, 1H), 3.14 (td, J = 7.2, 5.7 Hz, 2H), 2.81 (t, J = 7.3 Hz, 2H), 2.43 (s, 3H), 1.61 (d, J = 7.0 Hz, 3H); ^13^C{^1^H} NMR (101 MHz, DMSO) δ 167.93, 158.24, 156.02, 148.97, 138.61, 136.65, 136.40, 136.19, 135.68, 132.87, 132.44, 131.17, 123.21, 122.78, 121.59, 119.52, 59.23, 41.93, 37.28, 19.05, 15.74; SFC (Chiralpac AD-3, 50×4.6 mm I.D., 3uM, Mobile phase: Gradient elution 5%-40% IPA (0.05% DEA) in CO_2_, Flow rate 3 ml/min, PDA detector, Column Temp 35 C, Back pressure 100Bar) RT: 1.029 min, 96.8% ee; HRMS (ESI/Q-TOF) RT: 2.18 min, m/z: [M + H]+ Calcd for C_21_H_22_Cl_2_N_5_O_4_S 510.0769; Found 510.0759.

(R)-2-(4,5-dichloro-6-oxopyridazin-1(6H)-yl)-N-(4-methyl-3-(N-(2-(pyridin-2-yl)ethyl)sulfamoyl)phenyl)propanamide (21)

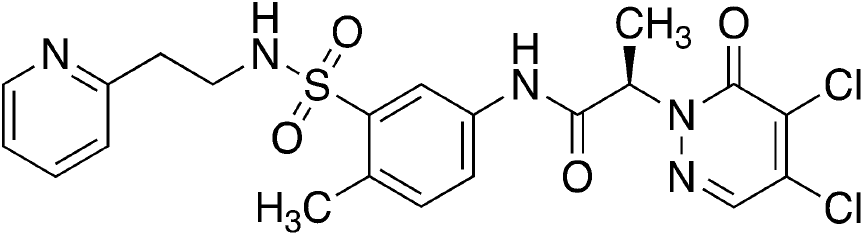

Was prepared analogously to Compound 20.

^1^H NMR (400 MHz, DMSO) δ 10.41 (s, 1H), 8.40 (ddd, J = 4.9, 1.9, 1.0 Hz, 1H), 8.32 (s, 1H), 8.07 (d, J = 2.3 Hz, 1H), 7.76 (t, J = 5.8 Hz, 1H), 7.71 (dd, J = 8.2, 2.3 Hz, 1H), 7.64 (td, J = 7.7, 1.9 Hz, 1H), 7.29 (d, J = 8.4 Hz, 1H), 7.17 (ddd, J = 7.6, 4.8, 1.2 Hz, 1H), 7.13 (dd, J = 7.8, 1.2 Hz, 1H), 5.44 (q, J = 7.0 Hz, 1H), 3.13 (dd, J = 7.6, 5.9 Hz, 2H), 2.81 (t, J = 7.3 Hz, 2H), 2.43 (s, 3H), 1.61 (d, J = 7.1 Hz, 3H); ^13^C{^1^H} NMR (101 MHz, DMSO) δ 167.93, 158.24, 156.02, 148.97, 138.61, 136.65, 136.40, 136.19, 135.68, 132.87, 132.44, 131.17, 123.21, 122.77, 121.59, 119.52, 59.23, 41.93, 37.28, 19.05, 15.74; SFC (Chiralpac AD-3, 50×4.6 mm I.D., 3uM, Mobile phase: Gradient elution 5%-40% IPA (0.05% DEA) in CO2, Flow rate 3 ml/min, PDA detector, Column Temp 35 C, Back pressure 100Bar) RT: 0.642 min, 97.8% ee; HRMS (ESI/Q-TOF) RT: 2.17 min, m/z: [M + H]+ Calcd for C_21_H_22_Cl_2_N_5_O_4_S 510.0769; Found 510.0759.

**Supplemental Scheme 8.**
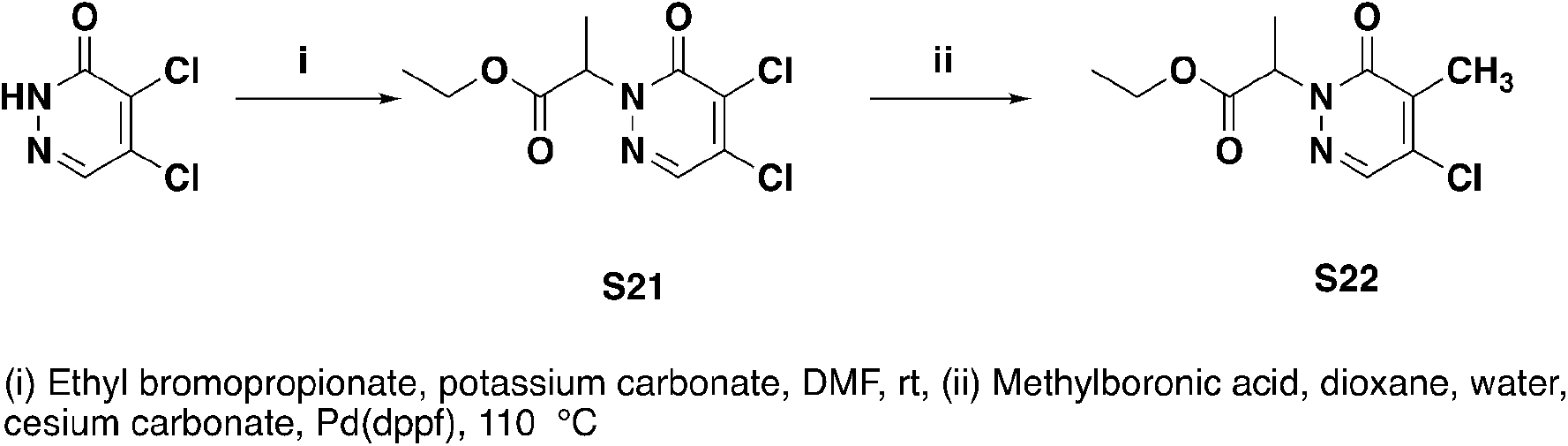
Synthesis of S22 for synthesis of Compound 22

(rac)-ethyl 2-(4,5-dichloro-6-oxopyridazin-1(6H)-yl)propanoate (S21)

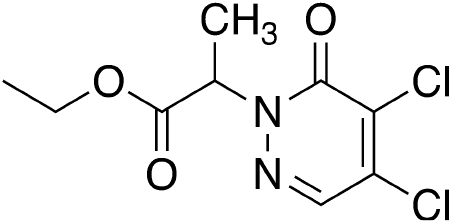

To a solution of 4,5-dichloropyridazin-3(2H)-one (CAS 932-22-9, 1 g, 6.06 mmol) and potassium carbonate (1.68 g, 12.12 mmol) in DMF (10 mL) was added ethyl 2-bromopropanoate (1.21 g, 6.67 mmol, 868.36 uL) at 25 °C, and the mixture was stirred at 50 °C for 2 hours. The mixture was diluted with ethyl acetate and water, layers separated, and the organic phase washed with saturated aqueous sodium chloride, dried over sodium sulfate, insoluble materials removed by filtration, volatiles removed under reduced pressure and the residue purified by column chromatography on silica gel eluting with a gradient of ethyl acetate in petroleum ether, to afford the title compound as a white oil (1.14 g, 71% yield). LRMS (m/z): Calcd [M+H]^+ for^ C_9_H_10_Cl_2_N_2_O_3_ 265.0; found 264.9; ^1^H NMR (400 MHz, CHLOROFORM-d) δ ppm 1.19 (t, J=7.13 Hz, 4 H) 1.62 (d, J=7.25 Hz, 4 H) 1.97 (s, 1 H) 4.15 (q, J=7.09 Hz, 2 H) 5.47 (q, J=7.21 Hz, 1 H) 7.77 (s, 1 H).

ethyl 2-(4-chloro-5-methyl-6-oxo-pyridazin-1-yl)propanoate (S22)

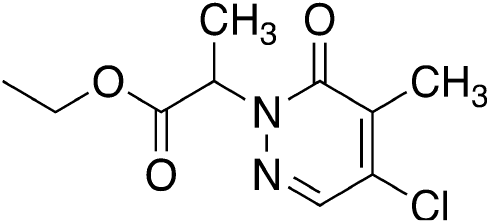

To a mixture of ethyl 2-(4,5-dichloro-6-oxo-pyridazin-1-yl)propanoate (S21, 14.5 g, 54.70 mmol) and methylboronic acid (3.11 g, 51.96 mmol) in dioxane (150 mL) and H2O (15 mL) was added Cs2CO_3_ (53.46 g, 164.09 mmol) and Pd(dppf) (2.00 g, 2.73 mmol). The mixture was stirred at 110°C for 2 hr, diluted with aqueous sodium hydrogen carbonate, extracted twice with ethyl acetate. The combined organic layers were washed with saturated aqueous sodium chloride, dried over sodium sulfate, insoluble materials removed by filtration, volatiles removed under reduced pressure, and the residue purified by column chromatography on silica gel eluting with a gradient of ethyl acetate in petroleum ether to afford the title compound as a white solid (4.12 g, 31% yield). ^1^H NMR (400 MHz, DMSO-d6) δ = 8.10 (s, 1H), 5.40 (q, J = 7.1 Hz, 1H), 4.12 (q, J = 7.0 Hz, 2H), 2.17 (s, 3H), 1.54 (d, J = 7.2 Hz, 3H), 1.16 (t, J = 7.1 Hz, 3H).

2-(4-chloro-5-methyl-6-oxo-pyridazin-1-yl)-N-[4-methyl-3-[2-(2-pyridyl)ethylsulfamoyl]phenyl]propanamide (22)

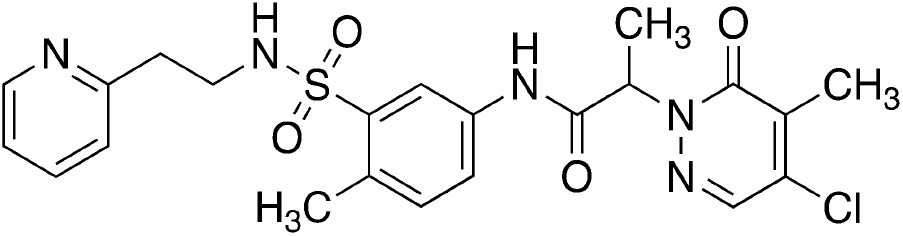

Prepared in an analogous manner to Compound 6 ^1^H NMR (400 MHz, DMSO-d_6_) δ = 10.41 (s, 1H), 8.40 (dd, J = 0.9, 4.8 Hz, 1H), 8.13 - 8.07 (m, 2H), 7.78 - 7.69 (m, 2H), 7.64 (dt, J = 1.8, 7.6 Hz, 1H), 7.29 (d, J = 8.4 Hz, 1H), 7.20 - 7.11 (m, 2H), 5.39 (q, J = 7.1 Hz, 1H), 3.18 - 3.10 (m, 2H), 2.81 (t, J = 7.3 Hz, 2H), 2.43 (s, 3H), 2.17 (s, 3H), 1.59 (d, J = 7.1 Hz, 3H); HRMS (ESI/Q-TOF) RT: 2.14 min, m/z: [M + H]+ Calcd for C_22_H_25_ClN_5_O_4_S 490.1315; Found 490.1310.

**Supplemental Scheme 9.**
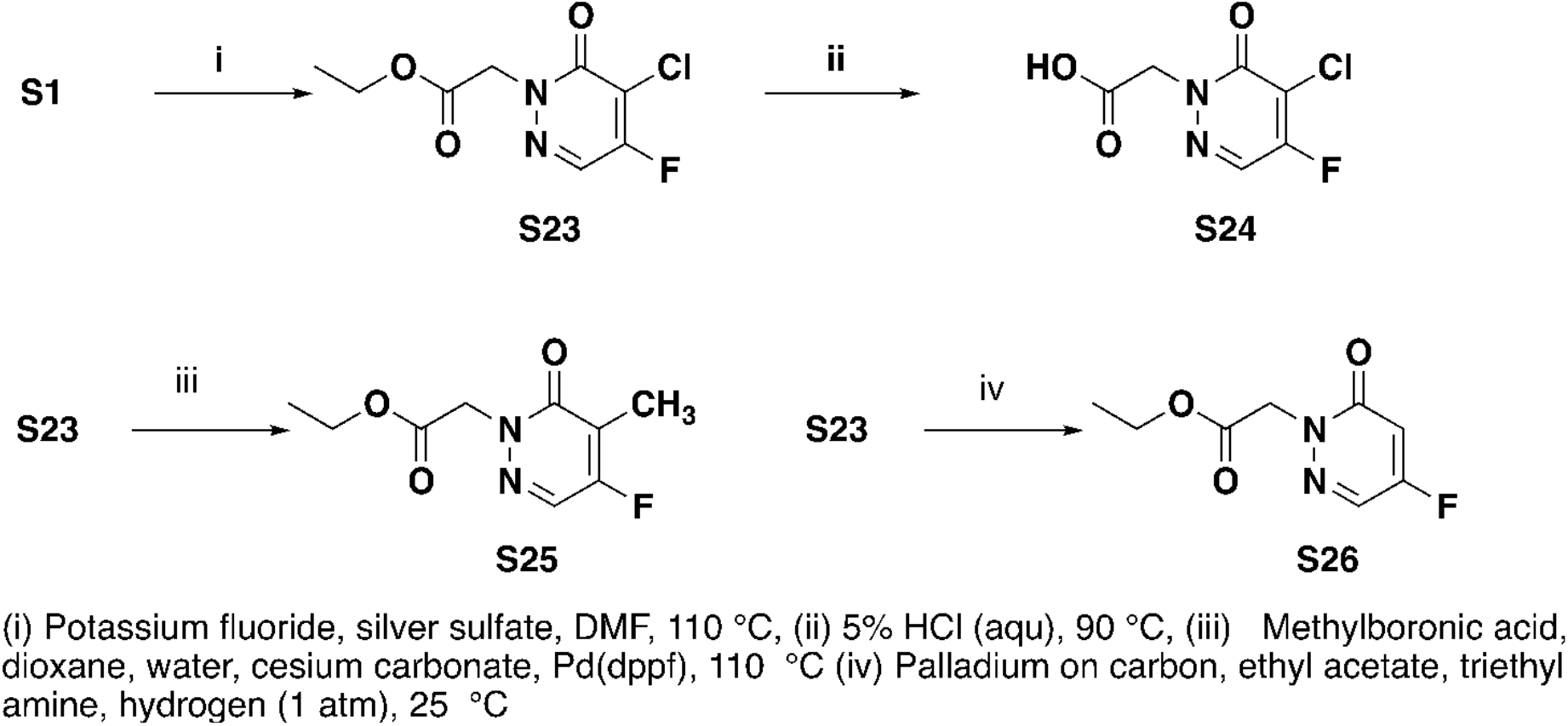
Synthesis of S24, S25, S26 for synthesis of compounds 23, 24, 25

ethyl 2-(5-chloro-4-fluoro-6-oxopyridazin-1(6H)-yl)acetate (S23)

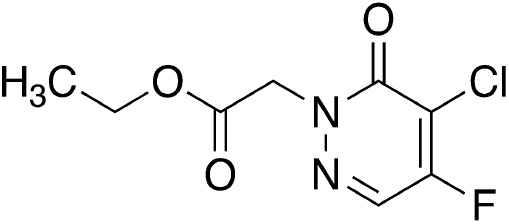

A mixture of ethyl 2-(4,5-dichloro-6-oxopyridazin-1(6H)-yl)acetate (S1, 1 g, 3.98 mmol), potassium fluoride (1.16 g, 19.92 mmol, 466.53 uL) and silver sulfate (124.19 mg, 398.30 umol, 67.49 uL) in DMF (10 mL) was stirred at 110 °C for 12 hours. The mixture was diluted with water and extracted twice with ethyl acetate, combined organic layers were washed with saturated aqueous sodium chloride, dried over sodium sulfate, insoluble materials removed by filtration, volatiles removed under reduced pressure and the residue purified by column chromatography on silica gel eluting with a gradient of ethyl acetate in petroleum ether to afford the title compound as a white solid (470 mg, 50% yield). LRMS (m/z): Calcd [M+H]^+^ for C_8_H_8_ClFN_2_O_3_ 235.0; found 235.0.

2-(5-chloro-4-fluoro-6-oxopyridazin-1(6H)-yl)acetic acid (S24)

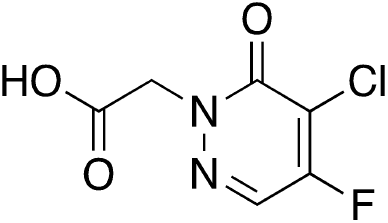

A mixture of ethyl 2-(5-chloro-4-fluoro-6-oxopyridazin-1(6H)-yl)acetate (S23, 100 mg, 426.24 umol) in hydrochloric acid (3 M in water, 10 mL) was stirred at 90 °C for 1 hour. The mixture was frozen and volatiles removed by lyophilization to afford the title compound as a white solid (100 mg), which was used without further manipulation. LRMS (m/z): Calcd [M+H]^+ for^ C_6_H_4_ClFN_2_O_3_ 207.0; found 207.1.

2-(5-chloro-4-fluoro-6-oxo-pyridazin-1-yl)-N-[4-methyl-3-[2-(2-pyridyl)ethylsulfamoyl]phenyl]acetamide (23)

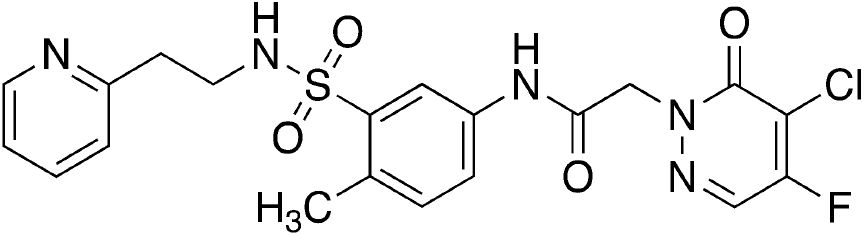

A mixture of 5-amino-2-methyl-N-[2-(2-pyridyl)ethyl]benzenesulfonamide (S4_6, 100 mg, 343.21 umol), 2-(5-chloro-4-fluoro-6-oxopyridazin-1(6H)-yl)acetic acid (S24, 85.07 mg, 411.85 umol), 2-chloro-1-methyl-pyridin-1-ium iodide (131.53 mg, 514.81 umol) and DIEA (88.71 mg, 686.42 umol, 119.56 uL) in THF (2 mL) was stirred at 25 °C for 2 hours. Volatiles were removed under reduced pressure and the residue was purified by prep-HPLC (column: Shim-pack C18 150*25*10um; mobile phase: [water (0.225%FA)-ACN]; B%: 5%-38%, 11min) to afford the title compound as a white solid (85 mg, 52% yield). ^1^H NMR (400 MHz, DMSO-d_6_) δ = 10.59 (s, 1H), 8.45 - 8.39 (m, 2H), 8.10 (d, J = 2.2 Hz, 1H), 7.78 (t, J = 5.7 Hz, 1H), 7.71 - 7.62 (m, 2H), 7.31 (d, J = 8.4 Hz, 1H), 7.18 (dd, J = 5.3, 7.0 Hz, 1H), 7.14 (d, J = 7.8 Hz, 1H), 5.00 (s, 2H), 3.18 - 3.13 (m, 2H), 2.82 (t, J = 7.3 Hz, 2H), 2.44 (s, 3H);^19^F NMR (376 MHz, DMSO-d_6_) δ −111.08; HRMS (ESI/Q-TOF) RT: 1.75 min, m/z: [M + H]+ Calcd for C_20_H_20_ClFN_5_O_4_S 480.0908; Found 480.0901.

ethyl 2-(4-fluoro-5-methyl-6-oxopyridazin-1(6H)-yl)acetate (S25)

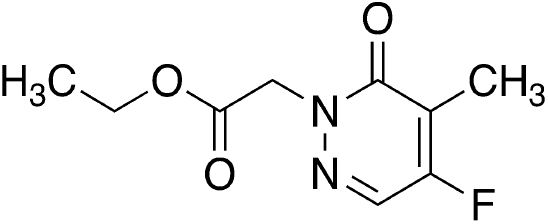

A mixture of ethyl 2-(5-chloro-4-fluoro-6-oxopyridazin-1(6H)-yl)acetate (S23, 1 g, 4.26 mmol), methylboronic acid (280.66 mg, 4.69 mmol), cesium carbonate (4.17 g, 12.79 mmol) and Pd(dppf)2 (155.94 mg, 213.12 umol) in dioxane (20 mL) and water (2 mL) was stirred at 110 °C for 6 hours. The mixture was cooled to room temperature, diluted with water, extracted twice with ethyl acetate, combined organic phases were washed with saturated aqueous sodium chloride, dried over sodium sulfate, insoluble materials removed by filtration, volatiles removed under reduced pressure and the residue was purified by column chromatography on silica gel eluting with a gradient of ethyl acetate in petroleum ether to provide the title compound as a pink solid (560 mg, 61% yield). LRMS (m/z): Calcd [M+H]^+^ for C_9_H_11_FN_2_O_3_ 215.1; found 215.0; ^1^H NMR (400 MHz, CHLOROFORM-d) δ = 7.79 (s, 1H), 4.89 (s, 2H), 4.29 - 4.23 (m, 2H), 2.15 (d, J = 2.6 Hz, 3H), 1.35 - 1.29 (m, 3H).

2-(4-fluoro-5-methyl-6-oxopyridazin-1(6h)-yl)-n-(4-methyl-3-(n-(2-(pyridin-2-yl)ethyl)sulfamoyl)phenyl)acetamide (24)

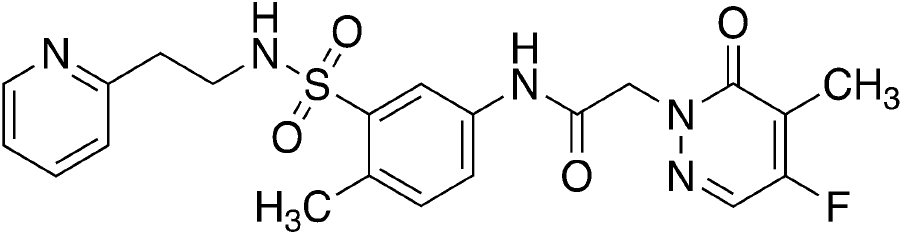

Was prepared analogously to Compound 23.

^1^H NMR (400 MHz, DMSO-d6) δ = 10.57 (s, 1H), 8.47 - 8.38 (m, 1H), 8.21 (s, 1H), 8.11 (d, J = 2.1 Hz, 1H), 7.78 (t, J = 5.7 Hz, 1H), 7.72 - 7.62 (m, 2H), 7.31 (d, J = 8.4 Hz, 1H), 7.19 (dd, J = 5.3, 7.0 Hz, 1H), 7.14 (d, J = 7.7 Hz, 1H), 4.92 (s, 2H), 3.19 - 3.11 (m, 2H), 2.82 (t, J = 7.3 Hz, 2H), 2.44 (s, 3H), 2.02 (d, J = 2.3 Hz, 3H); HRMS (ESI/Q-TOF) RT: 1.68 min, m/z: [M + H]+ Calcd for C_21_H_23_FN_5_O_4_S 460.1455; Found 460.1448.

ethyl 2-(4-fluoro-6-oxo-pyridazin-1-yl)acetate (S26)

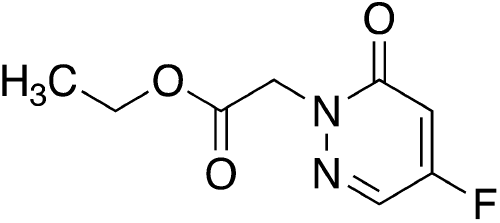

A mixture of ethyl 2-(5-chloro-4-fluoro-6-oxopyridazin-1(6H)-yl)acetate (S23, 300 mg, 1.21 mmol, 1 eq), palladium on carbon (60 mg, 5% w/w) and TEA (366.28 mg, 3.62 mmol, 503.82 uL) in EtOAc (6 mL) was stirred under an atmosphere of hydrogen at 25 °C for 12 hr. Insoluble materials were removed by filtration through Celite, volatiles removed under reduced pressure, and the residue purified by column chromatography on silica gel eluting with a gradient of ethyl acetate in petroleum ether to afford the title compound as a colorless oil (214 mg, 88% yield). LRMS (m/z): Calcd [M+H]^+^ for C_8_H_9_FN_2_O_3_ 201.1; found 201.0.

2-(4-fluoro-6-oxo-pyridazin-1-yl)-N-[4-methyl-3-[2-(2-pyridyl)ethylsulfamoyl]phenyl]acetamide (25)

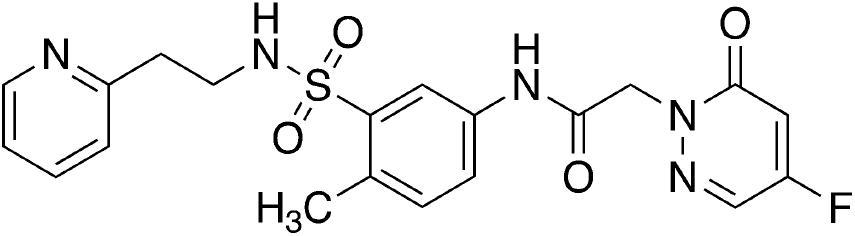

Was prepared analogously to Compound 23.

1H NMR (400 MHz, DMSO-d6) δ = 10.56 (s, 1H), 8.41 – 8.40 (d, 1H), 8.26 (d, 1H), 8.10 (d, 1H), 7.77 (m, 1H), 7.69 (m, 1H), 7.64 (m, 1H), 7.31 −7.29 (m, 1H), 7.14 – 7.12 (m, 1H), 7.02 – 6.99 (m, 1H), 4.92 (s, 2H), 3.18 - 3.13 (m, 2H), 2.84 – 2.80 (m, 2H), 2.44 (s, 3H); HRMS (ESI/Q-TOF) RT: 1.47 min, m/z: [M + H]+ Calcd for C_20_H_21_FN_5_O_4_S 446.1298; Found 446.1290.

**Supplemental Scheme 10.**
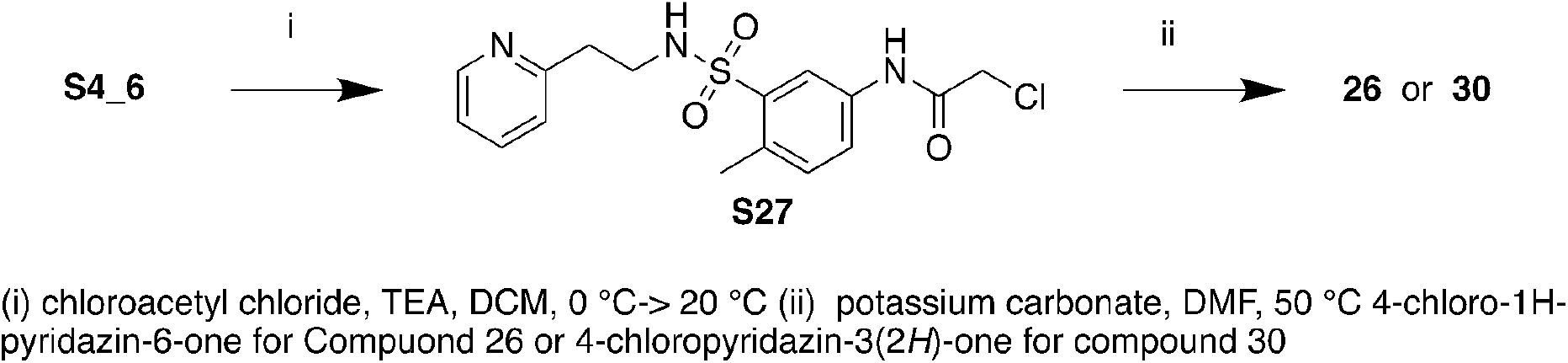
Synthesis of Compounds 26 and 30

2-chloro-N-[4-methyl-3-[2-(2-pyridyl)ethylsulfamoyl]phenyl]acetamide (S27)

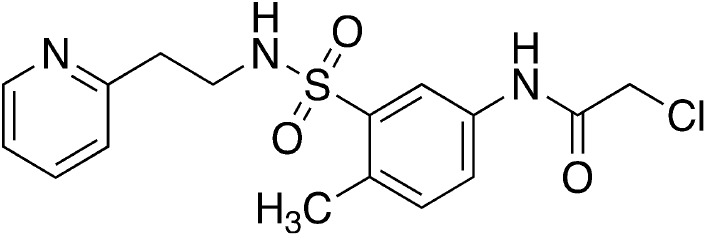

A mixture of 5-amino-2-methyl-N-[2-(2-pyridyl)emyl]benzenesulfonamide (S4_6, 300 mg, 1.03 mmol) and TEA (208.38 mg, 2.06 mmol, 286.62 uL) in DCM (5 mL) was treated with 2-chloroacetyl chloride (151.18 mg, 1.34 mmol, 106.46 uL) at 0°C. The mixture was stirred at 20°C for 12h. The mixture was diluted with water and extracted twice with ethyl acetate. Combined organic layers were washed with saturated aqueous sodium chloride, dried over sodium sulfate, insoluble materials removed by filtration, volatiles removed under reduced pressure and the residue purified by prep-TLC on silica gel, eluting with dichloromethane:methanol 10:1, to give the title compound as a yellow solid (200 mg, 53% yield). LRMS (m/z): Calcd [M+H]^+^ for C_16_H_18_ClN_3_O_3_S 368.1; found 368.1; ^1^H NMR (400 MHz, DMSO-d_6_) δ = 10.53 (s, 1H), 8.45 - 8.37 (m, 1H), 8.09 (d, J = 2.3 Hz, 1H), 7.82 - 7.72 (m, 2H), 7.65 (dt, J = 1.8, 7.6 Hz, 1H), 7.32 (d, J = 8.4 Hz, 1H), 7.21 - 7.12 (m, 2H), 4.26 (s, 2H), 3.20 - 3.14 (m, 2H), 2.83 (t, J = 7.4 Hz, 2H), 2.45 (s, 3H).

2-(4-chloro-6-oxopyridazin-1(6H)-yl)-N-(4-methyl-3-(N-(2-(pyridin-2-yl)ethyl)sulfamoyl)phenyl)acetamide (26)

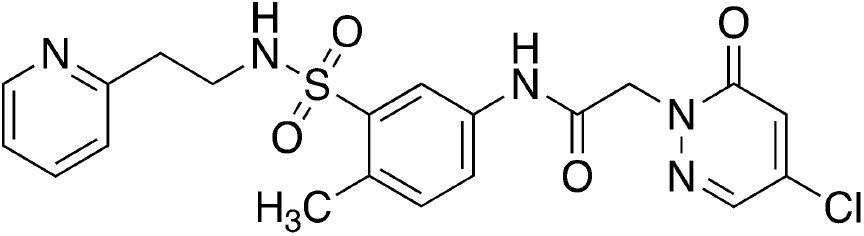

A mixture of 4-chloro-1H-pyridazin-6-one (35.51 mg, 272.07 umol), 2-chloro-N-[4-methyl-3-[2-(2-pyridyl)ethylsulfamoyl]phenyl]acetamide (S27, 100 mg, 247.33 umol, HCl salt) and potassium carbonate (102.55 mg, 742.00 umol) in DMF (2 mL) was stirred at 50°C for 2 hr. The mixture was diluted with water, extracted twice with ethyl acetate, combined organic phases were washed with saturated aqueous sodium chloride, dried over sodium sulfate, insoluble materials were removed by filtration, volatiles removed under reduced pressure and the residue was purified by prep-HPLC (column: Phenomenex Synergi C18 150*25*10um;mobile phase: [water(0.225%FA)-ACN];B%: 2%-32%,10min) to give the title compound as a yellow solid (45 mg, 36% yield). ^1^H NMR (400 MHz, DMSO) δ 10.56 (s, 1H), 8.40 (ddd, J = 4.9, 1.9, 0.9 Hz, 1H), 8.14 (d, J = 2.4 Hz, 1H), 8.09 (d, J = 2.3 Hz, 1H), 7.77 (t, J = 5.8 Hz, 1H), 7.69 (dd, J = 8.2, 2.3 Hz, 1H), 7.64 (td, J = 7.7, 1.9 Hz, 1H), 7.35 (d, J = 2.4 Hz, 1H), 7.30 (d, J = 8.3 Hz, 1H), 7.17 (ddd, J = 7.6, 4.9, 1.2 Hz, 1H), 7.13 (dt, J = 7.9, 1.1 Hz, 1H), 4.90 (s, 2H), 3.14 (td, J = 7.2, 5.6 Hz, 2H), 2.81 (t, J = 7.3 Hz, 2H), 2.43 (s, 3H); ^13^C{^1^H} NMR (101 MHz, DMSO) δ 164.96, 158.53, 158.24, 148.97, 139.12, 138.70, 136.54, 136.51, 136.41, 132.97, 131.10, 127.07, 123.20, 122.47, 121.59, 119.22, 54.62, 41.96, 37.28, 19.06; HRMS (ESI/Q-TOF) RT: 1.68 min, m/z: [M + H]+ Calcd for C_20_H_21_ClN_5_O_4_S 462.1002; Found 462.0994.

methyl (S)-2-(4-chloro-6-oxopyridazin-1(6H)-yl)propanoate (S28)

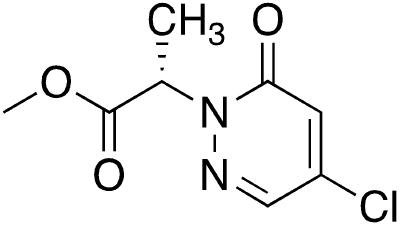

Prepared analogously to S19.

^1^H NMR (400 MHz, DMSO-d_6_) δ = 8.15 (d, J = 2.4 Hz, 1H), 7.33 (d, J = 2.4 Hz, 1H), 5.43 (q, J = 7.2 Hz, 1H), 3.64 (s, 3H), 1.53 (d, J = 7.1 Hz, 3H).

(2S)-2-(4-chloro-6-oxo-pyridazin-1-yl)propanoic acid (S29)

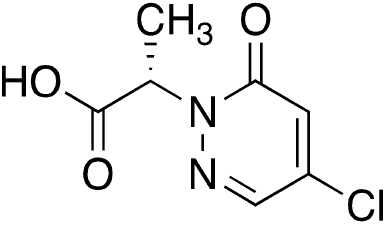

Prepared analogously to S20.

^1^H NMR (400 MHz, DMSO-d6) δ = 8.14 (d, J = 2.3 Hz, 1H), 7.31 (d, J = 2.3 Hz, 1H), 5.34 (q, J = 7.2 Hz, 1H), 1.52 (d, J = 7.2 Hz, 3H).

(2S)-2-(4-chloro-6-oxo-pyridazin-1-yl)-N-[4-methyl-3-[2-(2-pyridyl)ethylsulfamoyl]phenyl]propanamide (27) (BRD0639)

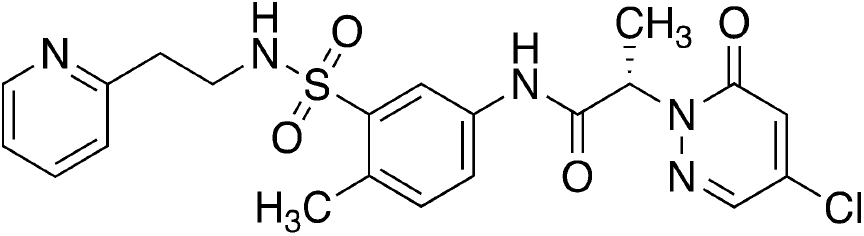

Prepared analogously to Compound 20.

^1^H NMR (400 MHz, DMSO) δ 10.42 (s, 1H), 8.43 – 8.38 (m, 1H), 8.17 (d, J = 2.4 Hz, 1H), 8.08 (d, J = 2.3 Hz, 1H), 7.75 (s, 1H), 7.71 (dd, J = 8.2, 2.3 Hz, 1H), 7.64 (td, J = 7.7, 1.9 Hz, 1H), 7.32 (d, J = 2.4 Hz, 1H), 7.29 (d, J = 8.4 Hz, 1H), 7.17 (ddd, J = 7.7, 4.9, 1.2 Hz, 1H), 7.13 (d, J = 7.7 Hz, 1H), 5.39 (q, J = 7.1 Hz, 1H), 3.14 (t, J = 7.3 Hz, 2H), 2.81 (t, J = 7.3 Hz, 2H), 2.43 (s, 3H), 1.60 (d, J = 7.1 Hz, 3H).;^13^C{^1^H} NMR (101 MHz, DMSO) δ 168.39, 158.64, 158.25, 148.97, 138.88, 138.61, 136.75, 136.40, 136.04, 132.85, 131.05, 126.60, 123.21, 122.67, 121.59, 119.43, 57.99, 41.94, 37.29, 19.05, 15.69; SFC (Chiralpac AS-3, 50×4.6 mm I.D., 3uM, Mobile phase: Gradient elution 5%-40% Methanol (0.05% DEA) in CO_2_, Flow rate 3 ml/min, PDA detector, Column Temp 35 C, Back pressure 100Bar) RT 2.037 min, 98.7% ee; HRMS (ESI/Q-TOF) RT: 1.88 min, m/z: [M + H]+ Calcd for C_21_H_23_ClN_5_O_4_S 476.1159; Found 476.1152.

(2R)-2-(4-chloro-6-oxo-pyridazin-1-yl)-N-[4-methyl-3-[2-(2-pyridyl)ethylsulfamoyl]phenyl]propanamide (28)

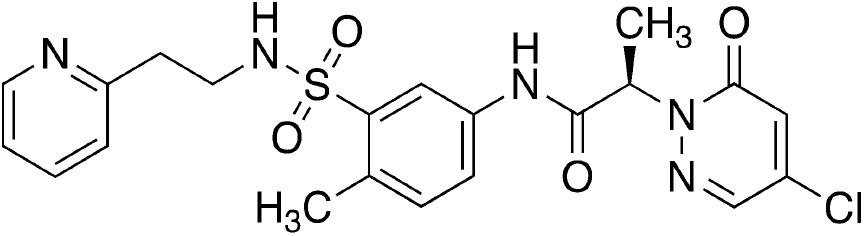

Was prepared analogously to Compound 20.

^1^H NMR (400 MHz, DMSO) δ 10.42 (s, 1H), 8.40 (ddd, J = 4.9, 1.9, 0.9 Hz, 1H), 8.17 (d, J = 2.4 Hz, 1H), 8.08 (d, J = 2.3 Hz, 1H), 7.75 (s, 1H), 7.71 (dd, J = 8.2, 2.3 Hz, 1H), 7.64 (td, J = 7.6, 1.9 Hz, 1H), 7.32 (d, J = 2.4 Hz, 1H), 7.29 (d, J = 8.4 Hz, 1H), 7.17 (ddd, J = 7.6, 4.9, 1.2 Hz, 1H), 7.13 (dt, J = 7.8, 1.1 Hz, 1H), 5.39 (q, J = 7.1 Hz, 1H), 3.14 (t, J = 7.3 Hz, 2H), 2.81 (t, J = 7.3 Hz, 2H), 2.43 (s, 3H), 1.60 (d, J = 7.1 Hz, 3H); ^13^C{^1^H} NMR (101 MHz, DMSO) δ 168.39, 158.64, 158.25, 148.97, 138.88, 138.61, 136.75, 136.40, 136.04, 132.85, 131.05, 126.60, 123.21, 122.67, 121.59, 119.43, 57.99, 41.94, 37.29, 19.05, 15.70; SFC (Chiralpac AS-3, 50×4.6 mm I.D., 3uM, Mobile phase: Gradient elution 5%-40% Methanol (0.05% DEA) in CO_2_, Flow rate 3 ml/min, PDA detector, Column Temp 35 C, Back pressure 100Bar) RT 1.700 min, 95.7% ee; HRMS (ESI/Q-TOF) RT: 1.89 min, m/z: [M + H]+ Calcd for C_21_H_23_ClN_5_O_4_S 476.1159; Found 476.1152.

(2S)-2-(4-chloro-6-oxo-pyridazin-1-yl)-N-methyl-N-[4-methyl-3-[2-(2-pyridyl)ethylsulfamoyl]phenyl]propanamide (29)

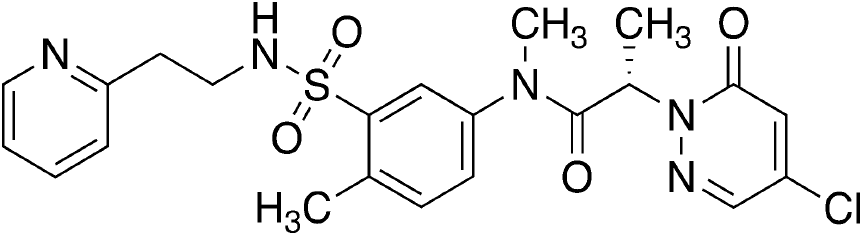

Was prepared analogously to Compound 20.

^1^H NMR (400 MHz, DMSO-d_6_) δ = 8.33 (br d, J = 4.8 Hz, 1H), 7.96 (d, J = 2.0 Hz, 1H), 7.88 (d, J = 1.6 Hz, 1H), 7.68 (dt, J = 1.9, 7.7 Hz, 1H), 7.52 (dd, J = 1.6, 8.0 Hz, 1H), 7.40 (d, J = 8.0 Hz, 1H), 7.24 - 7.17 (m, 2H), 6.86 (s, 1H), 5.26 (q, J = 6.8 Hz, 1H), 3.30 (br s, 2H), 3.26 (s, 3H), 2.91 (dt, J = 1.8, 7.2 Hz, 2H), 2.55 (s, 3H), 1.43 (d, J = 7.0 Hz, 3H); SFC (Chiralpac AD-3, 50×4.6 mm I.D., 3uM, Mobile phase: Gradient elution 5%-40% IPA (0.05% DEA) in CO_2_, Flow rate 3 ml/min, PDA detector, Column Temp 35 C, Back pressure 100Bar) RT: 2.072 min, 86.3% ee (minor isomer at 1.895 min); HRMS (ESI/Q-TOF) RT: 1.83 min, m/z: [M + H]+ Calcd for C_22_H_25_ClN_5_O_4_S 490.1315; Found 490.1308.

2-(5-chloro-6-oxo-pyridazin-1-yl)-N-[4-methyl-3-[2-(2-pyridyl)ethylsulfamoyl]phenyl]acetamide (30)

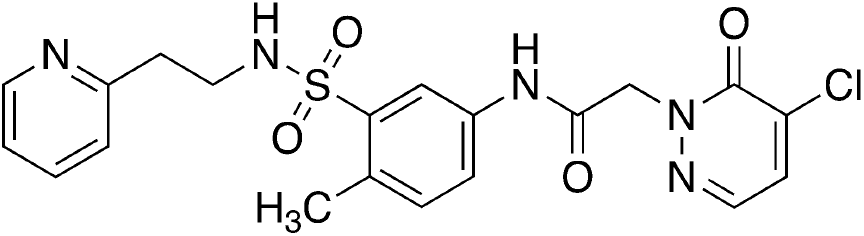

Prepared analogously to Compound 26.

^1^H NMR (400 MHz, DMSO) δ 10.59 (s, 1H), 8.41 (ddd, J = 4.8, 1.9, 0.9 Hz, 1H), 8.10 (d, J = 2.4 Hz, 1H), 7.95 (d, J = 4.5 Hz, 1H), 7.86 (d, J = 4.5 Hz, 1H), 7.77 (t, J = 5.8 Hz, 1H), 7.69 (dd, J = 8.2, 2.3 Hz, 1H), 7.64 (td, J = 7.6, 1.8 Hz, 1H), 7.30 (d, J = 8.4 Hz, 1H), 7.17 (ddd, J = 7.6, 4.9, 1.2 Hz, 1H), 7.13 (dt, J = 7.8, 1.1 Hz, 1H), 4.97 (s, 2H), 3.15 (td, J = 7.3, 5.7 Hz, 2H), 2.81 (t, J = 7.3 Hz, 2H), 2.43 (s, 3H);^13^C{^1^H} NMR (101 MHz, DMSO) δ 164.87, 158.24, 156.91, 148.97, 138.71, 136.54, 136.40, 136.07, 135.17, 132.98, 131.11, 130.90, 123.20, 122.47, 121.58, 119.21, 55.85, 41.97, 37.28, 19.07; HRMS (ESI/Q-TOF) RT: 1.52 min, m/z: [M + H]+ Calcd for C_20_H_21_ClN_5_O_4_S 462.1002; Found 462.0997.

**Supplemental Figure 3.**
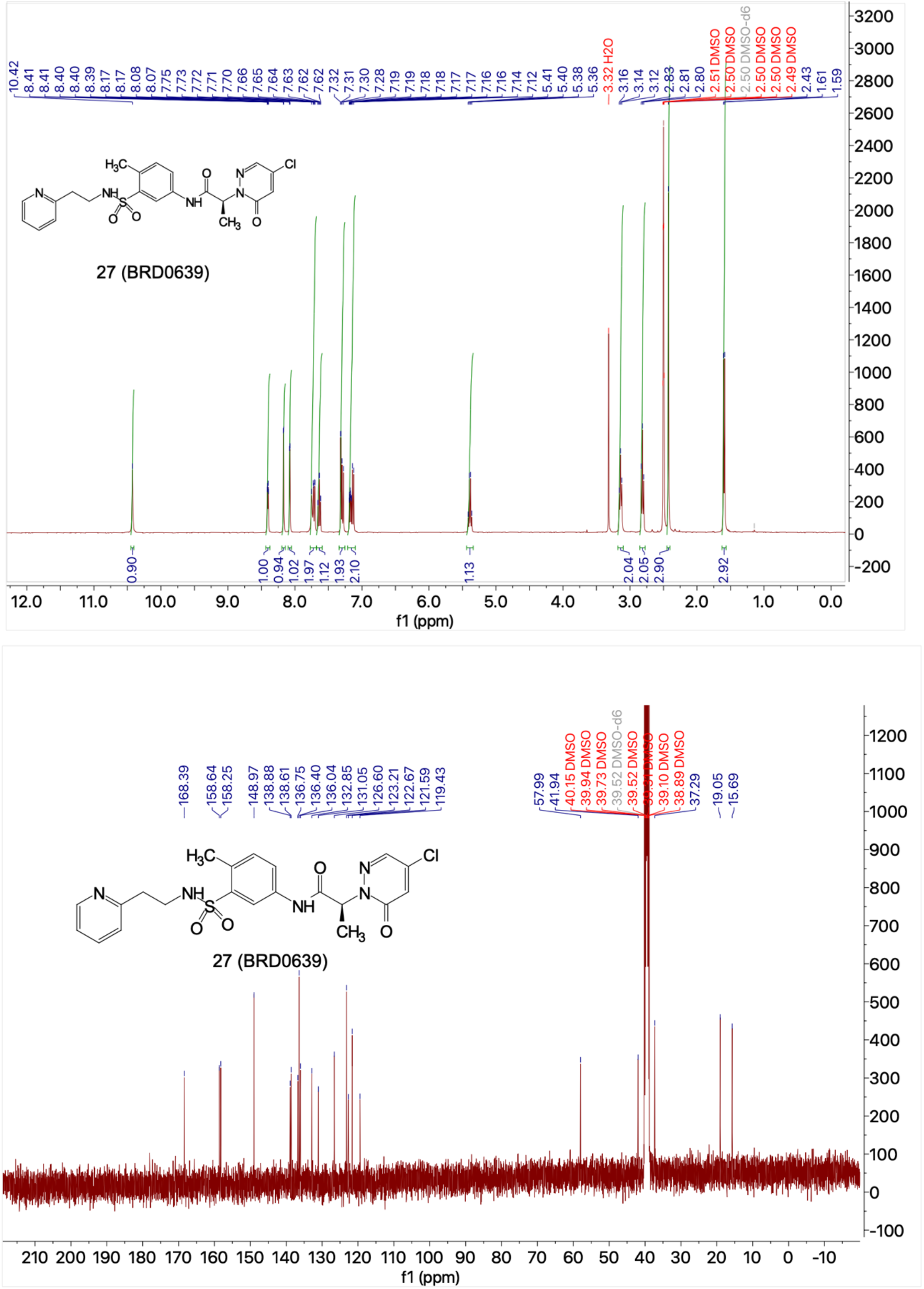
1H and 13C NMR spectra of BRD0639 (27)

**Supplemental Figure 4.**
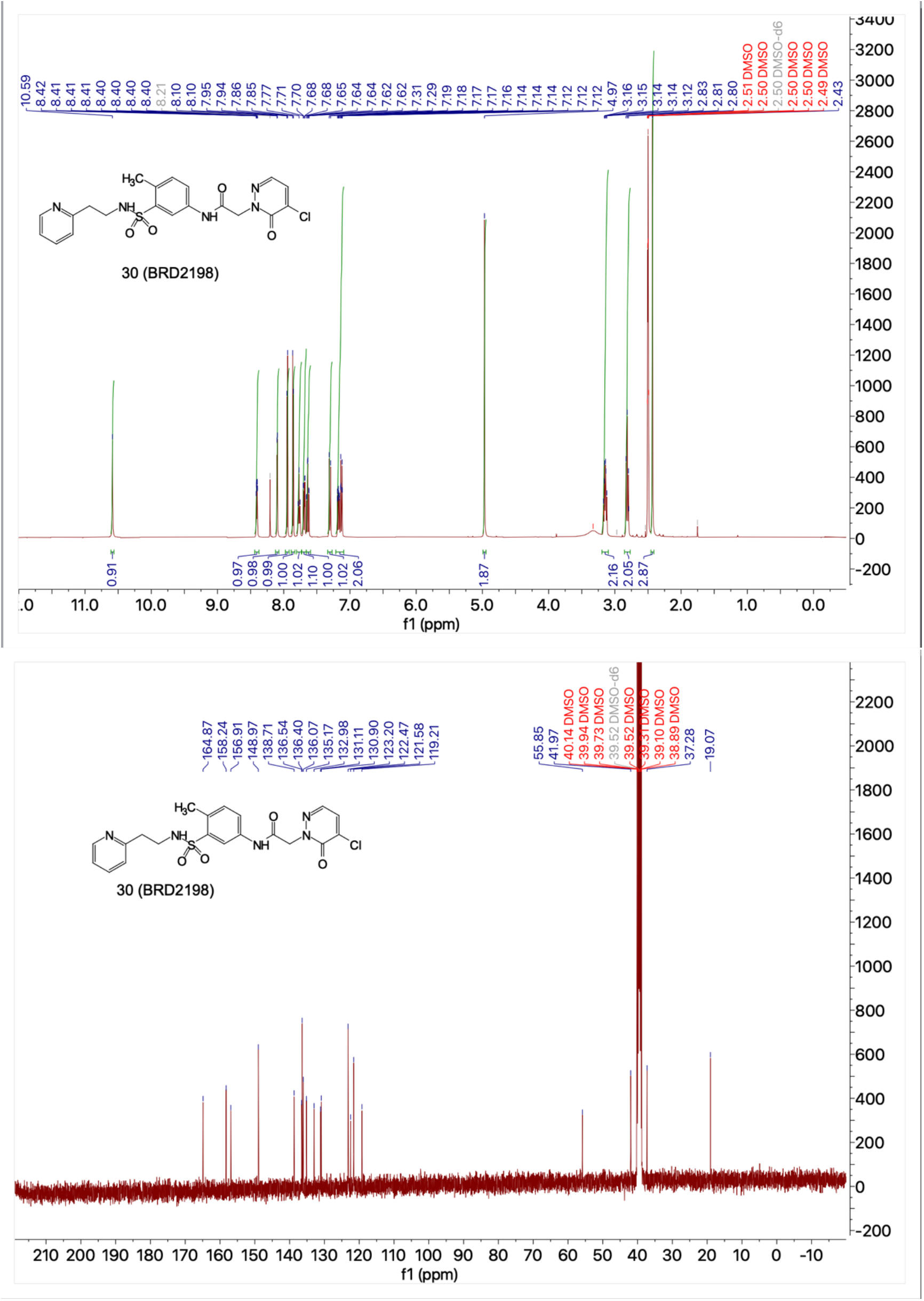
1H and 13C NMR spectra of BRD2198 (30)

